# Increased chromatin accessibility mediated by nuclear factor I drives transition to androgen receptor splice variant dependence in castration-resistant prostate cancer

**DOI:** 10.1101/2024.01.10.575110

**Authors:** Larysa Poluben, Mannan Nouri, Jiaqian Liang, Andreas Varkaris, Betul Ersoy-Fazlioglu, Olga Voznesensky, Irene I. Lee, Xintao Qiu, Laura Cato, Ji-Heui Seo, Matthew L. Freedman, Adam G. Sowalsky, Nathan A. Lack, Eva Corey, Peter S. Nelson, Myles Brown, Henry W. Long, Joshua W. Russo, Steven P. Balk

**Author notes:** Address correspondence to Steven P. Balk; or Joshua W. Russo. co-first authors (Larysa Poluben, Mannan Nouri).

## Abstract

Androgen receptor (AR) splice variants, of which ARv7 is the most common, are increased in prostate cancer (PC) that develops resistance to androgen signaling inhibitor drugs, but the extent to which these variants drive AR activity, and whether they have novel functions or dependencies, remain to be determined. We generated a subline of VCaP PC cells (VCaP16) that is resistant to the AR inhibitor enzalutamide (ENZ) and found that AR activity was independent of the full-length AR (ARfl), despite its continued high-level expression, and was instead driven by ARv7. The ARv7 cistrome and transcriptome in VCaP16 cells mirrored that of the ARfl in VCaP cells, although ARv7 chromatin binding was weaker, and strong ARv7 binding sites correlated with higher affinity ARfl binding sites across multiple models and clinical samples. Notably, although ARv7 expression in VCaP cells increased rapidly in response to ENZ, there was a long lag before it gained chromatin binding and transcriptional activity. This lag was associated with an increase in chromatin accessibility, with the AR and nuclear factor I (NFI) motifs being most enriched at these more accessible sites. Moreover, the transcriptional effects of combined NFIB and NFIX knockdown versus ARv7 knockdown were highly correlated. These findings indicate that ARv7 can drive the AR program, but that its activity is dependent on adaptations that increase chromatin accessibility to enhance its intrinsically weak chromatin binding.

## INTRODUCTION

Prostate cancer (PC) that recurs after standard androgen deprivation therapy (ADT) with a GnRH agonist or antagonist (castration-resistant prostate cancer, CRPC) is in most cases still dependent on androgen receptor (AR) activity driven by residual androgens ^1–3^. AR activity in CRPC tumors can be further suppressed by agents such as abiraterone, which decreases adrenal androgen synthesis, or use of more effective AR antagonists, such as enzalutamide (ENZ), apalutamide, and darolutamide ^4–6^. Unfortunately, patients treated with these androgen-signaling inhibitor (ASI) drugs invariably progress. Therapeutic options for these patients are then taxanes, or PARP inhibitors in men with *BRCA2* loss, but responses to these agents are not durable. A subset of tumors that progress despite ADT combined with ASI drugs become AR independent, with loss of AR expression in some cases, or reduced AR expression and loss of AR transcriptional activity in other cases ^7^. A subset of these AR-independent tumors express neuroendocrine markers and have undergone varying degrees of neuroendocrine differentiation, and is referred to as neuroendocrine PC ^8^. Others that are AR negative and do not express neuroendocrine markers have been termed double-negative PC ^7^. However, the majority of tumors that become resistant to ADT and ASI drugs continue to express high levels of AR and exhibit AR transcriptional activity.

The high levels of AR expression in CRPC are in part driven by amplification of the *AR* gene and an upstream enhancer, which occurs in the majority of cases ^9–11^, and can enhance responses to low levels of androgen. A smaller subset of these tumors has mutations in the AR-ligand binding domain (LBD) that allow alternative steroids or antagonists to act as agonists which may compromise the efficacy of available ASIs ^12–14^. Moreover, one particular mutation in the LBD allows ENZ to act as an agonist, although this is rarely found in clinical samples ^15, 16^. Multiple studies also suggest that activation of several signaling pathways may directly or indirectly enhance AR activity at low androgen levels or in the presence of antagonists ^17^. Finally, increasing evidence indicates that expression of AR-splice variants that are constitutively active due to a lack of the LBD, of which AR variant-7 (ARv7) is the most common, contributes to persistent AR activity in tumors that progress on ASI therapy ^18–23^.

ARv7 is generated by splicing to a cryptic exon downstream of exon 3 (encoding the DNA binding domain), resulting in an AR that is truncated after the DNA binding domain. ARv7 expression is low or undetectable in primary PC prior to ADT, but is expressed in most cases of CRPC and is generally further increased in tumors that become resistant to ASI drugs ^21, 22^. ARv7 can drive AR activity in cells where ARfl activity is suppressed ^24–27^. Moreover, expression of ARv7 in circulating tumor cells in men with CRPC is associated with resistance to treatment with ASI drugs, suggesting it is a mediator of resistance ^19, 23, 28^. However, most tumors expressing ARv7 also express high levels of ARfl, so the extent to which ARv7 is an ARfl-independent driver of AR activity or a biomarker of tumors that are more aggressive and resistant due to other mechanisms, including high levels of ARfl, remains unclear. To address the role and biological importance of ARv7 we have generated and characterized an ENZ-resistant subline of VCaP PC cells that expresses both high levels of ARfl and increased levels of ARv7. We find that ARv7 is the major driver of AR activity and is independent of ARfl, but that it is dependent on adaptations that enhance chromatin accessibility for its activity.

## RESULTS

### ENZ-resistance in VCaP cells is associated with AR reactivation and expression of ARv7

VCaP cells were cultured in medium with increasing concentrations of ENZ (up to 16 μM) over ∼2 months to select for ENZ resistance, alongside passage matched vehicle controls. Parental VCaP cells (grown in medium with 10% FBS) with VCaP cells exposed short term (4 days) to 16 μM ENZ (added to medium with 10% FBS) and to the VCaP cells adapted to 16 μM ENZ (VCaP16) were then compared. Addition of ENZ initially suppressed VCaP cell proliferation, but the proliferative rate of the ENZ-adapted VCaP16 cells (maintained on 16 μM ENZ) was comparable to that of the parental VCaP cells (**Figure 1A**). As expected, short-term ENZ treatment of VCaP cells (VCaP-E) initially suppressed the expression of several AR-regulated genes (*KLK3*, *KLK2*, *FKBP5*, and *NKX3.1*) (**Figure 1B**). However, expression of these genes was then partially restored or increased in the VCaP16 cells.

**Figure 1.**
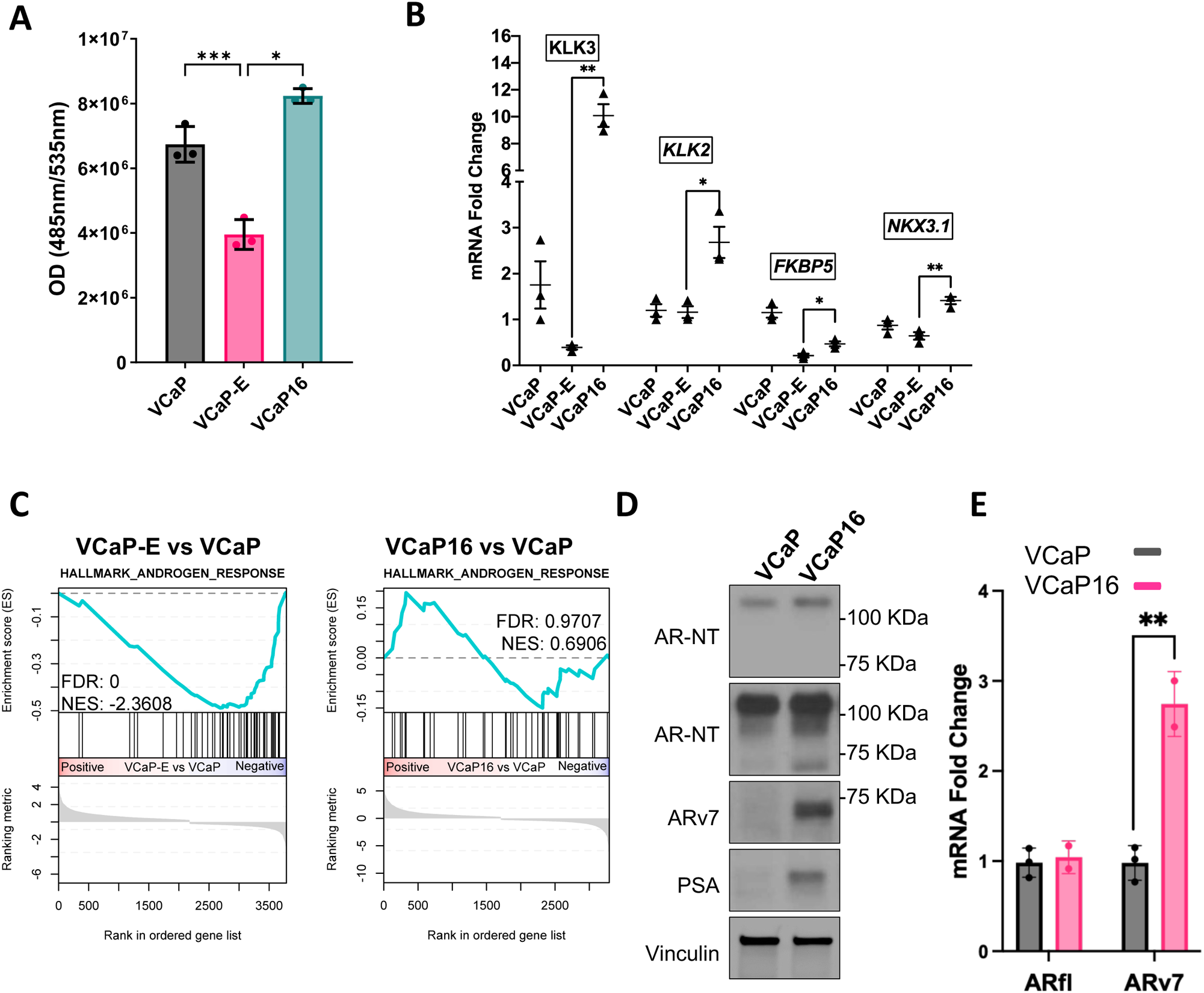
VCaP16 cells exhibit increased ARv7 expression and resistance to AR blockade. **A)** Cell proliferation assay for VCaP, VCaP treated with 16 μM ENZ (VCaP-E), and VCaP16 (maintained in 16 μM ENZ) at 7 days post-treatment (*p<0.05, **p<0.01, ANOVA). **B)** Expression of AR-regulated genes *KLK3*, *KLK2*, *FKBP5*, *NKX3.1* assessed by qRT-PCR in VCaP16, VCaP cells treated with ENZ for 4 days (VCaP-E), and parental VCaP cells (* p<0.05, **p<0.01, Mann-Whitney U). **C)** Enrichment plots for Hallmark Androgen Response gene set by GSEA comparing RNA-seq for VCaP cells treated with ENZ for 4 days (VCaP-E) or VCaP16 versus parental VCaP. Significantly differentially expressed genes (adjusted p-value < 0.05) were used as an input. **D)** Immunoblot of protein from VCaP16 and parental VCaP cells. **E)** Expression of ARfl and ARv7 assessed by qRT-PCR in VCaP (black) and VCaP16 (red) cells (*p<0.05, **p<0.01, Unpaired t-test, two-tailed).

For a more comprehensive assessment we carried out RNA-seq on acutely ENZ-treated VCaP and VCaP16 and compared them to parental VCaP. Consistent with the above results, VCaP cells acutely exposed to ENZ for 4 days had marked decreases in Hallmark gene sets related to Androgen Response (NES −2.4) (**Figure 1C**, left) and to proliferation, including G2M Checkpoint (NES −3.54) and E2F Targets (NES −2.90) (not shown), when compared to parental VCaP (**Figure S1A, B**). Interestingly, the MTORC1 gene set was also markedly decreased (NES −3.02) following acute ENZ treatment, consistent with previous data showing that androgen stimulates this pathway ^29^. Notably, no Hallmark gene sets were significantly increased in the acutely ENZ treated cells.

Gene sets related to proliferation and MTORC1 were also decreased in VCaP16 relative to parental VCaP, but to a lesser degree (NES −2.16 for G2M Checkpoint) than in acutely ENZ-treated VCaP cells (**Figure S1A, C**). Moreover, unlike VCaP cells treated initially with ENZ, VCaP16 exhibited no significant changes in the Androgen Response gene set compared to VCaP cells, indicating that AR activity was substantially restored (**Figure 1C, right**). This restoration was not associated with a substantial increase in expression of ARfl protein or mRNA (**Figure 1D, E**). Moreover, DNA sequencing did not show the AR LBD mutation reported to enhance ENZ agonist activity or other AR exonic mutations ^15, 16^ (**Figure S1D**). In contrast, the restored AR-signaling activity in VCaP16 was associated with an increase in the ARv7 protein and mRNA, as well as a substantial increase in PSA/KLK3 mRNA and protein expression (**Figure 1B, D, E**). Overall, these findings show that AR activity in the VCaP16 cells was substantially restored, and this was associated with an increase in ARv7.

### AR activity in VCaP16 cells is driven by ARv7

We next used siRNA to determine the contribution of ARfl versus ARv7 to the AR activity in the VCaP16 cells. Remarkably, depletion of ARfl with an siRNA against exon 7 (siEx7) had no clear effect on the AR regulated *KLK3* gene (**Figure 2A**). In contrast, selective depletion of ARv7 (siV7) markedly decreased *KLK3* expression, which was also decreased by an siRNA in exon 1 targeting both ARfl and ARv7 (siEx1). We then carried out RNA-seq to compare gene expression in VCaP16 cells treated with control versus ARv7 siRNA, which confirmed that the ARv7 siRNA markedly decreased expression of Hallmark Androgen Response genes (**Figure 2B, S1A**). Conversely, RNA-seq also confirmed that selective downregulation of ARfl with the siEx7 did not significantly decrease expression of genes in this gene set (**Figure 2C, S1A**). Notably, principal component analysis similarly showed that the ARfl knockdown with siEx7 had minimal effect on overall gene expression, while the ARv7 and Ex1 siRNA samples were separated from the control siRNA and clustered together (**Figure 2D**).

**Figure 2.**
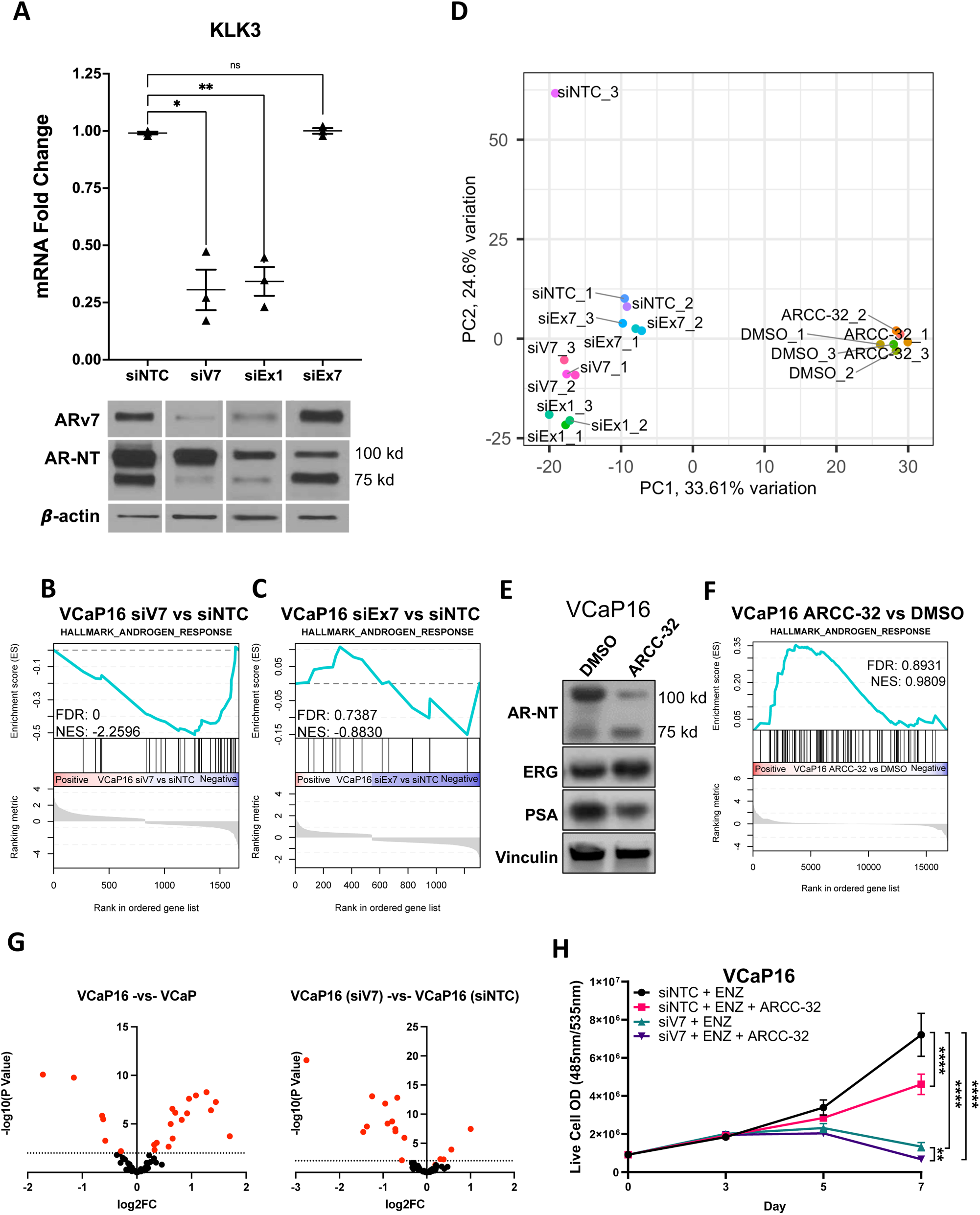
AR activity in VCaP16 cells is dependent on ARv7. **A)** *Upper, KLK3* gene expression by qRT-PCR and *Lower*, immunoblot for ARfl and ARv7 in VCaP16 cells after siRNA knockdown with non-targeting control (siNTC), ARv7 (siV7), both ARv7 and ARfl (siEx1), or just Arfl (siEx7) siRNA. **B, C)** Enrichment for Hallmark Androgen Response gene set in RNA isolated from VCaP16 cells with siRNA knockdown of ARv7 (siV7) or ARfl (siEx7). Significantly differentially expressed genes (adjusted p-value < 0.05) were used as an input. **D)** Principal component analysis of RNA-seq data from VCaP16 cells with siRNA knockdown described in *A),* with addition of VCaP16 cells in DMSO (DMSO) and VCaP16 cells treated with ARCC-32 (500 nM) for 36 hrs. **E)** Immunoblot of protein from VCaP16 in DMSO and VCaP16 cells treated with ARCC-32 (500 nM) for 36 hrs. **F)** Enrichment for Hallmark Androgen Response gene set in RNA-seq data from VCaP16 cells in DMSO and VCaP16 cells treated with ARCC-32 (500 nM) for 36 hrs. All differentially expressed genes were used as an input. **G) Differential** gene expression of the 59 genes that make up the ARv7 gene set from Sharp et al. comparing VCaP16 versus VCaP cells (left) or VCaP16 treated with ARv7 siRNA versus non-target control siRNA (siNTC) (right). **H)** Live cell DNA fluorescence in VCaP16 cells treated with non-target control siRNA (siNTC) or ARv7 siRNA (siV7) +/− 500 nM ARCC-32.

To further assess the contribution of ARfl to AR activity, we treated the VCaP16 cells with an AR degrader (AR PROTAC®, ARCC-32, ARD) targeted to the LBD to selectively decrease ARfl protein (**Figure 2E**). For this experiment we used a relatively high concentration of ARCC-32 (500 nM) to effectively compete with ENZ, which was kept in the medium to avoid transient stimulation of ARfl by androgen in the FBS medium. Notably, the AR degrader decreased expression of PSA, possibly reflecting more potent suppression of ARfl than obtained with the siEx7 siRNA. Despite the decrease in PSA protein, RNA-seq showed that overall AR activity was not decreased by the AR degrader, with a trend toward increased activity **(Figure 2F**). Principal component analysis similarly showed that the AR degrader had minimal overall impact on gene expression as compared to the vehicle (DMSO) control (**Figure 2D**).

To assess the extent to which ARv7 activity in VCaP16 is reflective of clinical samples, we examined a panel of genes whose expression was correlated with ARv7 protein expression in clinical CRPC ^21^. Notably, expression of this gene set was increased in VCaP16 compared to VCaP, and was decreased by siRNA targeting ARv7, consistent with ARv7 functioning similarly in VCaP16 and in CRPC (**Figure 2G**). Finally, we assessed the effects of depleting ARfl versus ARv7 on proliferation. Growth of VCaP16 cells was suppressed by the AR degrader targeting ARfl, but it was more markedly suppressed by ARv7 siRNA (**Figure 2H).** Consistent with this ARv7 dependence, Gene Set Enrichment Analysis (GSEA) showed that siARv7-treated VCaP16 cells had marked decreases in E2F targets, G2M Checkpoint, and MTORC1 signaling gene sets, in addition to the decrease in the Androgen Response gene set (see **Figure S1A**). In contrast, these gene sets were not decreased in the siEx7 or AR-degrader treated cells. Together these findings indicated that AR activity in the ENZ-adapted VCaP16 cells was being driven primarily by ARv7.

### ARv7 has limited chromatin binding capacity acutely following ENZ treatment

We reported previously that ARv7 increases rapidly in response to AR inhibition (in absolute levels and relative to ARfl) ^30^. Consistent with our previous report, ARv7 was increased in VCaP cells within 2 days of ENZ treatment, and levels peaked by 7-14 days (**Figure 3A**). However, this increase in ARv7 alone was not sufficient to drive AR-transcriptional activity, as it took several more weeks in culture before the cells became ENZ-resistant with restoration of AR activity. As noted above, this did not reflect selection for cells with ENZ-responsive AR mutations. To determine more broadly whether this reflected selection for a subclone we compared VCaP versus VCaP16 cells by whole exome sequencing (WES), which showed no significant differences in copy number or driving mutations (**Figure S2**).

**Figure 3.**
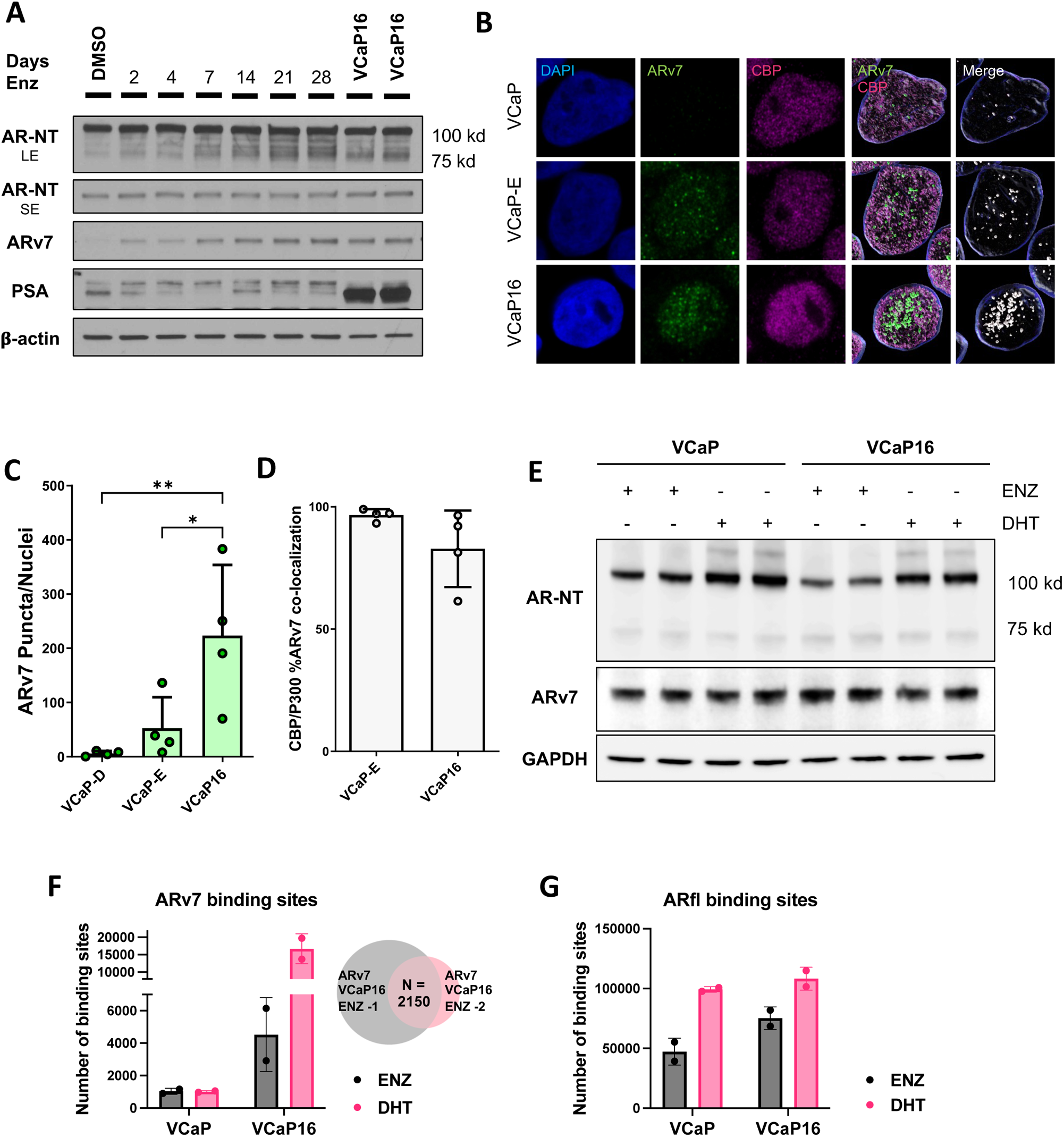
ARv7 induced acutely by ENZ is nuclear but lacks chromatin binding capacity. **A)** Immunoblot of protein isolated from VCaP cells treated with 16 ⍰M ENZ from 2-28 days and protein isolated from VCaP16 cells. AR-NT, antibody targeting AR N-terminus (long and short exposures). **B)** Confocal immunofluorescence imaging of ARv7 and CBP/p300 in DMSO or 16 μM ENZ (4 days) treated VCaP cells versus VCaP16 cells (maintained in 16 μM ENZ). **C, D)** Quantification of z-stack 3-dimensional images produced from panel showing number of ARv7 puncta per nuclei in four fields (C) and fraction of ARv7 puncta that also contain CBP/p300 (D). **E)** Immunoblot of chromatin fraction isolated from VCaP and VCaP16 cells following formalin crosslinking for ChIP. VCaP cells were grown in 10% FBS supplemented with 16 ⍰M ENZ for 72 hrs and media was then replaced with fresh 10% FBS medium supplemented with 16 μM ENZ or 10 nM DHT for 4 hrs prior to crosslinking. VCaP16 cells were in maintenance media (10% FBS, 16⍰M ENZ) and then media was replaced with fresh 10% FBS medium supplemented with 16 ⍰M ENZ or 10 nM DHT for 4 hrs prior to crosslinking. **F)** ARv7 binding sites based on ChIP-seq in samples from (E) and Venn diagram showing overlap between ARv7 sites in VCaP16 cells (ENZ) in biological replicates. **G)** ARfl binding sites as seen by AR C-terminal antibody ChIP-seq on samples in (E).

To determine whether the lack of initial transcriptional activity was due to failure of ARv7 accumulation in the nucleus, we used immunofluorescence (IF) to examine ARv7 localization. ARv7 was nuclear in both the short-term ENZ-treated cells and in the VCaP16 cells, indicating ARv7 lack of activity was not due to exclusion from the nucleus (**Figure S3A**). Interestingly, ARfl was also primarily nuclear in both the short-term ENZ-treated VCaP (VCaP-E) and VCaP16 cells (**Figure S3B**). To further delineate ARv7 nuclear localization we performed additional co-immunofluorescence of ARv7 and CBP, followed by confocal microscopy. ARv7 was localized primarily in discrete puncta both at 4 days following ENZ treatment and in the VCaP16 cells (**Figure 3B**). There was substantial colocalization of CBP in these structures, consistent with them being transcriptional condensates. Although the number of puncta per cell was greater in the VCaP16 cells (**Figure 3C**), a similar fraction was colocalized with CBP in the VCaP16 cells versus the VCaP-E cells (**Figure 3D**).

These observations indicated that ARv7 expressed acutely following ENZ treatment partitions into discrete structures consistent with condensates, and that its initial lack of transcriptional activity reflects a defect at a subsequent step. Therefore, we next assessed ARv7 chromatin binding by immunoblotting and ChIP-seq in VCaP cells following 3 days in ENZ (VCaP-E) versus in VCaP16 cells. Notably, ARv7 protein levels were comparable after formaldehyde crosslinking in the chromatin fraction of VCaP-E and VCaP16 cells **(Figure 3E**). In contrast, ChIP-seq showed that ARv7 chromatin binding at specific sites was low in VCaP-E cells, and was substantially increased in the VCaP16 cells (2150 common sites in replicate samples) (**Figure 3F**). ARfl binding was also greater in the VCaP16 cells versus the VCaP-E cells, although the fold increase was less than that for ARv7 (**Figure 3G**). Notably, the number of identified ARfl sites in the VCaP16 cells (52,536 sites) was much greater than the number of ARv7 sites, which may reflect more binding and/or relative efficiency of the available ChIP antibodies (see below). Finally, acute DHT treatment (4 hours) caused an increase in ARfl binding (as expected), but also increased ARv7 binding in the VCaP16 cells, which may reflect mechanisms including ARfl/ARv7 heterodimer formation (see below). Together these data show that the adaptation to ENZ is associated with an increase in the ability of ARv7 to bind specific sites on chromatin.

### ARv7 binds chromatin independently of ARfl

The persistent AR activity in VCaP16 cells after depletion of ARfl indicated that ARv7 can transactivate independently of ARfl. Consistent with this hypothesis, degradation of ARfl using ARCC-32 did not impair ARv7 chromatin binding in VCaP16 cells (**Figure 4A**). We further examined the ARCC-32-treated VCaP16 cells by ChIP-seq. While there was a modest decrease in ARv7 peak intensities at shared sites in response to the ARCC-32 (**Figure 4B**), the number of sites was not decreased, with a trend towards an increased number of ARv7 binding sites in the ARCC-32-treated cells (**Figure 4C**). Moreover, the majority of the 3,480 ARv7 sites found in the control VCaP16 cells were also present in the ARCC-32-treated cells (2,682, 77%) (**Figure 4D**).

**Figure 4.**
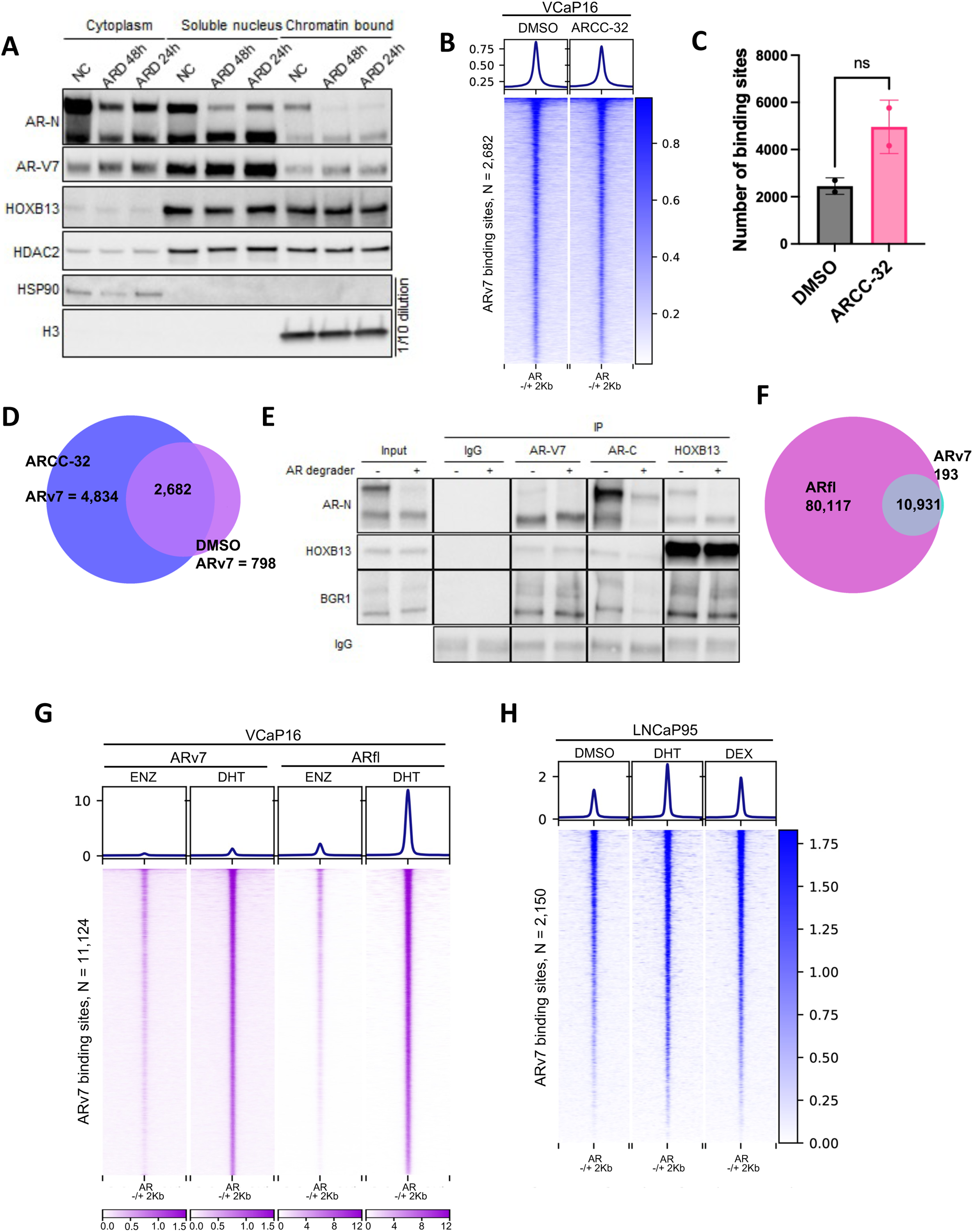
ARv7 chromatin binding in VCaP16 cells is independent of ARfl. **A)** VCaP16 cells in basal medium (16 μM ENZ) were treated for 24 or 48 hrs with vehicle (NC) or ARCC-32 (500 nM). Cells were then separated into cytoplasmic, soluble nuclear, or chromatin fractions and immunoblotted as indicated. **B)** Heatmap showing the intensity of ARv7 binding in VCaP16 cells in their basal medium (with 16 μM ENZ) with addition of DMSO or ARCC-32 (500 nM) for 36 hrs. ChIP-seq signal is centered at ARv7 sites common between both conditions. **C)** Number of ARv7 binding sites in VCaP16 cells in basal medium (with 16 μM ENZ) treated with DMSO or ARCC-32 (500 nM) for 36 hrs. **D)** Venn diagram showing the intersection of merged ARv7 sites in VCaP16 cells in DMSO (DMSO) or ARCC-32 (500 nM). **E**) VCaP16 cells were double crosslinked with DSP and formaldehyde. The chromatin fraction was then isolated and solubilized by DNA digestion with benzonase, followed immunoprecipitation with ARv7, AR C-terminal (ARfl), or HOXB13 antibodies and immunoblotting as indicated. **F)** Venn diagram showing the intersection of ARv7 and ARfl sites in VCaP16 cells following 4 hr treatment with DHT (10 nM). **G)** Heatmap showing increased binding intensity of ARv7 and ARfl in VCaP16 cells following 4 hr treatment with DHT (10 nM). ChIP-seq signal is centered at ARv7 sites identified in VCaP16 cells following 4 hr treatment with DHT (10 nM). **H)** Heatmap showing the enrichment of ARv7 binding in LNCaP95 cells in basal medium treated for 2 hrs with DMSO, DHT (10 nM) (DHT), or dexamethasone (DEX, 100 nM). ChIP-seq signal is centered at ARv7 sites identified in VCaP16 cells.

We next used the AR degrader ARCC-32 to determine whether ARv7 complexes on chromatin are dependent on ARfl. For these studies cells were initially treated with ARCC-32, and protein complexes on chromatin were then stabilized by double fixation with the cleavable cross-linker DSP and formaldehyde. This was followed by chromatin isolation and solubilization by DNA digestion with benzonase. ARCC-32 again did not diminish the amount of ARv7 associated with chromatin (**Figure 4E**). Immunoprecipitation with an ARv7 specific antibody showed that the trace levels of ARfl that were lost in the ARCC-32-treated cells which could reflect heterodimers or possibly adjacent homodimers on DNA. Notably, ARCC-32 did not decrease the amount of HOXB13 or BRG1 that co-precipitated with ARv7, indicating these interactions were not dependent on ARfl. Finally, immunoprecipitation with a HOXB13 antibody further demonstrated that the ARv7 interaction with HOXB13 was independent of ARfl. These findings in VCaP16 cells are consistent with results we recently reported in LNCaP95 and CWR22Rv1 cells, where we similarly found that ARv7 was binding to chromatin independently of ARfl ^31^.

While these data indicate that ARv7 does not act predominantly as a heterodimer with ARfl in VCaP16 cells, studies with ectopically expressed ARv7 clearly show it can act both as a homodimer and heterodimerize with ARfl ^32–35^. A previous study in LNCaP95 cells (LNCaP cells adapted to androgen depleted medium) also found that DHT stimulation increased ARv7 chromatin binding, suggesting ARv7 heterodimerization with ARfl ^27^. This DHT-stimulated increase in ARv7 binding was similarly observed in the VCaP16 cells (see Figure 2F). The gained ARv7 binding sites in the VCaP16 cells overlapped with ARfl sites, consistent with ARfl/ARv7 heterodimerization (**Figure 4F**). Notably, a heat map of the ChIP-seq data indicates that DHT broadly and proportionately enhances ARv7 binding to sites that are bound more weakly prior to DHT (**Figure 4G**).

In addition to dimerization, another mechanism by which ARfl may enhance ARv7 binding is through assisted chromatin loading, wherein binding of an ARfl homodimer to a site makes the site more available for subsequent binding by ARv7. Indeed, a previous study found that glucocorticoid receptor (GR) binding at a glucocorticoid responsive element enhances binding of a variant GR to the same site ^36^. To test whether ARv7 binding might be enhanced by binding of a steroid receptor at the same sight, we took advantage of the overlap between GR and ARv7 binding sites and asked whether GR activation would enhance ARv7 binding at these sites. LNCaP95 cells were treated with vehicle, DHT, or dexamethasone (DEX, GR specific agonist), followed by ARv7 ChIP-seq. We then examined ARv7 binding to the *FKBP5* gene, which is both AR and GR regulated. Consistent with assisted loading, DEX enhanced ARv7 binding at multiple sites in this gene (**Figure S4A**). Similarly, ARv7 binding was increased by DEX in the AR/GR (and mineralocorticoid receptor, MR) regulated *SCNN1A* gene. We then examined two previous GR ChIP-seq data sets to identify GR/ARv7 shared sites, and found that DEX caused a broad increase in ARv7 binding to these sites (**Figure S4B, C**). This broad increase was also observed when we examined all ARv7 sites (**Figure 4H**). Together these findings indicate that ARv7 functions independently of ARfl in the VCaP16 cells, but in the presence of androgen can function cooperatively with ARfl through heterodimerization, assisted loading, or possibly other mechanisms. The results also suggest that GR, in addition to directly activating a subset of AR-target genes ^37^, may indirectly enhance ARfl and/or ARv7 activity by assisted loading. Notably, GR mRNA was not increased in VCaP16 versus VCaP-E cells, while there was a modest increase in MR expression (∼31% increase).

### ARv7 cistrome is enriched for higher affinity ARfl binding sites

Previous studies have been inconsistent as to whether ARv7 has a cistrome and/or transcriptome that is distinct from ARfl ^26, 38–42^. In LNCaP95 cells it was found previously that essentially all ARv7 binding sites were also ARfl sites ^27^. Similarly, ARv7 ChIP-seq in VCaP16 cells revealed consistent ∼2150 peaks, virtually all of which (97%) overlapped with ARfl peaks (**Figure 5A, B**). Notably, while the number of called ARv7 peaks is low relative to ARfl peaks, inspection of the heat maps centered on total ARfl sites indicates that there is a correlation between ARfl and ARv7 binding across most or all ARfl peaks **(Figure 5C**). It is not clear whether the lower number of called ARv7 peaks reflects weaker ARv7 binding to chromatin or lower efficiency of the ARv7 antibody used for ChIP (or both). In support of the former hypothesis, profile plots clearly show that ARfl binding is markedly greater at called ARv7 sites, suggesting these are being picked up by the ARv7 antibody as they reflect higher affinity sites for ARv7 as well as for ARfl (**Figure 5D**). Notably, ARfl binding is also greater in the VCaP16 cells relative to the VCaP-E cells, suggesting that adaptations in the VCaP16 cells that increased ARv7 binding may also increase binding of ENZ-liganded ARfl (**Figure 5E**). As expected, DHT increased ARfl binding in both VCaP and VCaP16 cells.

**Figure 5.**
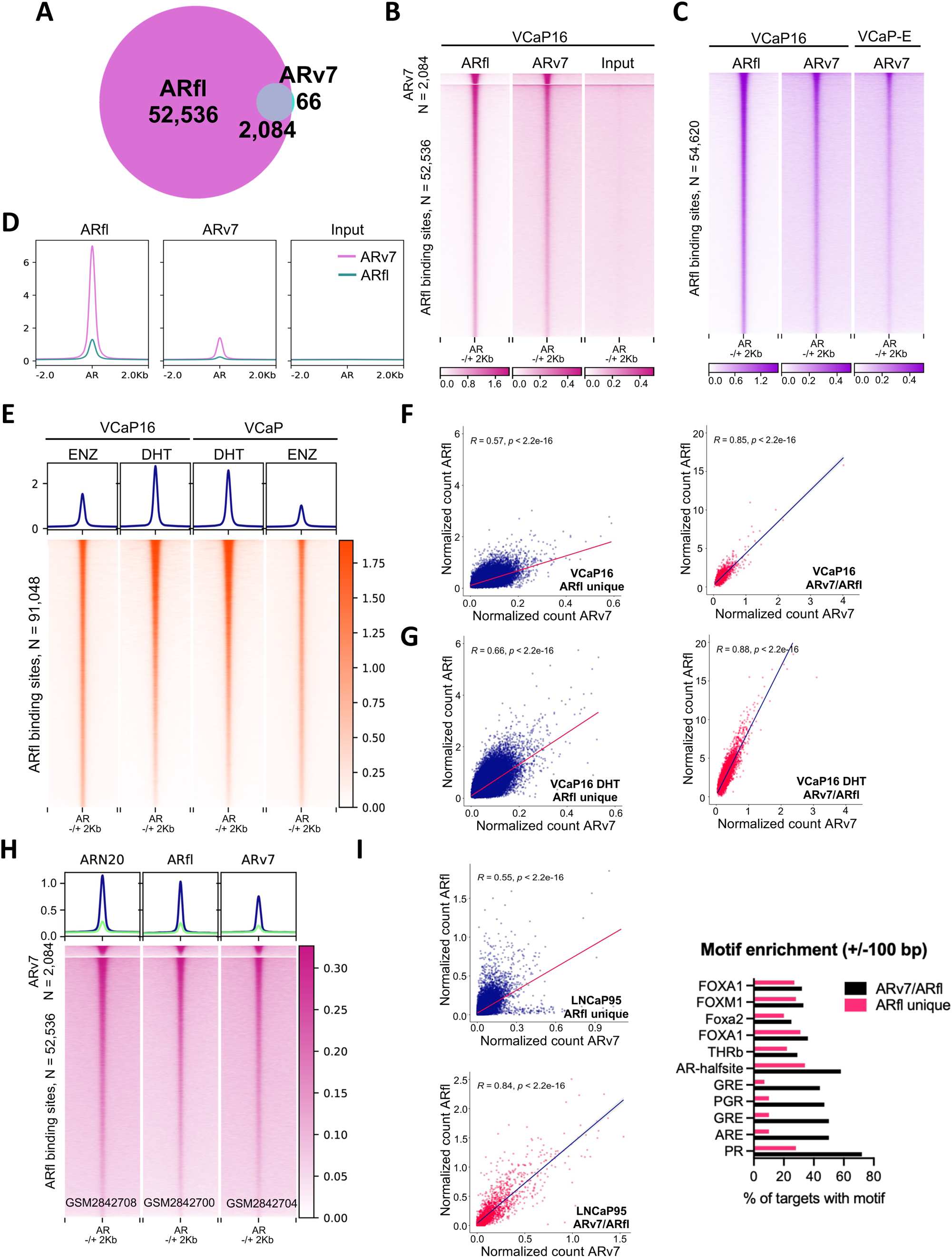
ARv7 binding sites overlap with ARfl binding sites with proportionate peak intensities. **A)** Venn diagram showing the intersection of ARv7 and ARfl sites in VCaP16 cells in basal medium (with 16 μM ENZ). **B)** Heatmap showing correlation between ARfl and ARv7 binding at called ARv7 sites (upper) and at ARfl unique sites. **C)** Heatmap showing correlation between ARv7 and ARfl binding intensities across all ARfl sites in VCaP16, and showing ARv7 in VCaP after short term ENZ treatment (4 days) (VCaP-E). **D)** Profile plots showing peak intensities for ARfl and ARv7 in VCaP16 cells at ARv7 versus ARfl unique binding sites. ChIP-seq signal is centered at ARv7/ARfl shared (ARv7) and ARfl unique (ARfl) sites identified in VCaP16. Input sample does not show signal enrichment at centered genomic regions. **E)** ARfl binding intensities in 1) VCaP16 maintained in basal medium with 16 μM ENZ, 2) VCaP16 treated with DHT (10 nM) for 4hrs, 3) VCaP cells treated with ENZ for 72 hrs followed by removal of ENZ and addition of DHT (10 nM) for 4 hrs, and 4) VCaP cells treated with ENZ for 72 hrs. ChIP-seq signal is centered at total ARfl sites identified in VCaP16 cells stimulated with DHT (10 nM) for 4hrs. **F)** Pearson’s correlation of normalized read counts: ARv7 versus ARfl called at shared ARfl/ARv7 and ARfl unique sites in VCaP16 cells and **G)** in VCaP16 cells treated with DHT (10 nM) for 4hrs. **H)** Heatmap showing the AR signal enrichment with AR N-terminal (GSM2842708), AR C-terminal (GSM2842700), and ARv7 (GSM2842704) antibodies in LNCaP95 cells. ChIP-seq signal is centered at shared ARfl/ARv7 (ARv7) sites and ARfl unique sites identified in VCaP16 cells**. I)** Pearson’s correlation of normalized read counts: ARv7 (GSM2842704) versus ARfl (GSM2842700) in LNCaP95 cells at shared ARfl/ARv7 (ARv7) sites and ARfl unique sites identified in VCaP16 cells. **J)** Top enriched motifs (+/−100 kb from the peak center) at ARfl unique sites and at shared ARfl/ARv7 sites in VCaP16 cells.

To further assess the relationship between ARv7 versus ARfl binding in VCaP16 cells, we plotted normalized read counts for ARv7 versus for ARfl at ARfl unique sites (sites where ARv7 was not called by the HOMER algorithm) and at ARv7 sites (which are all shared with ARfl) (**Figure 5F**). Significantly, ARv7 binding was highly correlated with ARfl binding at ARv7 sites and at ARfl unique sites, supporting the conclusion that ARv7 is also binding proportionately to ARfl at these latter ARfl unique sites. We also compared ARv7 versus ARfl binding in VCaP16 cells after DHT treatment, and saw a similar strong correlation at ARv7 and ARfl unique sites (**Figure 5G**). Interestingly, the slope of the line (ratio of ARfl to ARv7 binding) was greater in the DHT-treated cells, consistent with DHT driving stronger binding of ARfl.

We next reexamined previous ARfl and ARv7 ChIP-seq data in LNCaP95 cells ^27^ to determine whether ARv7 sites identified in VCaP16 cells are similarly higher affinity sites for ARfl and ARv7 in LNCaP95. Indeed, peak intensities with an AR N-terminal Ab, AR C-terminal Ab, and ARv7 Ab were all markedly greater at VCaP16 ARv7 sites versus ARfl unique sites (**Figure 5H**). There was also a similar correlation between ARfl and ARv7 binding intensities at ARfl unique and ARv7 sites identified in the LNCaP95 cells (**Figure 5I**). Examination of additional AR ChIP-seq data sets in CWR22RV1, LNCaP95, and LNCaP further showed increased ARv7 binding at the ARv7 sites identified in VCaP16 cells, and indicated that these correspond to high affinity ARfl binding sites (**Figure S5A, B, C**). Moreover, an analysis of normal prostate, primary PC clinical samples and PDX CRPC models also showed increased ARfl binding at ARv7 binding sites identified in VCaP16 (**Figure S5D**) ^43, 44^. Finally, we found that the majority of VCaP16 ARv7 binding sites (89.3%) overlapped ARfl sites that were shared by at least two tumors in an AR ChiP-seq analysis of 88 primary PC ^45^ (**Figure S5E**).

We then carried out a motif analysis comparing ARv7 and ARfl binding sites to determine whether identified ARv7 binding sites are associated with distinct features that may mediate increased binding. Notably, relative to ARfl unique sites, the ARv7 sites were greatly enriched for the AR/GR/PR motif, consistent with them being high affinity binding sites (**Figure 5J, Figure S6**). Taken together these results show that the ARv7 and ARfl cistromes in VCaP16 are largely overlapping, and also overlapping with the cistrome of the DHT liganded AR. Moreover, they show that ARv7 binding intensity correlates with affinity for the DHT liganded AR, and that called ARv7 sites in VCaP16 are high affinity ARfl sites across multiple models and clinical samples.

### ARv7 transcriptome overlaps with that of ARfl

We next compared the initial effects of ENZ treatment in VCaP cells with siRNA-mediated depletion of ARv7 in VCaP16 cells. This showed a very strong correlation between genes regulated by ARfl in VCaP cells and genes regulated by ARv7 in VCaP16 cells, indicating that the ARfl transcriptome in VCaP cells was largely restored by ARv7 in VCaP16 cells (**Figure 6A**). A previous study in LNCaP95 cells had found a marked difference in the ARfl versus ARv7 transcriptomes, with ARv7 acting to suppress expression of a subset of growth promoting genes ^27^. Notably, the previous study in LNCaP95 cells compared effects of depleting ARfl versus ARv7, whereas the analysis here compared blocking ARfl in VCaP cells with ARv7 depletion in VCaP16 cells. Therefore, we next compared effects of depleting ARfl versus ARv7 in VCaP16 cells, as was done previously in LNCaP95 cells. In this case there was a much weaker correlation between genes altered in VCaP16 by depletion of ARfl (exon 7 siRNA) versus depletion of ARv7 (ARv7 siRNA), with ARfl depletion increasing the expression of many genes that are decreased by ARv7 depletion (**Figure 6B**).

**Figure 6.**
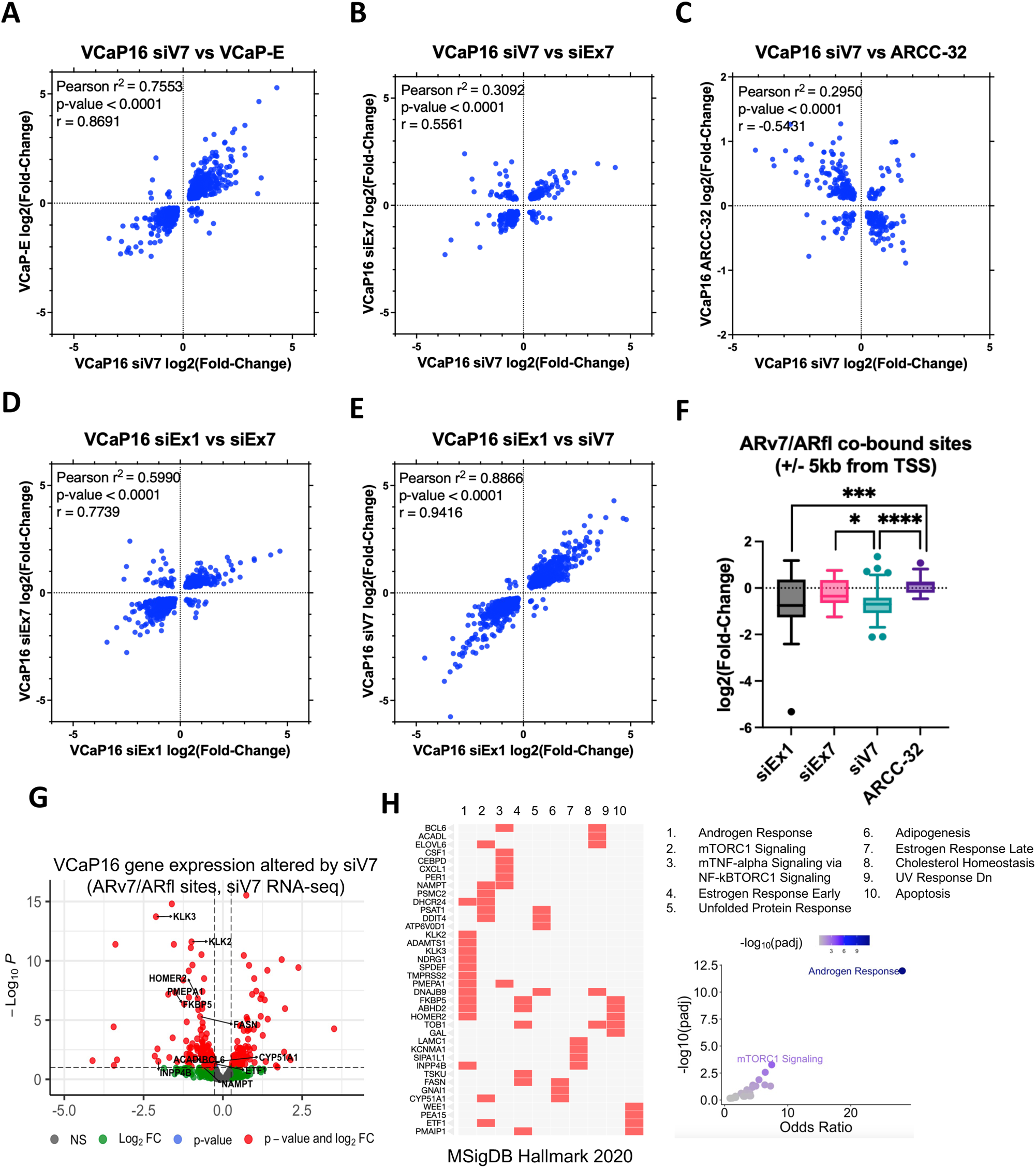
ARv7 transcriptome in VCaP16 is highly correlated with ARfl in VCaP. **A)** Correlation of log2(Fold-Change) values of significantly differentially expressed genes (padj < 0.05) in RNA-seq data from (x-axis) VCaP16 with siRNA knockdown of ARv7 versus non-targeting control (VCaP16 siV7) and (y-axis) VCaP cells treated with ENZ for 4 days versus parental VCaP cells (VCaP-E). **B)** Correlation of (x-axis) VCaP16 with siRNA knockdown of ARv7 versus non-targeting control (VCaP siV7) and (y-axis) VCaP16 with siRNA knockdown of Exon7 (targets ARfl) versus non-targeting control (VCaP siEx7). **C)** Correlation of (x-axis) VCaP16 with siRNA knockdown of ARv7 versus non-targeting control (VCaP16 siV7) and (y-axis) VCaP16 treated with ARCC-32 (500 nM) versus VCaP16 in DMSO (VCaP16 ARCC-32). **D)** Correlation of (x-axis) VCaP16 with siRNA knockdown of Exon1 (targets ARfl and ARv7) versus non-targeting control (VCaP siEx1) and (y-axis) VCaP16 with siRNA knockdown of Exon7 (targets ARfl) versus non-targeting control (VCaP siEx7). **E)** Correlation of (x-axis) VCaP16 with siRNA knockdown of Exon1 (targets ARfl and ARv7) versus non-targeting control (VCaP siEx1) and (y-axis) VCaP16 with siRNA knockdown of ARv7 versus non-targeting control (VCaP16 siV7). **F)** Differential expression (log2(Fold-Change) values) of ARv7-regulated genes (where ARv7 binding sites are located within +/− 5 kb from TSS) altered by siV7, siEx1, siEx7 or ARCC-32 in RNA-seq from VCaP16 cells. Differentially expressed genes with padj < 0.05 (siV7, siEx1, siEx7) and p-value < 0.05 (ARCC-32) were included. Mann-Whitney U, two-tailed: siV7 vs siEx7, p-value = 0.0308; siV7 vs siEx1, p-value = 0.8151; siV7 vs ARCC-32, p-value < 0.0001; siEx1 vs siEx7, p-value = 0.0609; ARCC-32 vs siEx7, p-value = 0.1148; ARCC-32 vs Ex1, p-value = 0.0005. **G)** Volcano plot of differentially expressed genes linked to ARv7 binding sites in VCaP16 cells (+/− 100 kb from TSS) in RNA-seq data from VCaP16 with siRNA knockdown of ARv7 (siV7) versus non-targeting control. **H)** Enrichment analysis by Enrichr of downregulated genes from G.

Similar results were obtained when the AR degrader ARCC-32 was used to deplete ARfl (**Figure 6C**). We also compared effects of an siRNA against exon 1 (targeting both ARfl and ARv7) with siRNA targeting ARfl (exon 7 siRNA) (**Figure 6D**) or ARv7 (**Figure 6E**). Notably, the strongest correlation was with the latter ARv7 siRNA, which further supports the conclusion that ARv7 is the predominant driver of AR activity in the VCaP16 cells. We next looked specifically at genes associated with ARv7 binding sites. These were greatly decreased by siRNA targeting both ARfl and ARv7 (Exon 1 siRNA) and by siRNA targeting just ARv7, further showing that these genes are being driven primarily by ARv7 (**Figure 6F**). In contrast, the siRNA targeting just ARfl (Exon 7 siRNA) had a much more modest effect on the ARv7 associated genes, while ARfl depletion with ARCC-32 modestly increased these genes.

The enhanced expression of many ARv7 stimulated genes and the observed trend towards augmentation in the Hallmark AR Responsive gene set (see Figure 2F) following ARfl depletion were surprising. We hypothesized that this enhancement in VCaP16 cells is a result of coactivator proteins being sequestered by the ENZ-liganded ARfl, and these coactivators become available for ARv7 coactivation upon ARfl depletion. To test this hypothesis, we carried out ChIP-seq for H3K27ac, BRD4, and p300 in VCaP16 cells treated with vehicle or ARCC-32. Consistent with the RNA-seq results, ARCC-32 did not have a clear effect on H3K27Ac at ARv7 or ARfl binding sites (**Figure S7A**). Moreover, it did not have an overall effect on BRD4 at ARv7 or ARfl binding sites (**Figure S7B**). In contrast, p300 binding was decreased (**Figure S7C**). We then examined two directly ARv7 regulated genes (with ARv7 binding sites) whose expression was increased by ARCC-32, GRIN3A and POLR2M (**Figure S7D**). For both genes, enrichment of H3K27Ac and BRD4 at ARv7 binding sites was increased in response to ARCC-32, although p300 was decreased, possibly reflecting a greater role for CBP versus p300 as a coactivator for ARv7 (**Figure S7E, F**). Overall, these findings show that ARfl depletion does not significantly decrease ARv7 activity. Instead, it appears to enhance ARv7 activity at a subset of genes, an effect that may be attributed to an increase in coactivator availability.

Further examination of the genes with called ARv7 binding sites that were altered in response to ARv7 siRNA showed downregulation of multiple canonical AR-regulated genes such as *KLK2* and *KLK3* (**Figure 6G**). These genes were highly enriched for the Hallmark Androgen Response gene set, and also significantly enriched for gene sets related to mTORC1 signaling, TNFα signaling, and lipid metabolism, indicating that ARv7 plays a key role in the restoration of these metabolic functions (**Figure 6H**).

### ARv7 binding sites are enriched for enhancer marks

The predominant role of ARv7 in driving the AR transcriptome suggested that sites where ARV7 was binding may have particularly strong enhancer activity. Therefore, we next carried out ChIP-seq for marks of active enhancers (H3K27ac and H3K4me1) in VCaP and VCaP16 cells. Indeed, centering on all AR sites in VCaP16 cells, H3K27ac was higher at ARv7 sites versus ARfl unique sites (**Figure S8A**). This same finding was observed in VCaP cells, indicating that the sites where ARv7 binds in VCaP16 are also potent enhancers in VCaP, which are presumably being driven by the androgen liganded ARfl. Notably, H3K27ac peak intensities across ARfl unique and ARv7 sites were comparable in VCaP16 versus VCaP cells, consistent with the restoration of AR transcriptional activity in the VCaP16 cells. The results for H3K4me1 were similar to H3K27ac, with increased signal at ARv7 sites in both VCaP16 and VCaP cells (**Figure S8B**). Taken together these data indicate that ARv7 sites are enriched for enhancer activity, but do not provide a basis for the gain in ARv7 binding to chromatin in the VCaP16 cells.

### Adaptation to ENZ is associated with increased chromatin accessibility

We next used ATAC-seq to determine whether the adaptation to ENZ in the VCaP16 cells and gain in ARv7 chromatin binding was associated with alterations in chromatin accessibility. Overall, there was an increase in the global intensity and number of ATAC sites in VCaP16 versus VCaP or VCaP-E cells(**Figure 7A, B**). We then identified the subset of ATAC sites that had significantly increased peak intensity in VCaP16 versus VCaP (ATAC UP sites), as well as sites that were not significantly changed and a small number that were decreased in peak intensity (**Figure 7C**). Profile plots indicate that chromatin accessibility at the ATAC UP sites was modestly increased after short term ENZ (VCaP-E cells), and that there was also a modest overall increase in sites identified as unchanged in VCaP16 versus VCaP (Unchanged sites) (**Figure 7D**).

**Figure 7.**
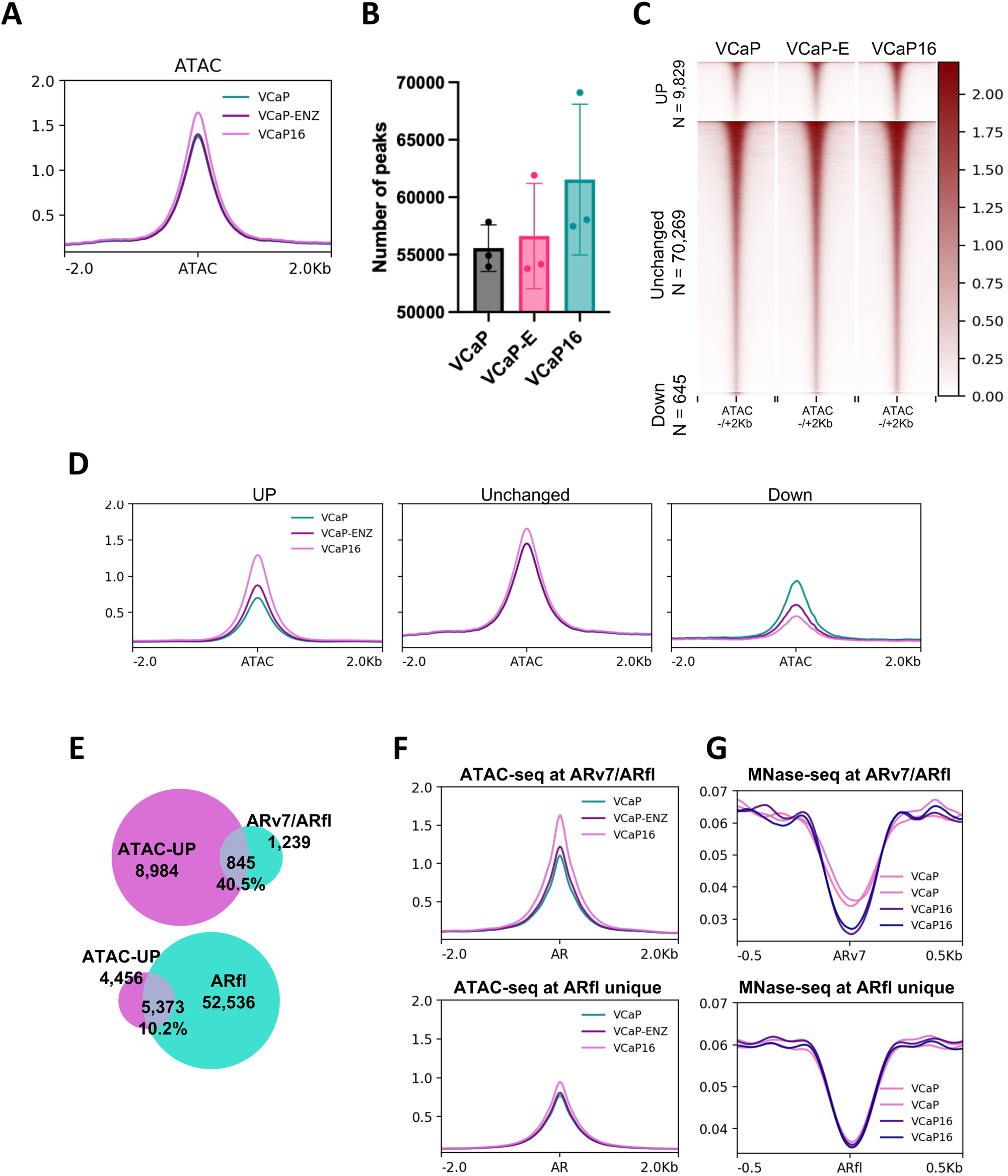
Chromatin accessibility in VCaP16 cells is increased at ARv7 binding sites. **A)** Profile plot showing ATAC-seq signal in parental VCaP cells, VCaP cells treated with ENZ for 4 days (VCaP-E) and VCaP16 (centered on all sites). **B)** ATAC-seq sites in parental VCaP cells, VCaP cells treated with ENZ for 4 days (VCaP-E) and VCaP16. **C)** Heatmap and **D)** profile plots showing the ATAC-seq signal strength in parental VCaP cells, VCaP cells treated with ENZ for 4 days (VCaP-E) and VCaP16. ATAC-seq signal is centered at sites with increased ATAC signal (UP sites), sites called as unchanged (Unchanged sites), or sites with decreased ATAC signal (Down sites) in VCaP16 versus parental VCaP cells as identified with CoBRA pipeline. **E)** Overlap of differentially more accessible sites (ATAC-UP) with shared ARv7/ARfl (upper panel) and ARfl unique binding sites (lower panel) in VCaP16 cells. **F)** Profile plots showing ATAC-seq signal enrichment and **G)** nucleosome occupancy by MNase-seq at ARv7/ARfl (upper panels) and ARfl unique binding sites (lower panels) in VCaP16 cells.

Notably, while the ATAC-UP sites were only a minority of the total ATAC sites, about 40% of ARv7 sites overlapped with these sites (**Figure 7E**). In contrast, only about 10% of ARfl unique sites overlapped with these ATAC-UP sites. We then assessed ATAC-seq signal centered on ARv7 binding sites, and found this was increased in VCaP16 versus VCaP (**Figure 7F**). In contrast, the ATAC-seq signal at ARfl unique sites was lower and much more modestly increased in the VCaP16 cells. These findings indicated that overall chromatin accessibility was increased in the VCaP16 cells, particularly at ARv7 binding sites.

The transposase used for ATAC-seq may also generate shorter fragments in areas of chromatin that are nucleosome depleted. Therefore, we next carried out an analysis of ATAC-seq fragment length data to determine whether there were shorter fragments generated at ARv7 sites in VCaP16. However, this analysis did not reveal fragment length differences between ARv7 and ARfl sites in VCaP or VCaP16 cells (**Figure S9**). To further directly test the hypothesis that chromatin around ARv7 sites was more open in VCaP16, we carried out MNase-seq to identify nucleosome protected versus exposed regions. Notably, we found greater depletion of nucleosomes around ARv7 sites in the VCaP16 cells versus the VCaP cells (**Figure 7G**). In contrast, there was no clear difference between VCaP16 and VCaP at ARfl unique sites. Overall, these results indicate that VCaP cells adapt to ENZ through nucleosome depletion around ARv7 binding sites to enhance ARv7 binding.

We next examined the LuCaP series of PDXs to identify models that may be driven by ARv7 to determine whether they had increased chromatin accessibility at ARv7 sites ^46^. We first used a recently published study to identify LuCaP PDXs that had the greatest increases in ARv7 when they relapsed after castration ^22^. In parallel we assessed enrichment for a gene set associated with ARv7 ^22^, which together indicated that the castration-resistant model LuCaP 77CR had the highest ARv7 activity, followed by 105CR and 136CR (**Figure S10A, B**). Finally, we found that one of two LuCaP 77CR and 96CR tumors, and two of two 105CR and 136CR tumors, formed a distinct cluster by unsupervised clustering of ARv7-signature genes expression (**Figure S10C**). Based on these findings, we examined LuCaP 105 versus LuCaP 105CR by ATAC-seq, which showed an overall increase in chromatin accessibility in LuCaP 105CR versus the parental castration-sensitive LuCaP 105 (**Figure S11A**). Moreover, ATAC-seq signal at ARfl sites identified in VCaP16 was also increased in the LuCaP 105CR versus parental 105 PDXs, and signal at ARv7 sites identified in VCaP16 was further markedly increased relative to the parental LuCaP 105 PDXs (**Figure S11B**). Together these findings are consistent with similar adaptive mechanisms driving AR reactivation in the PDX models and in VCaP16.

### Single cell ATAC-seq suggests an intermediate stage in the transition from VCaP to VCaP16

To study chromatin adaptations in VCaP16 versus VCaP on a single cell level, we carried out single cell ATAC-seq (scATAC-seq). Employing K-means clustering on the combined cell data (VCaP, VCaP-E, and VCaP16), we uncovered eight distinct clusters within the chromatin landscape (**Figure S12A**). Notably, cluster 1 was relatively specific to parental VCaP cells, while cluster 3 was relatively specific to VCaP16 cells. An intermediate cluster (cluster 2) was observed across all conditions, positioning itself between clusters 1 and 3 on a t-SNE plot. Our scATAC-seq analysis corroborated our previous findings from bulk ATAC-seq data, demonstrating higher chromatin accessibility at AR motifs in VCaP16 versus VCaP (**Figure S12B, S13A, S13B**). This heightened accessibility was particularly pronounced in the VCaP16 and VCaP-specific clusters (cluster 3 versus cluster 1) (**Figure S12B, S13C**). These findings suggest that the adaptive processes in VCaP16 cells are conducive to unlocking potent AR-binding sites. Notably, the chromatin accessibility analysis also revealed increased accessibility to AR motifs in VCaP-E compared to VCaP cells (**Figure S13A, B**). This observation suggests the initiation of chromatin changes following short-term exposure to ENZ.

Interestingly, cluster 2, the intermediate cluster that is common across all treatment states, exhibited chromatin features similar to those in cluster 3, specific to VCaP16. This observation leads to the hypothesis that cluster 2 may represent a pool of cells driving the development of ENZ resistance, ultimately giving rise to the population seen in cluster 3. To further investigate this hypothesis, we performed trajectory inference analysis. This analysis revealed a pseudotime progression from cluster 1 (VCaP-specific) through cluster 2 (intermediate) to cluster 3 (VCaP16-specific) (**Figure S12C**). The trajectory suggests a transition from the VCaP phenotype through an intermediate stage into the VCaP16 phenotype, providing insights into the evolution of ENZ resistance. A complementary analysis using another clustering algorithm revealed the same results (**Figure S14**). Additionally, clusters 4, 5, 6, 7, and 8 exhibited consistent chromatin structures across all cell lines (not shown).

### ARv7 activity in VCaP16 is dependent on Nuclear Factor I transcription factors

The most highly enriched transcription factor motifs at the ATAC-UP sites in VCaP16 were the common AR/GR/PR/MR motifs (**Figure 8A**). As expected, there was also enrichment for FOXA1, GRHL2, and HOXB13 motifs. However, the second most enriched motifs were for Nuclear Factor I (NFI) family transcription factors (NFIA, B, C, X). Comparison with ATAC-Unchanged and ATAC-Down sites similarly showed enrichment for the NFI motif at ATAC-UP sites (**Figure 8B**), and footprint analysis further supported enrichment of NFI transcription factors at ATAC-UP sites (**Figure 8C**).

**Figure 8.**
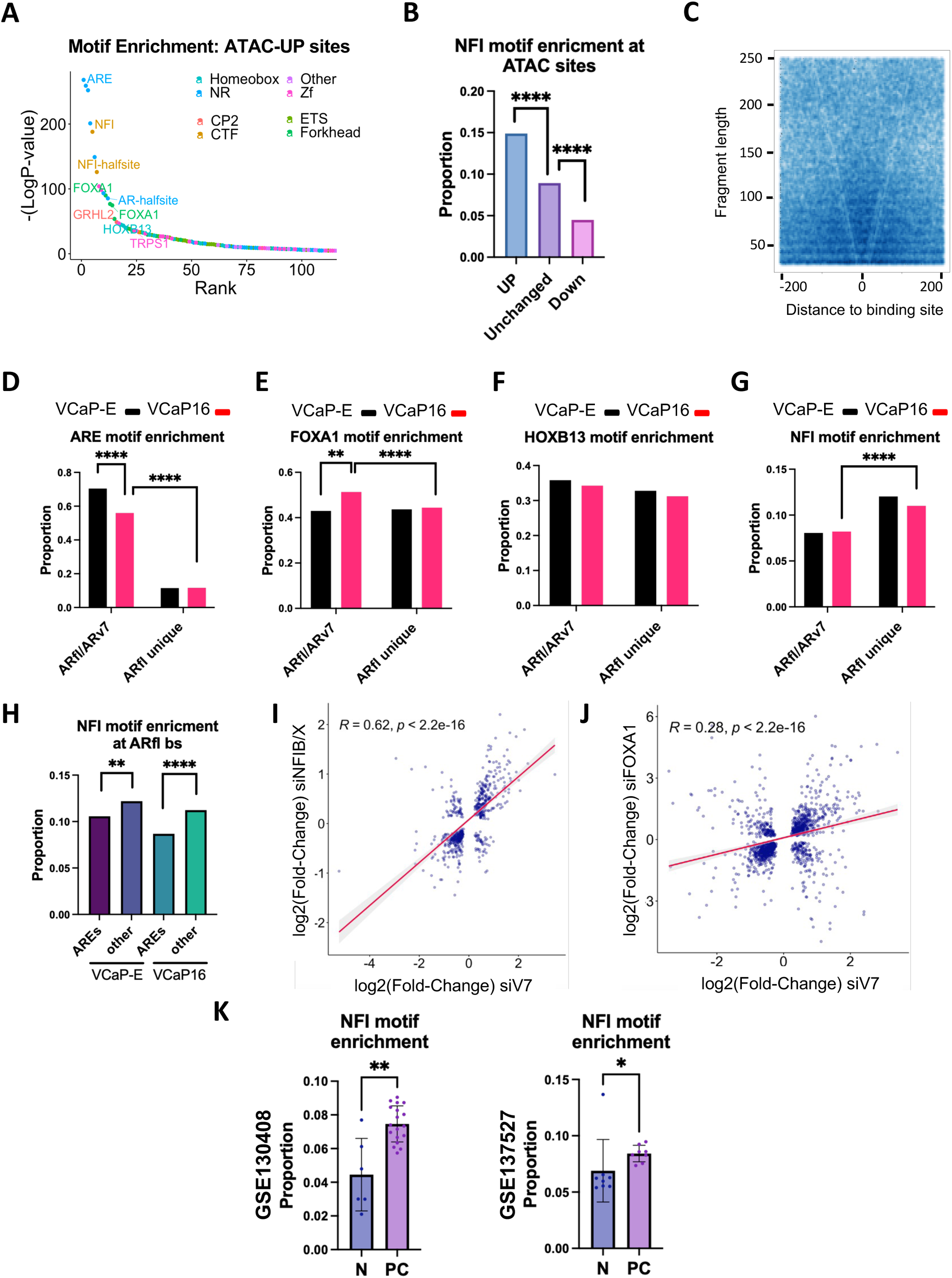
ARv7 activity in VCaP16 is dependent on Nuclear Factor I transcription factors. **A)** Motif enrichment analysis by HOMER at differentially more accessible sites (ATAC-UP) in VCaP16 cells. **B)** NFI motif enrichment by HOMER at ATAC-seq differentially more accessible sites (UP), unchanged (Unchanged) or differentially closed sites (Down) in VCaP16 versus parental VCaP cells as identified with CoBRA pipeline (****p<0.0001, Fisher Exact Test). **C)** Vplot showing NFI motif footprint at differentially more accessible sites (ATAC-UP) in VCaP16 cells. **D)** Enrichment for the ARE, **E)** HOXB13, **F)** FOXA1 and **G)** NFI motifs by HOMER at ARv7/ARfl shared and ARfl unique sites in VCaP cells after short term ENZ treatment (VCaP-E) versus in VCaP16 cells. **H)** NFI motif enrichment by HOMER at AR binding sites with canonical full AREs versus those with non-consensus AREs (others) in VCaP cells after short term ENZ treatment (VCaP-E) versus in VCaP16 cells. **I)** Pearson’s correlation of log2(Fold-Change) values of significantly differentially expressed genes (padj < 0.1) altered with siRNA knockdown of ARv7 (siV7) versus siRNA knockdown of NFIB and NFIX (siNFIB/X) and **J)** versus siRNA knockdown of FOXA1 (siFOXA1) in VCaP16 cells. **K)** NFI motif enrichment by HOMER at AR binding sites in normal prostate tissue (N) versus primary prostate cancer (PC) in two clinical datasets, GSE139498 and GSE137527 (*p<0.05, **p<0.01, Mann-Whitney test).

We next assessed for enrichment of the ARE, FOXA1, HOXB13, and NFI motifs at ARv7 and ARfl unique sites in VCaP-E cells versus in VCaP16 cells. Notably, while ARE and AR-half motifs were enriched at ARv7 versus ARfl unique sites in both cells, there was greater enrichment of the ARE motif at ARv7 sites in the VCaP-E versus VCaP16 cells, indicating that ARv7 binding was most highly biased towards canonical AREs after short-term ENZ treatment (**Figure 8D**, **Figure S15A**). Conversely, the FOXA1 motif was enriched at ARv7 sites in VCaP16 versus VCaP-E cells (**Figure 8E**), while enrichment for the HOXB13 motif was comparable at ARv7 sites in both cell lines (**Figure 8F**). In contrast to the FOXA1 and HOXB13 motifs, enrichment for the NFI motif was seen at ARfl unique versus ARv7 sites in both VCaP-E and VCaP16 cells, with greatest enrichment at ARfl unique sites in VCaP-E cells (**Figure 8G**). One interpretation of this finding is that NFI transcription factors preferentially support ARfl and ARv7 binding at weaker AREs. Therefore next we assessed the NFI motif at AR-binding sites with canonical versus non-canonical AREs, and found a significantly higher enrichment of the motif at non-canonical AREs (**Figure 8H**).

NFI proteins play major roles in the development of diverse tissues, and can have oncogenic or tumor suppressor functions in varying contexts ^47^. Moreover, NFI proteins can bind nucleosomes, control chromatin loop boundaries, and increase chromatin accessibility ^48–52^. Notably, previous studies in PC have identified interactions between AR, FOXA1 and NFI proteins, and in particular implicated NFIB in the regulation of AR activity and PC development ^53–56^. Examination of the Human Protein Atlas shows that both NFIB and NFIX are highly expressed in prostate (**Figure S16A**), with NFIX being most highly expressed amongst AR-expressing PC cell lines (**Figure S16B**). We confirmed by qRT-PCR the preferential expression of NFIX in LNCaP cells, and of NFIX and NFIB in VCaP cells (**Figure S16C**). Consistent with this pattern of expression, DepMap CRISPR and RNAi screens showed dependency on NFIB and NFIX, but not NFIA or NFIC, in AR-positive PC cells (VCaP and 22RV1), with no NFI dependency in AR-negative PC cells (DU145 and PC3) (**Figure S16C - F**). Finally, our RNA-seq data also showed higher NFIB and NFIX (versus NFIA and NFIC) in VCaP and VCaP16, although the difference was more modest and levels for each NFI were comparable in VCaP and VCaP16 cells (**Figure S16G**).

Based on these data we next assessed the effects of combined siRNA-mediated knockdown of NFIB and NFIX in VCaP16 cells. RNA-seq confirmed that the siRNA decreased both NFIB and NFIX, without compensatory increases in NFIA or NFIC (**Figure S17A**). This NFIB/X depletion did not have a consistent effect on canonical AR-regulated genes, with two such genes (ACPP and NKX3.1) being the most significantly increased and decreased, respectively (**Figure S17B**), and a modest decrease in the Hallmark Androgen Response gene set (**Figure S17C**). Hallmark gene sets related to cell cycle were more significantly decreased (E2F Targets, G2M Checkpoint, and MYC Targets), while there was an increase in the EMT gene set (**Figure S17C**). To more directly assess the contribution of NFI to ARv7 function, we compared the transcriptional effects of NFIB/X versus ARv7 depletion. Notably, these effects were strongly correlated (**Figure 8I**). There was also a correlation between transcriptional effects of ARv7 and FOXA1 knockdown, but the correlation with NFIB/X depletion was greater (**Figure 8J**).

To further interrogate the role of NFI proteins we looked specifically at expression of genes linked to ARv7 binding sites versus ARfl unique sites (which are weak ARv7 sites). There was a correlation between effects of ARv7 and NFIB/X knockdown at the former ARv7-binding site linked genes, and a stronger correlation with the latter ARfl-binding site linked genes, indicating that genes linked to these weaker sites are more dependent on NFIB/X (**Figure S18A**). A similar analysis for FOXA1 siRNA showed weaker correlations for both ARv7-and ARfl-unique site linked genes (**Figure S18B**). Interestingly, while the majority of Hallmark gene sets were decreased by ARv7 knockdown, many gene sets were increased by both FOXA1 and NFIB/X knockdown, suggesting cooperative effects of FOXA1 and NFI transcription factors at many genes that are not AR regulated (**Figure S17C**).

An overall conclusion from these results is that NFI proteins in ENZ-adapted cells contribute to an increase in chromatin accessibility to support ARv7 and ARfl chromatin binding, particularly at weak AR binding sites that do not have canonical AREs. Based on these observations we examined AR binding sites at genes that were altered by ARv7 siRNA in VCaP16 cells (see figure 6). Notably, only a small fraction of AR binding sites linked to these ARv7 regulated genes had canonical AREs (**Figure S19A**). We next similarly examined AR binding sites at genes that were altered by both depletion of ARv7 and NFIB/X, and found that most also had noncanonical AREs (**Figure S19B**). These findings further support the conclusion that NFI proteins enhance ARv7 activity at weak AREs.

Interestingly, previous studies have found that the AR cistrome in PC diverges from that in normal prostate epithelium, with AR binding sites in PC being associated with FOXA1 and HOXB13 motifs in PC ^43, 44^. Moreover, analysis of AR-binding sites in normal prostate versus primary PC shows that the sites in normal prostate are more highly enriched for canonical full AREs (**Figure S20A**). Notably, in two available data sets we found that the NFI motif was significantly enriched at AR-binding sites in PC versus normal prostate (**Figure 8K**) ^43, 57^. Furthermore, in both the normal prostate and PC tissues, the NFI motif was enriched at AR binding sites that did not have the canonical full ARE (**Figure S20B, C**). Together these findings indicate that NFI proteins may also contribute to AR reprogramming during PC development. Consistent with this hypothesis, a previous study found that NFIB levels by IHC were increased in PC versus matched normal prostate ^53^, although analyses of NFIB and NFIX mRNA in multiple data sets do not show clear increases in tumor versus normal tissues (**Figure S20D**).

## DISCUSSION

ARv7 and related AR variants have constitutive transcription activity and their increased expression is associated with development of resistance to available ASI therapies. However, the extent to which ARv7 drives the AR program, whether it is dependent on ARfl or has other unique dependencies, and whether it has novel functions, remain to be fully determined. To model the progression from androgen-dependence to ENZ resistance in patients we generated VCaP cells that were resistant to ENZ. We found that these VCaP16 cells had features observed most commonly in the clinic including restoration of the AR-transcriptional program, high level of ARfl expression, and increased expression of ARv7. Notably, knockdown or degradation of ARfl did not suppress the AR-transcriptional program in these cells, in contrast to knockdown of ARv7 that did markedly suppress the AR-transcriptional program. Consistent with these results, we showed biochemically that ARv7 complexes on chromatin were not dependent on or associated with ARfl. Significantly, while ARv7 was increased acutely in response to ENZ in the VCaP-E cells, this was not sufficient to drive the AR-transcriptional program. Together these observations indicated that further adaptations are needed to enhance ARv7 binding to chromatin and its ability to drive the AR-transcriptional program.

We determined that while ARv7 expressed in VCaP-E cells acutely exposed to ENZ was nuclear, it’s binding to DNA as assessed by ChIP-seq was low relative to ARv7 in the ENZ-adapted VCaP16 cells. This gain in ARv7 chromatin binding in VCaP16 cells was associated with a broad increase in chromatin accessibility as assessed by ATAC-seq. This increase in chromatin accessibility was greater at identified ARv7 binding sites relative to ARfl unique sites, where ARv7 binding is below the peak threshold, and the depletion of nucleosomes at ARv7 binding sites in VCaP16 cells versus parental VCaP cells was confirmed by MNase-seq. Transcription factor motifs enriched at sites with increased chromatin accessibility in VCaP16 cells (ATAC-UP sites) included those for AR, FOXA1, and HOXB13, consistent with previous studies showing that FOXA1 and HOXB13 can contribute to AR chromatin binding. However, the most enriched motif at ATAC-UP sites (after the AR motif) was the NFI transcription factor motif. Moreover, the transcriptional effects of depleting NFIB and NFIX in VCaP16 cells were strongly correlated with those of ARv7 depletion. Together these findings indicate a major role for NFI proteins in the enhancement of chromatin accessibility and in supporting ARv7 function.

Previous studies have shown varying overlap between the cistromes and transcriptomes of ARv7 and ARfl, and indicated that ARv7 may have distinct transcriptional functions ^26, 38–42^. However, in LNCaP95 cells (expressing endogenous ARfl and ARv7) we reported previously that all ARv7 binding sites detected by an ARv7 antibody (that were also detected by an AR N-terminal antibody) were also ARfl sites (detected by an AR C-terminal antibody) ^27^. We similarly found in VCaP16 cells that the vast majority of called ARv7 sites were also ARfl sites. While these results indicate that the ARv7 cistrome largely mirrors that of ARfl, this certainly does not rule out ARv7 unique sites in some contexts. Indeed, ZFX has been reported to be a novel ARv7 partner that drives ARv7 binding to unique non-canonical sites that are located at promoters and also require BRD4 ^58^. Very high levels of ARv7, either endogenous or exogenous, may possibly also drive ARv7 to unique sites. Conversely, some sites recognized by the available ARv7 antibodies may not be specific, particularly if they are not also detected by an AR N-terminal antibody. In other cases, ChIP with AR N-terminal antibody after ARfl knockdown may identify some sites bound by ARv7 that are not strong enough sites to meet statistical thresholds under basal conditions. More studies are clearly needed to understand further mechanisms that may drive ARv7 (or other AR splice variants) to sites that are not bound by the agonist liganded ARfl.

Notably, the number of ARfl sites in the VCaP16 cells, even in the presence of ENZ, was much greater than the number of called ARv7 sites. This may in part reflect more specific ARfl antibodies for ChIP, so that reads at more sites reach statistical significance over background, and/or intrinsically weaker ARv7 binding. In either case, we found that peak normalized counts for ARv7 were highly correlated with those for ARfl (in VCaP16 and LNCaP95 cells), and this correlation extended to ARfl peaks that were not called as ARv7 sites. These data indicate that the ARv7 cistrome in VCaP16 (and LNCaP95 cells) largely overlaps that of ARfl, but with ARv7 having lower affinity and hence with ARv7 binding sites being under called. Indeed, previous FRAP studies have shown that nuclear ARv7 has markedly higher mobility relative to agonist liganded ARfl, indicating a more rapid DNA binding off rate ^33, 35^. We propose this relatively weak ARv7 chromatin binding capacity is why adaptations that enhance chromatin accessibility are needed to support its activity in VCaP16 cells.

With respect to the ARv7 transcriptome, in our current studies we found a strong correlation between genes regulated by ARv7 in VCaP16 cells and those regulated by ARfl in VCaP, supporting the conclusion that ARv7 takes over the transcriptional functions of ARfl. In contrast, there was minimal correlation between effects of knockdown of ARv7 versus knockdown or degradation of ARfl. Indeed, ARfl degradation increased expression of many genes that were decreased by ARv7 knockdown. We propose that this reflects increased availability of some coactivator proteins that are being nonproductively engaged by the ENZ-liganded ARfl. Significantly, this coactivator redistribution could mitigate beneficial effects of ARfl degraders that are now being tested in the clinic. Notably, our previous study in LNCaP95 cells did not find a clear correlation between the transcriptional effects of ARv7 versus ARfl knockdown, with ARv7 knockdown increasing expression of some genes that were decreased by ARfl knockdown ^27^. Consistent with this previous result in LNCaP95, ARv7 knockdown increased expression of many genes that were decreased by ARfl degradation. This increase may reflect cofactor redistribution, a repressive function of ARv7 at some sites, and/or a greater ability of the ARfl versus ARv7 (despite ENZ in the medium) to transactivate at these genes.

Previous studies have found that compared to ARfl sites, ARv7 sites are associated with open chromatin, with consensus full AREs, and with HOXB13 ^40, 41^. Notably, HOXB13, but not FOXA1, has been found to bind directly to ARv7, consistent with an increased dependence on HOXB13 versus FOXA1 (which is associated with and interacts with ARfl) ^40, 41, 59, 60^. We similarly found that ARv7-binding sites in VCaP16 were enriched for consensus full AREs relative to ARfl unique sites. Interestingly, this enrichment was greatest in VCaP-E cells, indicating that increased chromatin accessibility in the VCaP16 cells was allowing ARv7 to bind to lower affinity AREs. We similarly found enrichment for the HOXB13 motif at ARv7 sites, but also for the FOXA1 motif. Moreover, enrichment of the FOXA1 motif at ARv7 sites was greater in the VCaP16 cells, supporting a role for FOXA1 in the transition to ARv7 driven ENZ resistance. Notably, in contrast to HOXB13 and FOXA1 motifs, the NFI motif was enriched at ARfl unique versus ARv7 sites. As these ARfl unique sites are also weak ARv7 sites, these findings, in conjunction with the enrichment of NFI at ATAC-UP sites, indicate that NFI plays a major role in the transition to ENZ resistance by depleting nucleosomes and hence broadly facilitating ARv7 binding, and in particular at weak sites. In support of this conclusion, the NFI motif was enriched at AR-binding sites that do not have canonical full ARE in multiple PC models. Moreover, transcriptional effects of ARv7 and NFIB/X depletion were highly correlated, consistent with NFI proteins broadly supporting ARv7 function.

NFI proteins (NFIA, B, C, X) bind DNA as homo-or heterodimers and can act as pioneer factors by binding to nucleosomes and increasing chromatin accessibility ^48–52^. In addition to roles in the development of diverse tissues, they have context dependent oncogenic or tumor suppressor functions ^47^. NFIB in small cell lung cancer promotes metastases by broadly increasing chromatin accessibility ^48^, while NFIB and NFIX in hair follicles act to maintain stem cell super-enhancers ^52^. Most recently NFIB was also found to facilitate licensing for DNA replication ^61^. Previous studies in PC cells have found an association of NFI proteins with AR-binding sites ^54–56^ and identified direct interactions between NFI proteins and FOXA1 that can stabilize an AR-FOXA1-NFI complex^55^. In functional studies, NFIB knockdown in LNCaP cells had variable effects on AR-regulated genes, while prostate tissue explants from *Nfib* gene knockout mice showed hyperplasia that may reflect an AR-independent effect as it was not responsive to castration ^54^. Our findings are consistent with these previously described interactions between NFI proteins and AR, and reveal a further major role for NFI proteins in the progression to ENZ resistance by increasing chromatin accessibility and supporting ARv7 binding, particularly at sites with weak AREs.

We further hypothesize that these results reflect a role for NFI proteins in the reprogramming of the AR that occurs during PC development ^43^, as the NFI motif is significantly enriched at AR-binding sites in PC versus in normal prostate epithelium. Notably, based on mRNA levels, NFI expression is not increased during PC development or the progression to ENZ-resistance, although one IHC study found increased NFIB protein in primary PC to adjacent normal prostate epithelium ^53^. However, NFI protein levels or activity may be modulated by alternative splicing or posttranslational mechanisms, with one recent study showing that arginine methylation by CARM1 facilitates the NFIB-mediated maintenance of open chromatin states in small cell lung cancer ^62^. Further studies to address NFI expression and activity during PC development and progression are warrented.

## METHODS

### Cell Culture

VCaP cells (ATCC, Manassas, VA) were maintained in DMEM with 10% fetal bovine serum (FBS). LNCaP95 cells ^63^ were maintained in RPMI 1640 medium supplemented with 10% charcoal/dextran-stripped fetal bovine serum (CSS), 1% L-Glutamine and 1% penicillin/streptomycin. Cell line identities were established by short tandem repeat profiling and interspecies contamination test (IDEXX). Cells were screened for mycoplasma with MycoAlert detection kit (Lonza) at receipt or thaw, and then monthly for the duration of project. Cells were employed for 20 passages only before new cell lines were thawed. For generation of VCaP16 ENZ-resistant model, we seeded cells on plates in standard parental media and, after overnight attachment, switched to medium with 2-fold increasing concentrations of ENZ, ending at 16 µM ENZ after ∼ 2 months. The VCaP16 cells were maintained in 16 μM ENZ, and medium on all cell models was changed every 2-3 days.

### Immunoblotting and immunoprecipitation

For immunoblotting cells were harvested in RIPA lysis buffer supplemented with a protease and phosphatase inhibitor cocktails (Thermo Fisher). Lysis was allowed to proceed for 30-60 minutes on ice, before cells were spun down at maximum RPM in a microfuge and cell debris pellets removed. Protein concentrations were determined by Bicinchoninic acid (BCA) assay (Thermo Fisher). Lysates were prepared with 4X Laemmli buffer (BioRad, #1610747) and β-mercaptoethanol. Samples were run on pre-cast 4-15% gradient gels (BioRad), and fast-transferred to nitrocellulose membranes using the TransBlot Turbo system (BioRad). Separation of cells into cytoplasmic, soluble nuclear, and chromatin fractions was carried out using a Subcellular Protein Fractionation Kit (Thermo Fisher #78840).

For assessment of ARv7/ARfl interaction, cells were incubated in PBS with 1.5 mM dithiobis (succinimidyl propionate) (DSP, Thermo Fisher #22585) for 20 min, followed by incubation in 1% formaldehyde for 10 min at room temperature. Cross-linking was quenched by adding glycine to 0.1 M. Cell nuclei were then extracted and resuspended in 1/2 dilution IP buffer (Thermo Fisher #87788) and sonicated for 2 min (5s on/5s off) in a Diagenode Bioruptor in 4 ℃ water bath with medium power setting. The sample was combined with 10×TBS and 10% Triton X-100 to give final concentration of 1X TBS/1%Triton X-100, and 100 mM MgCl2 was added to give 1 mM MgCl2 final concentration. Benzonase (Sigma #71205) was then added (90 U per 50 μL, which was determined to be optimum in initial studies) ^31^. Benzonase digestion was for 15 min at 37℃ on a 750 RPM shaker, and then 50 μL of each aliquot was processed for reverse cross-linking, followed by immunoblotting.

### Antibodies and Reagents

The antibodies and concentrations used for immunostaining and immunoblotting blotting were: anti-Vinculin (Santa Cruz Biotechnology, #sc-73614, 1:5000), anti-GAPDH (Cell Signaling Technologies, #5174S, 1:5000), anti-AR (NT) (RevMab Biosciences, #31-1135-00, 1:2000), anti-ARv7 (RevMAb Biosciences, #31-1109-00, 1:2000), anti-CBP-P300 (Thermo-Fisher, #MA5-13634, 1:50 for IF, 1:2000 for WB), anti-FOXA1 (Cell Signaling Technologies, #53528S, Rabbit Host, 1:2000), anti-FOXA1 (Sigma, #WH0003169M1, Mouse Host, 1:2000), anti-PSA (Cell Signaling Technologies, #5365S, 1:2000), anti-H3 (Cell Signaling Technologies, #4499S, 1:2000). The siRNA for non-targeting control (Dharmacon, #D-001810-01-05), ARfl (exon 7, Dharmacon, #VOZOB-000001, UCAAGGAACUCGAUCGUAUUU), ARv7 (Dharmacon, #SRYWD-000001, GUAGUUGUAUCAUGAUU) or both (exon1, Dharmacon, #SRYWD-000005, CAAGGGAGGUUACACCAAAUU) were purchased from Dharmacon. Enzalutamide was obtained from SelleckChem (TX, USA). AR degrader ARCC-32 was provided by Arvinus Pharmaceuitcals. RIPA buffer and Triton-X100 were purchased from Thermo Fisher. RNaseA and Dynabeads were purchased from Invitrogen.

### Immunofluorescence

Cells were fixed with 4% paraformaldehyde at room temperature on a rocker in the dark, and then washed three times with PBS. Fixed cells were washed with TBS and permeabilized with 0.1% Triton X-100 for 10 minutes at room temperature. After three washes with TBS, the sections were incubated with 5% normal donkey serum (Jackson ImmunoResearch Lab Inc, West Grove, PA) for an hour at room temperature. Slides were then incubated with mouse anti-Androgen Receptor C-terminal (ARfl) (1:100, Abcam #ab227678), rabbit anti-ARv7 (1:200, RevMab Biosciences, 31-1109-00), and mouse anti-CBP/p300 monoclonal antibody (NM11, Thermo-Fisher MA5-13634) overnight at 4°C. The slides were washed three times and incubated with Alexa Fluor 647 conjugated donkey anti-mouse secondary antibody (Jackson ImmunoResearch Lab, 1:300) or Cy3 conjugated donkey anti-rabbit secondary antibody (Jackson ImmunoResearch Lab, 1:300). Samples were counterstained with Hoechst 33342 (Invitrogen) and washed three times with TBS. The slides were mounted with Prolong Gold anti-fade mounting media (Invitrogen). A Zeiss LSM 880 Inverted Live-cell Laser Scanning Confocal Microscope with Airyscan module was employed for all imaging. Z-stack images were captured for all markers to generate a 3D dataset for quantification with Imaris 9.8.2 software (Oxford Instruments).

### Quantitative real-time PCR

RNA was extracted from cell lines using an RNeasy Plus kit (Qiagen) according to the manufacturer’s instructions. Subsequent qPCR was performed using the TaqMan Fast Virus 1-Step Master Mix and StepOnePlus Real-Time PCR System (Thermo Fisher). The primers purchased from TaqMan were: GAPDH (Thermo Fisher, #4310884E), AR-FL (Thermo Fisher, Hs001711772_m1), AR-v7 (Thermo Fisher, #4331348, Custom Assay ID: AI6ROCI), KLK3 (Thermo Fisher, Hs02576345_m1), KLK2 (Thermo Fisher, Hs00428383_m1), FKBP5 (Thermo Fisher, Hs01561006_m1), TMPRSS2 (Thermo Fisher, hs01120965_m1), and SLC45A3 (Thermo Fisher, Hs01026319_g1).

### Proliferation Assays

For proliferation of VCaP cells exposed to ENZ, cells were plated at 20,000 cells/well on day −2. Experimental and control treatments were conducted and initial readings taken on day 0. CyQuant® Direct Cell Proliferation Assay (Invitrogen) was employed for determination of cell number based on fluorescent DNA staining, with dead cell exclusion provided by CyQuant® Suppressor solution (Invitrogen). An equal volume of CyQuant® solution, prepared on treatment day, was added to cells in culture medium, incubated for 60 minutes at 37°C, and fluorescence read on a SpectraMAX iD3 microplate reader (excitation/emmission 485nm/525nm). To assess effects of depleting ARfl or ARv7, cells were treated with siARv7, ARCC-32 (obtained from Arvinas, New Haven, CT), or vehicle on days 0 and 4.

### siRNA Transfection

Transfection with RNAiMAX reagents (Invitrogen) was conducted per manufacturer’s instructions for up to 72 hours. The siRNA for non-targeting control (Dharmacon, #D-001810-01-05), ARfl (exon 7, Dharmacon, #VOZOB-000001, UCAAGGAACUCGAUCGUAUUU), ARv7 (Dharmacon, #SRYWD-000001, GUAGUUGUAUCAUGAUU) or both (exon1, Dharmacon, #SRYWD-000005, CAAGGGAGGUUACACCAAAUU) were purchased from Dharmacon.

### ChIP-Sequencing

ChIP-seq was performed as previously described ^27^. The cells were cultured and plated to ensure they reached a healthy confluency of 90% on 10 cm plates before crosslinking. Briefly, cells were crosslinked with 1% formaldehyde (single fixation) or 2 mM fresh disuccinimidyl glutarate and 1% formaldehyde (double fixation) and then chromatin sonicated to 200–600 bp. ChIP was carried out using antibodies against H3K27Ac (Diagenode, C15410196), H3K4me1 (Abcam, ab8895), C-terminal ARfl (Spring Bioscience, clone SP242), ARv7 (RevMAb Biosciences USA Inc., 31-1109-00), FOXA1 (Cell Signaling Technologies, 53528S), BRD4 (Diagenode, C15410337), p300 (Cell Signaling Technologies, 57625S) and then conjugation to Protein A and G Dynabeads (Thermo Fisher Scientific). ChIP DNA was purified using the ChIP DNA Clean & Concentrator^TM^ Kit (Zymo Research, D5205). ChIP-seq libraries were generated using the ThruPLEX DNA-seq Kit (Rubicon Genomics), Accel-NGS 2S Plus DNA Libraries Kit with Unique Dual Indexing (Swift Biosciences, 29096/290384) or SimpleChIP ChIP-seq Multiplex Oligos for Illumina with Dual Index Primers (Cell Signaling Technology, 47538) according to the manufacturer’s instructions. Libraries were pooled and sequenced on the Illumina NextSeq500 platform at the Molecular Biology Core Facility (DFCI) or NovaSeq6000 platform at the Novogene (Sacramento, CA)

### MNase-Sequencing

MNase-seq was performed as previously described ^64^. VCaP and VCaP16 cells were cultured in complete DMEM media supplemented with 10% FBS and 1% Penicillin (5000 IU/ml)-Streptomycin (5000 μg/ml) solution. VCaP16 cells were cultured with the addition of 16 μM ENZ. Twenty-four hours prior to harvesting, fresh culture medium was added. Cells were crosslinked with 1% formaldehyde and then 2.5 × 10^6^ cells were treated with 60 U of MNase (Worthington, LS004797) and Exonuclease III (E. coli) (New England BioLab, M0206S). DNA was purified using ChIP DNA Clean & Concentrator^TM^ Kit (Zymo Research, D5205). Libraries were generated using the SimpleChIP ChIP-seq Multiplex Oligos for Illumina with Dual Index Primers (Cell Signaling Technology, 47538) according to the manufacturer’s instructions. Libraries were pooled and sequenced on the NovaSeq6000 platform at the Novogene (Sacramento, CA).

### ATAC-seq

For bulk ATAC-seq cells were crosslinked with fresh 1% formaldehyde for 10 minutes, then quenched with glycine and frozen. Cells were thawed in cold ATAC-seq buffer (ASB, 10 mM Tris-HCl pH 7.4, 10 mM NaCl, and 3 mM MgCl2) and centrifuged. Cells were lysed by resuspension in .05 mL of lysis buffer (ASB with 0.1% NP40, 0.1% Tween-20, and 0.01% digitonin), followed by addition of 1 mL ASB with 0.1% Tween-20 and centrifugation. Nuclei were resuspended in 50 μL tagmentation reaction containing 25 μL 2X tagmentation buffer (Illumina 15027866), 16.5 μL PBS, 0.5 μL Tween-20, 0.5 μL 1:1 water diluted digitonin, 2.5 μL Tn5 transposase (Illumina 15027865), and 5 μL water, and incubated at 37°C for 30 minutes. Transposed DNA was purified using Qiagen columns. Libraries were amplified and sequencing was performed in thirty-five base pair paired-end reads on a NextSeq instrument (Illumina). For single cell ATAC-seq, approximately 7000 cells were targeted for each sample and processed according to the 10X Genomics scATAC-seq sample preparation protocol (Chromium Single Cell ATAC Library & Gel Bead Kit, 10Å∼ Genomics). Libraries were sequenced as paired-end reads on a NextSeq instrument (Illumina).

### RNA-seq analysis

Total RNA was isolated using the RNeasy Plus Kit (Qiagen) following the manufacturer’s instructions. mRNA libraries were generated using the Illumina TruSeq stranded mRNA sample kit. Raw reads were analyzed using a pipeline for RNA-seq analysis - Visualization Pipeline for RNA-seq analysis (VIPER) based on workflow management system Snakemake (https://github.com/hanfeisun/viper-rnaseq). The read alignment to hg19 reference genome was performed using STAR aligner (2.7.0f) with default parameters. Gene expression (FPKM values) was quantitated with Cufflinks (v2.2.1). Bedtools (v2.27.1) genomecov and bedGraphToBigWig v 4 were used to generate bigwig files.

### ChIP-seq analysis

The data was analyzed using a pipeline CHromatin enrIchment ProcesSor Pipeline (CHIPS) based on workflow management system Snakemake (https://github.com/liulab-dfci/CHIPS) unless otherwise stated. Raw reads were aligned to hg19 reference genome using Burrows-Wheeler Aligner (bwa mem) 0.7.15-r1140 with default parameters; MACS2 algorithm (2.2.7.1) was used to call peaks; MACS2 (2.2.7.1) and bedGraphToBigWig (v 4) were used to generate bigwig files. Read counts for downstream analyses were quantified using the bioliquidator/bamliquidator tool from Docker image (https://hub.docker.com/r/bioliquidator/bamliquidator). Genomic regions were annotated using Hypergeometric Optimization of Motif EnRichment tool (HOMER v3.12). Motif enrichment analysis was performed using HOMER v3.12. Output bam files (from CHIPS pipeline) for ARv7 and ARfl in VCaP and VCaP16 cells were used for secondary peak calling with HOMER v3.12 to generate bed files that were used for further downstream analysis. Bigwig files for samples with biological duplicates were averaged using a combination of the wiggletools and wigToBigWig tools.

### MNase-seq analysis

The raw sequencing data underwent analysis using a modified version of the nf-core/mnaseseq pipeline. Raw reads were mapped to hg19 with Burrows-Wheeler Aligner 0.7.8 (bwa mem) with default parameters. Unmapped reads, not primary reads, reads aligned to different locations and low-quality reads were filtered out with samtools 1.9. Bibwig files were generated with deeptools 3.0.2. Profile plots and heatmaps were generated with deeptools 3.0.2 to visualize the nucleosome footprints.

### WES analysis

Genomic DNA was isolated using DNeasy Blood & Tissue Kit (Qiagen). Libraries for Whole Exome Sequencing (WES) were prepared, and the sequencing was performed on NovaSeq6000 platform at the Novogene (Sacramento, CA) aiming for 100X coverage. The analysis of WES data followed the established Genome Analysis Toolkit (GATK 4.1.9.0) Best Practices Workflows. The raw reads were aligned to the hg38 reference genome using the Burrows-Wheeler Aligner (BWA) version 0.7.8. The identification of candidate somatic variants was carried out using the Mutect2 function from GATK version 4.1.9.0. Identified somatic variants were annotated using Funcotator, which is a part of GATK 4.1.9.0. ANNOVAR (ANNOtate VARiation) tool was utilized for the interpretation and prioritization of the filtered somatic variants. To assess copy ratio alterations, we employed a suite of functions from GATK 4.1.9.0, including PreprocessIntervals, CollectReadCounts, DenoiseReadCounts, and PlotDenoisedCopyRatios, which also allowed for visualization of the data.

### ATAC-seq analysis

The data were analyzed using a pipeline CHromatin enrIchment ProcesSor Pipeline (CHIPS) based on workflow management system Snakemake (https://github.com/liulab-dfci/CHIPS) and Containerized Bioinformatics workflow for Reproducible ChIP/ATAC-seq Analysis (CoBRA) (https://bitbucket.org/cfce/cobra; cfce-cobra.readthedocs.io/en/latest/).

### scATAC-seq analysis

The raw FASTQ files underwent preprocessing using the Cell Ranger ATAC pipeline, version 1.2.0, provided by 10X Genomics (https://support.10xgenomics.com/single-cell-atac/software). The data was aligned to hg19. Dimensionality reduction was achieved using Latent Semantic Analysis (LSA) applied to the filtered peak-barcode matrices. The data was divided into 8 clusters through k-means clustering. Visualization of these clusters was performed using t-distributed stochastic neighbor embedding (t-SNE). For gene annotations, the GENCODE Gene track, version 28lift37, was utilized. For further downstream analysis and visualization of scATAC-Seq data, the Loupe Browser, version 6.5.0, was employed. Signac and Seurat R packages were used to complement the analysis. Latent Semantic Indexing (LSI) dimensionality reduction was applied for the Uniform Manifold Approximation and Projection (UMAP) algorithm. Clusters were defined using the Spectral Latent Manifold (SLM) algorithm. Trajectory Inference: To uncover the global structure of the identified clusters and convert this structure into smooth lineages represented by pseudotime, the Slingshot algorithm (Trajectory Inference for Single-Cell Data) was employed.

## DATA AVAILABILITY

Public datasets used in this publication are available at GEO via the provided accession numbers: AR ChIP-seq data GSE106561 (GSM2842708, GSM2842700, GSM2842704), GSE99378 (GSM2643239, GSM2643240, GSM2643241, GSM2643242, GSM2643245, GSM2643246, GSM2643247, GSM2643248), GSE143906 (GSM4276560, GSM4276561), GSE130408 (GSM4569829 – GSM4569867), GSE39879 (GSM980655, GSM980664), GSE137527 (GSM4081278 – GSM4081325); RNA-seq data GSE126078;expression profiling by array GSE21034 (GSM526290 – GSM5262318, GSM526265 – GSM526283, GSM526134 – GSM526264). Genomic data generated in this study are deposited in GEO under the accession number GSE252902.

## ACKNOWLEDGMENTS

This work was supported by NCI P01 CA163227 (MB, PSN, EC, HL, SPB), NCI P50 CA272390 (MB, SPB), NCI U54 CA156732 (SPB), NCI R01CA272934 (SPB), DoD PCRP Idea Award W81XVVH-20-100925 (SPB), Koch Institute-Dana Farber/Harvard Cancer Center Bridge Project Award (SPB), and a Prostate Cancer Foundation Challenge Award (SPB). JWR was supported by a Prostate Cancer Foundation Young Investigator Award (18YOUN24) and DoD PCRP Early Investigator Award (W81XWH-18-1-0531). AV was supported by a DoD PCRP Physician Research Award (PC200820) and ASCO (Young Investigator Award, 2021A010981). NAL was supported by funding from TUBITAK (114Z491), DoD (W81XWH-21-1-0234) and CIHR (PJT-173331). AGS was supported by the Intramural Research Program of the National Cancer Institute, NIH. MLF was supported by the Claudia Adams Barr Program for Innovative Cancer Research, the Dana-Farber Cancer Institute Presidential Initiatives Fund, the H.L. Snyder Medical Research Foundation, the Cutler Family Fund for Prevention and Early Detection, the Donahue Family Fund, DoD (W81XWH-21-1-0339, W81XWH-22-1-0951), NIH (R01CA251555, R01CA227237, R01CA262577, R01CA259058) and a Movember PCF Challenge Award. We thank Dr. Stephen Plymate for thoughtful input during the course of these studies.

## CONFLICTS OF INTEREST

PSN has received consulting fees from Janssen, Merck and Bristol Myers Squibb and research support from Janssen for work unrelated to the present studies. No conflicts are reported by the other authors.

## Notes

### Competing Interest Statement

The authors have declared no competing interest.

### Summary of Updates

This version has been revised to include further data related to NFI transcription factors.

