## Supplemental Figures for "Increased chromatin accessibility mediated by nuclear factor I drives transition to androgen receptor splice variant dependence in castration-resistant prostate cancer"

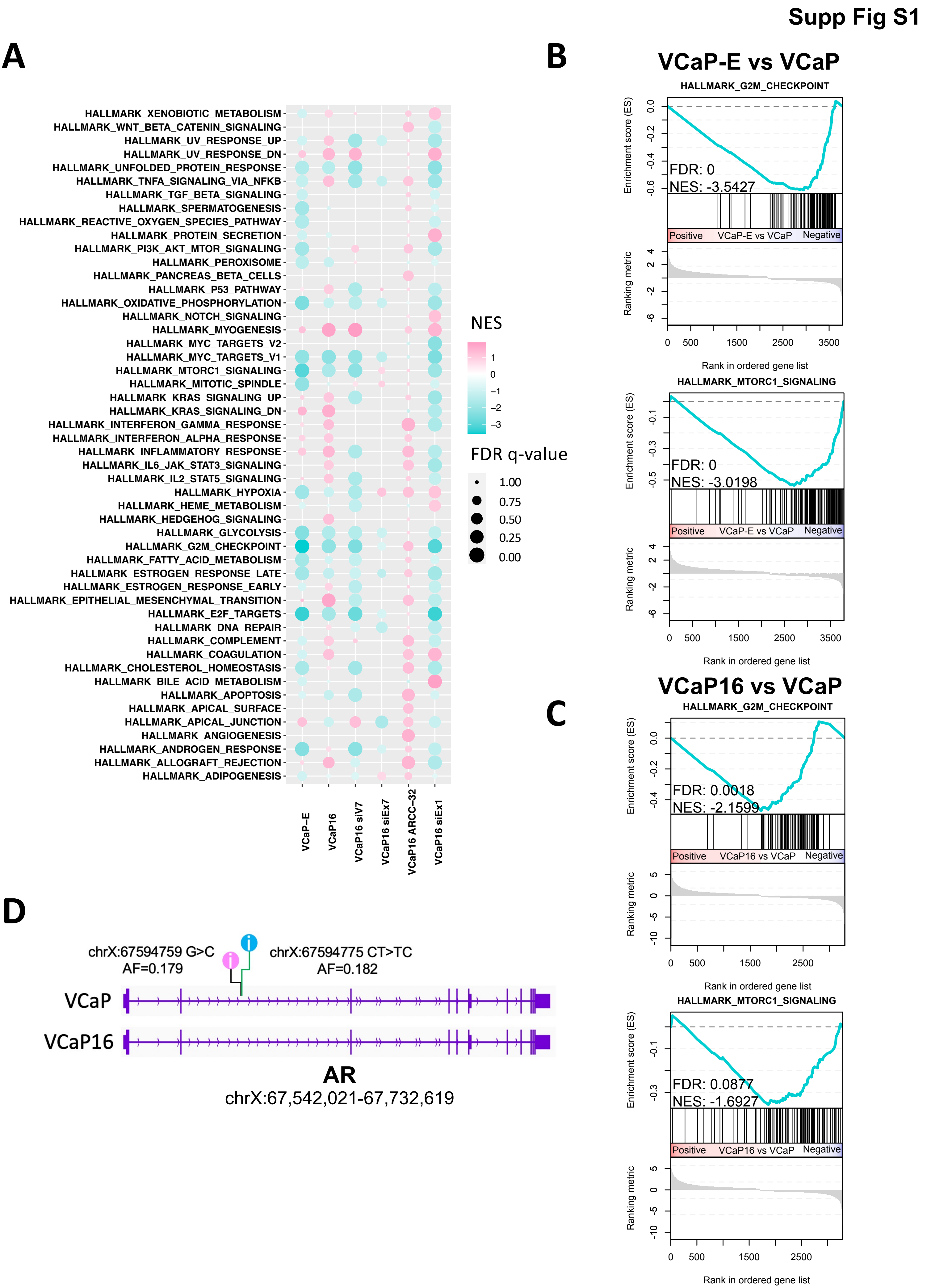

Supp Fig S1
A
B
VCaP-E vs VCaP
HALLMARK_G2M_CHECKPOINT
FDR: 0
NES: -3.5427
HALLMARK_MTORC1_SIGNALING
FDR: 0
NES: -3.0198
Positive
VCaP-E vs VCaP
Negative
Enrichment score (ES)
Ranking metric
Rank in ordered gene list
C
VCaP16 vs VCaP
HALLMARK_G2M_CHECKPOINT
FDR: 0.0018
NES: -2.1599
HALLMARK_MTORC1_SIGNALING
FDR: 0.0877
NES: -1.6927
Positive
VCaP16 vs VCaP
Negative
NES
FDR q-value
1.00
0.75
0.50
0.25
0.00
HALLMARK_XENOBIOTIC_METABOLISM
HALLMARK_WNT_BETA_CATENIN_SIGNALING
HALLMARK_UV_RESPONSE_UP
HALLMARK_UV_RESPONSE_DN
HALLMARK_UNFOLDED_PROTEIN_RESPONSE
HALLMARK_TNFA_SIGNALING_VIA_NFKB
HALLMARK_TGF_BETA_SIGNALING
HALLMARK_SPERMATOGENESIS
HALLMARK_REACTIVE_OXYGEN_SPECIES_PATHWAY
HALLMARK_PROTEIN_SECRETION
HALLMARK_PI3K_AKT_MTOR_SIGNALING
HALLMARK_PEROXISOME
HALLMARK_PANCREAS_BETA_CELLS
HALLMARK_P53_PATHWAY
HALLMARK_OXIDATIVE_PHOSPHORYLATION
HALLMARK_NOTCH_SIGNALING
HALLMARK_MYOGENESIS
HALLMARK_MYC_TARGETS_V2
HALLMARK_MYC_TARGETS_V1
HALLMARK_MTORC1_SIGNALING
HALLMARK_MITOTIC_SPINDLE
HALLMARK_KRAS_SIGNALING_UP
HALLMARK_KRAS_SIGNALING_DN
HALLMARK_INTERFERON_GAMMA_RESPONSE
HALLMARK_INTERFERON_ALPHA_RESPONSE
HALLMARK_INFLAMMATORY_RESPONSE
HALLMARK_IL6_JAK_STAT3_SIGNALING
HALLMARK_IL2_STAT5_SIGNALING
HALLMARK_HYPOXIA
HALLMARK_HEME_METABOLISM
HALLMARK_HEDGEHOG_SIGNALING
HALLMARK_GLYCOLYSIS
HALLMARK_G2M_CHECKPOINT
HALLMARK_FATTY_ACID_METABOLISM
HALLMARK_ESTROGEN_RESPONSE_LATE
HALLMARK_ESTROGEN_RESPONSE_EARLY
HALLMARK_EPITHELIAL_MESENCHYMAL_TRANSITION
HALLMARK_E2F_TARGETS
HALLMARK_DNA_REPAIR
HALLMARK_COMPLEMENT
HALLMARK_COAGULATION
HALLMARK_CHOLESTEROL_HOMEOSTASIS
HALLMARK_BILE_ACID_METABOLISM
HALLMARK_APOPTOSIS
HALLMARK_APICAL_SURFACE
HALLMARK_APICAL_JUNCTION
HALLMARK_ANGIOGENESIS
HALLMARK_ANDROGEN_RESPONSE
HALLMARK_ALLOGRAFT_REJECTION
HALLMARK_ADIPOGENESIS
VCaP-E
VCaP16
VCaP16 siV7
VCaP16 siEx7
VCaP16 ARCC-32
VCaP16 siEx1
D
chrX:67594759 G>C
AF=0.179
chrX:67594775 CT>TC
AF=0.182
VCaP
VCaP16
AR
chrX:67,542,021-67,732,619

**Supplementary Figure S1. AR gene expression in VCaP16 cells depends on ARv7. A)** Gene set enrichment analysis (GSEA) of RNA-seq data from VCaP cells treated with ENZ for 4 days (VCaP-E), VCaP16 cells (maintained in 16 μM ENZ), VCaP16 cells with siRNA knockdown of ARv7 (siV7), VCaP16 cells with siRNA knockdown of ARfl (siEx7), VCaP16 cells treated with ARCC-32 for 36 hrs, and VCaP16 cells with siRNA knockdown of both ARv7 and ARfl (siEx1). **B)** Enrichment plots of Hallmark G2M Checkpoint and MTORC1 Signaling by GSEA of RNA-seq data from VCaP cells treated with ENZ for 4 days (VCaP-E) and **C)** VCaP16 cells versus parental VCaP. Significantly differentially expressed genes with padj < 0.05 (VCaP-E, VCaP16, VCaP16 siV7, VCaP siEx1, VCaP siEx7) or all genes (VCaP16 ARCC-32) were used as an input. **D)** Whole exome sequencing analysis of the *AR* gene in VCaP16 and parental VCaP showing two intronic variants picked up in both, but the absence of novel exonic alterations in VCaP16 cells.

### Whole Exome Sequencing

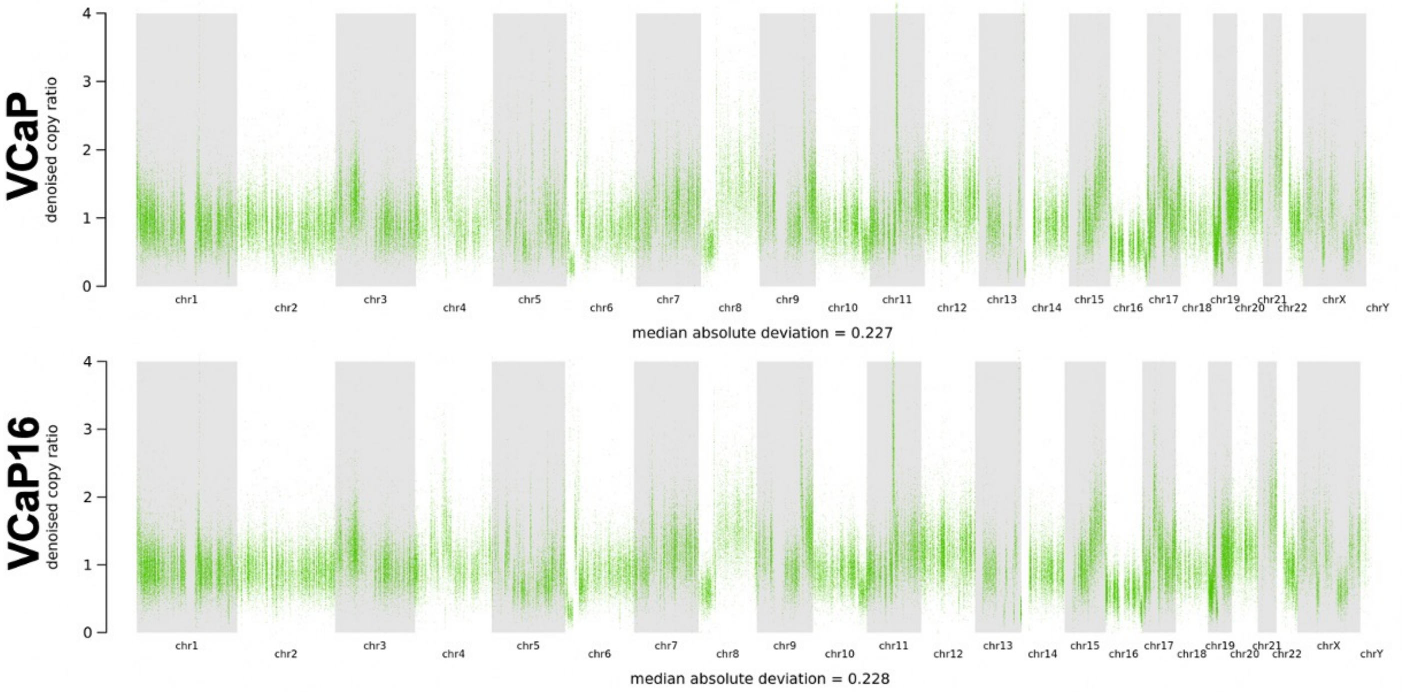

**Supplementary Figure S2. Copy number alterations in VCaP versus VCaP16.** Whole exome sequencing was done on VCaP16 and parental VCaP and used to assess for copy number alterations in VCaP16 and parental VCaP cells. No additional losses or gains were found in VCaP16 versus VCaP.

**Supp Fig S3**

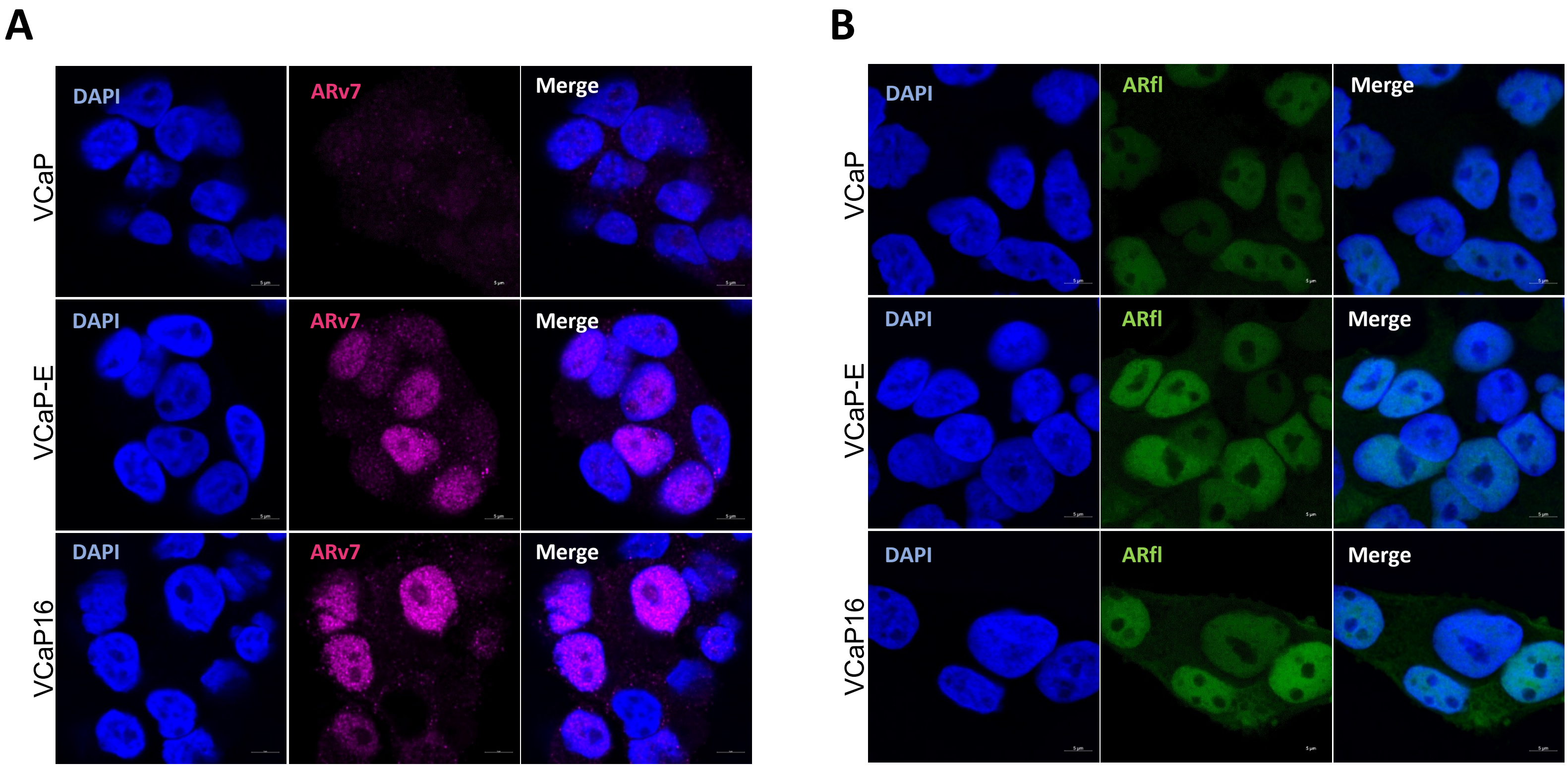

**Supplementary Figure S3. Nuclear localization of ARv7 and ARfl.** Immunofluorescence was assessed in VCaP16, VCaP, and VCaP cells treated with ENZ (16 μM for 4 days) (VCaP-E). Cells were stained with an anti-ARv7 Ab (**A**) or anti-ARfl Ab directed at the C-terminal of the LBD (**B**).

### Supp Fig S4

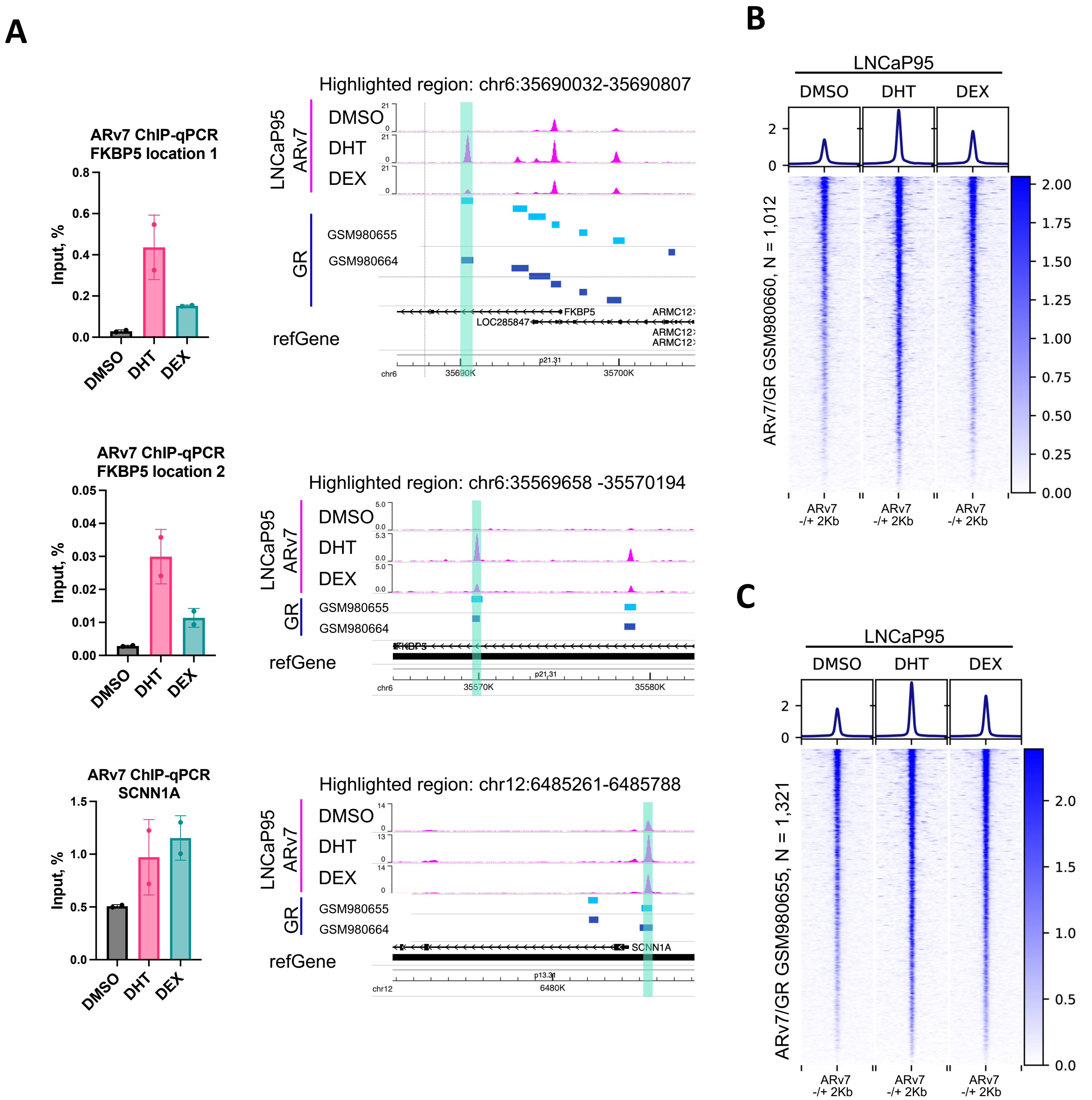
A
ARv7 ChIP-qPCR
FKBP5 location 1
Input, %
DMSO
DHT
DEX
Highlighted region: chr6:35690032-35690807
LNCaP95
ARv7
GR
GSM980655
GSM980664
refGene
FKBP5
LOC285847
ARMC12
chr6
35690K
35700K
p21.31
ARv7 ChIP-qPCR
FKBP5 location 2
Highlighted region: chr6:35569658 -35570194
35570K
35580K
ARv7 ChIP-qPCR
SCNN1A
Highlighted region: chr12:6485261-6485788
SCNN1A
chr12
6480K
p13.31
B
LNCaP95
ARv7/GR GSM980660, N = 1,012
ARv7
-/+ 2Kb
C
ARv7/GR GSM980655, N = 1,321

**Supplementary Figure S4. Assisted loading of ARv7 onto chromatin. A)** Enrichment of ARv7 binding at indicated sites by ChIP-qPCR (enrichment versus input, %) and screenshots of ARv7 binding intensities by ChIP-seq at known ARv7-regulated genes (FKBP5 and SCNN1A) in LNCaP95 cells (maintained in ENZ) treated with DMSO, 10 nM DHT (DHT), or 100 nM dexamethasone (DEX) for 2 hrs. Glucocorticoid receptor (GR) binding sites in VCaP cells stimulated with dexamethasone (DEX) are shown in sky-blue (GSM980665) and blue (GSM980664). **B, C)** Heatmaps showing the enrichment of ARv7 binding in LNCaP95 cells treated with DMSO (DMSO), 10 nM DHT (DHT) or 100 nM dexamethasone (DEX) for 2 hrs. ARv7 ChIP-seq signal is centered at intersection of ARv7 and GR binding sites from GSM980660 (B) and GSM980665 (C).

**Supp Fig S5**

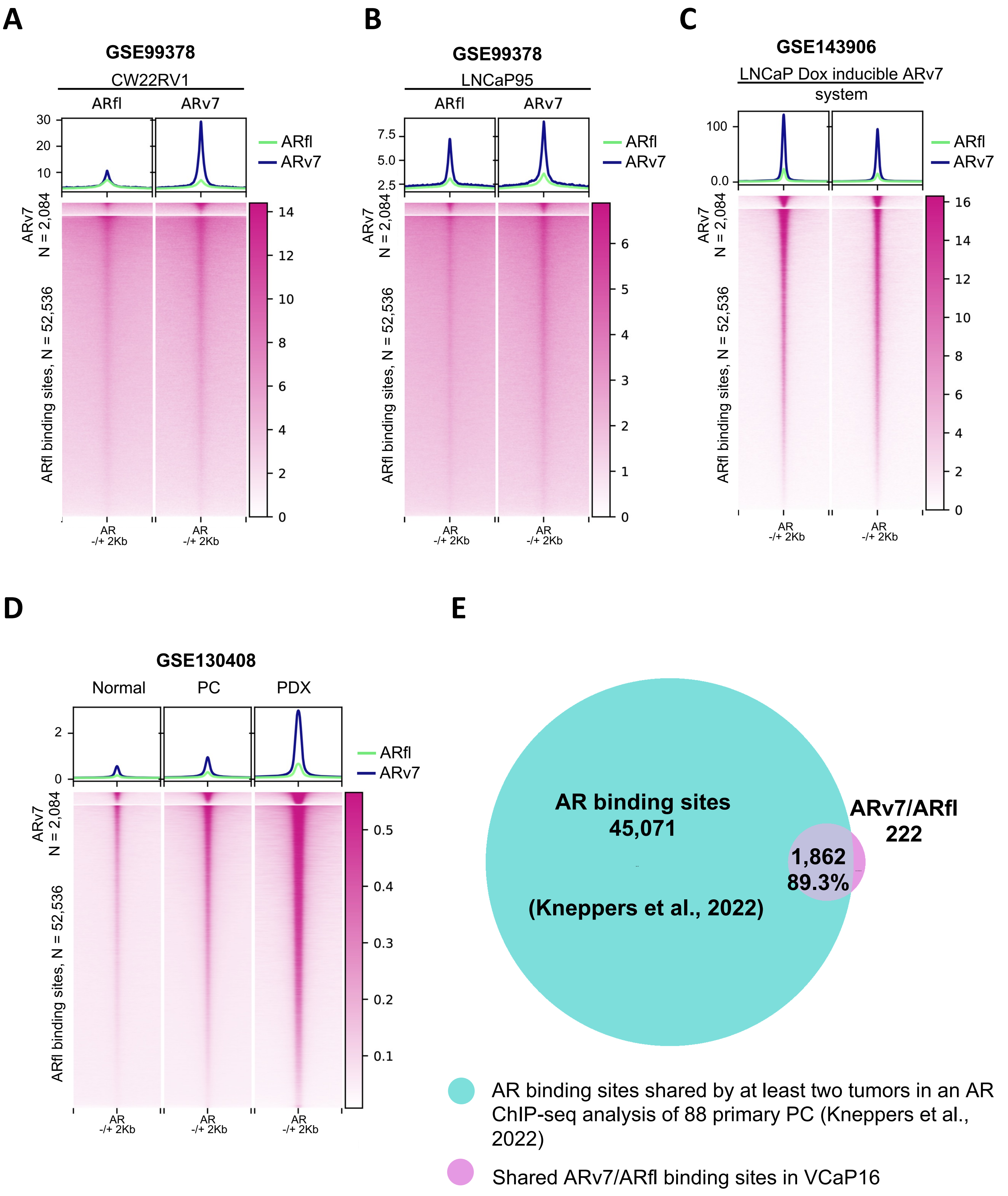
A
GSE99378
CW22RV1
ARfl
ARv7
ARfl
ARv7
ARv7 N = 2,084
ARfl binding sites, N = 52,536
AR -/+ 2Kb
B
GSE99378
LNCaP95
ARfl
ARv7
ARfl
ARv7
ARv7 N = 2,084
ARfl binding sites, N = 52,536
AR -/+ 2Kb
C
GSE143906
LNCaP Dox inducible ARv7 system
ARfl
ARv7
ARv7 N = 2,084
ARfl binding sites, N = 52,536
AR -/+ 2Kb
D
GSE130408
Normal
PC
PDX
ARfl
ARv7
ARv7 N = 2,084
ARfl binding sites, N = 52,536
AR -/+ 2Kb
E
AR binding sites
45,071
(Kneppers et al., 2022)
ARv7/ARfl
222
1,862
89.3%
AR binding sites shared by at least two tumors in an AR ChIP-seq analysis of 88 primary PC (Kneppers et al., 2022)
Shared ARv7/ARfl binding sites in VCaP16

**Supplementary Figure S5. ARv7 binding sites identified in VCaP16 are shared in other cell lines and are high affinity ARfl sites in cell lines and in clinical samples. A)** Heatmap showing the ARfl (left) and ARv7 (right) signal intensities in CW22RV1 (GSE99378), **B)** LNCaP95 (GSE99378), and **C)** LNCaP cells with inducible ARv7 (GSE143906). ChIP-seq signal is centered at shared ARfl/ARv7 (ARv7) sites (green) or ARfl unique sites (blue) identified in VCaP16 cells**. D)** Heatmap showing enrichment of ARfl binding to chromatin in normal prostate epithelium (Normal), primary prostate tumors (PC) and metastatic prostate cancer specimens (GSE130408). Multiple bigwig files are averaged per each condition. ChIP-seq signal is centered at shared ARfl/ARv7 (ARv7) sites (green) and ARfl unique sites (blue) identified in VCaP16 cells**. E)** Overlap between ARv7 sites in VCaP and ARfl sites by at least two tumors in an ARfl ChIP-seq study of 88 primary tumors.

**Supp Fig S6**

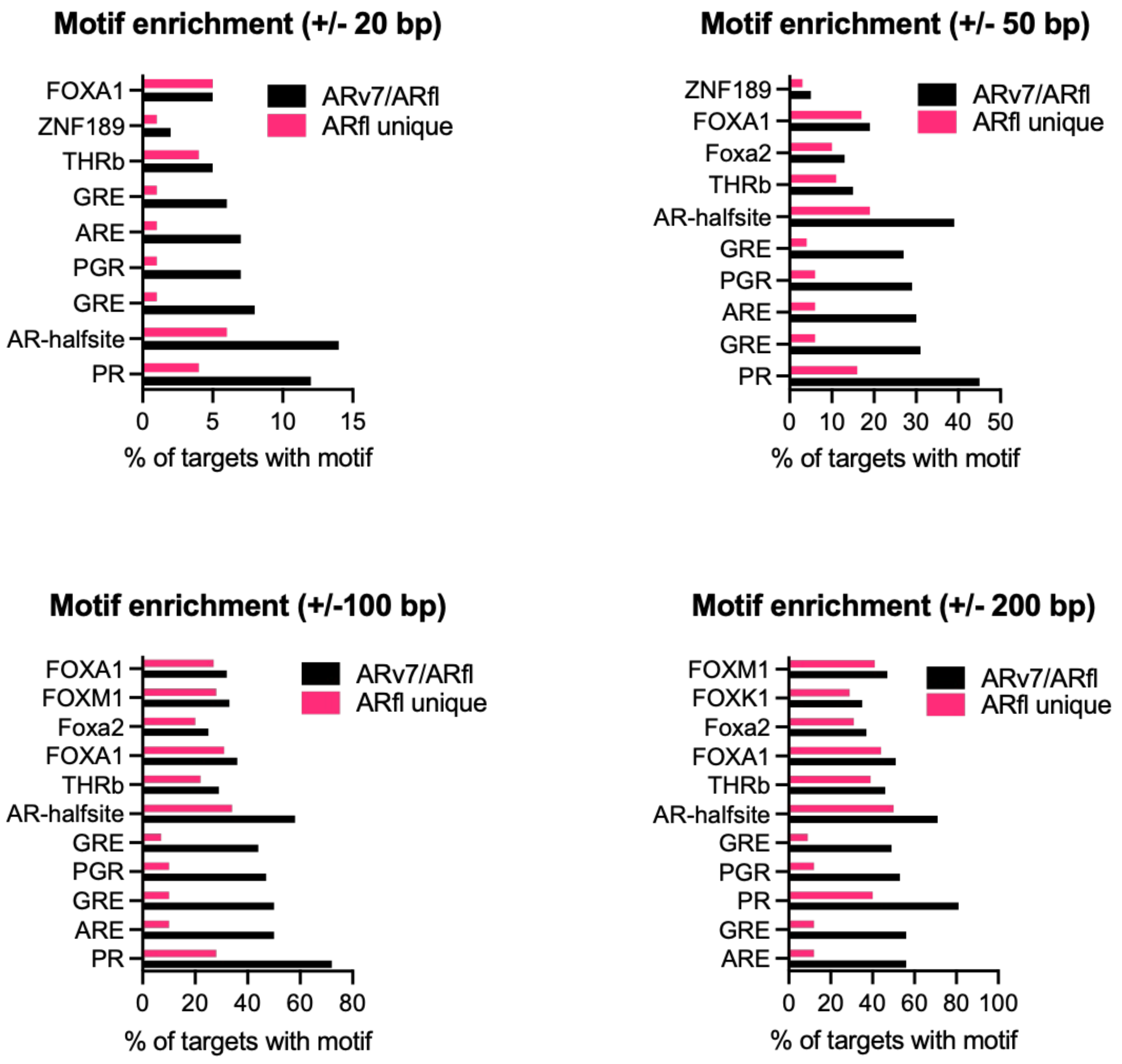
Motif enrichment (+/- 20 bp)
FOXA1
ZNF189
THRb
GRE
ARE
PGR
GRE
AR-halfsite
PR
ARv7/ARfl
ARfl unique
0 5 10 15
% of targets with motif
Motif enrichment (+/- 50 bp)
ZNF189
FOXA1
Foxa2
THRb
AR-halfsite
GRE
PGR
ARE
GRE
PR
ARv7/ARfl
ARfl unique
0 10 20 30 40 50
% of targets with motif
Motif enrichment (+/-100 bp)
FOXA1
FOXM1
Foxa2
FOXA1
THRb
AR-halfsite
GRE
PGR
GRE
ARE
PR
ARv7/ARfl
ARfl unique
0 20 40 60 80
% of targets with motif
Motif enrichment (+/- 200 bp)
FOXM1
FOXK1
Foxa2
FOXA1
THRb
AR-halfsite
GRE
PGR
PR
GRE
ARE
ARv7/ARfl
ARfl unique
0 20 40 60 80 100
% of targets with motif

**Supplementary Figure S6. Motif analysis of ARv7/ARfl binding sites and ARfl unique binding sites in VCaP16 cells.** Motifs enriched within 20, 50 100, or 200 bp from the center of an ARv7/ARfl or ARfl unique site are shown.

Supp Fig S7

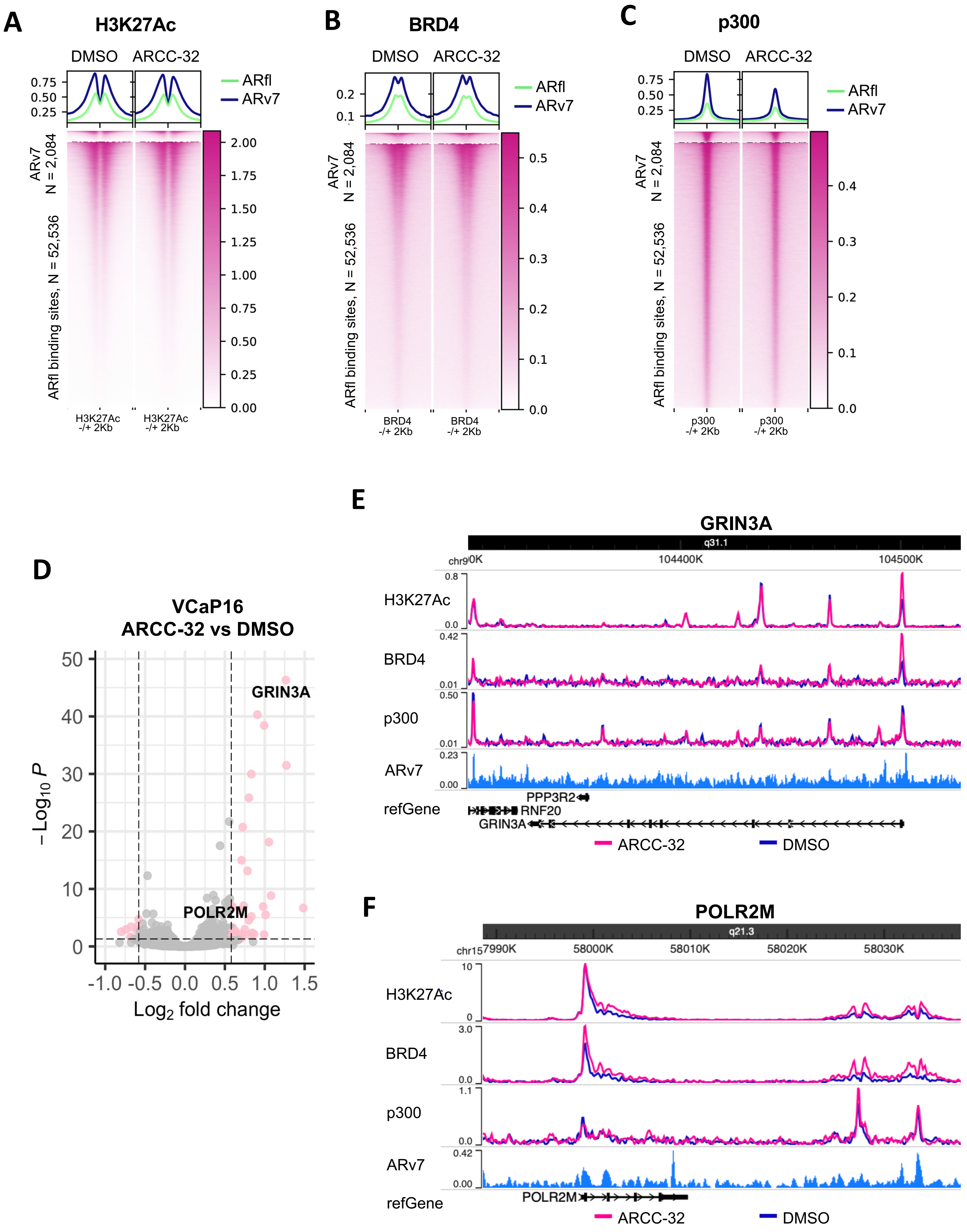
A
H3K27Ac
DMSO
ARCC-32
ARfl
ARv7
ARv7 N = 2,084
ARfl binding sites, N = 52,536
H3K27Ac -/+ 2Kb
B
BRD4
BRD4 -/+ 2Kb
C
p300
p300 -/+ 2Kb
D
VCaP16
ARCC-32 vs DMSO
GRIN3A
POLR2M
–Log10 P
Log2 fold change
E
GRIN3A
q31.1
chr9
104400K
104500K
H3K27Ac
BRD4
p300
ARv7
refGene
PPP3R2
RNF20
GRIN3A
ARCC-32
DMSO
F
POLR2M
q21.3
chr15
57990K
58000K
58010K
58020K
58030K
H3K27Ac
BRD4
p300
ARv7
refGene
POLR2M
ARCC-32
DMSO

**Supplementary Figure S7. Degradation of ARfl increases H3K27Ac and BRD4 binding at a subset of sites. A)** Levels of H3K27Ac, **B)** BRD4 and **C)** p300 by ChIP-seq in VCaP16 cells (maintained in basal medium with 16 μM ENZ) treated with DMSO (DMSO) or 500 nM ARCC-32 (ARCC-32) for 24 hrs. ChIP-seq signal is centered at shared ARfl/ARv7 (ARv7) sites and ARfl unique sites identified in VCaP16 cells**. D)** Volcano plot of differentially expressed genes from RNA-seq data in VCaP16 treated with 500 nM ARCC-32 for 24 hrs versus VCaP16. **E)** and **F)** Screenshots of differential enrichment of H3K27Ac, BRD4 and p300 on chromatin by ChIP-seq (from A, B and C) in response to ARCC-32 treatment at the *GRIN3A* and *POLR2M* genes.

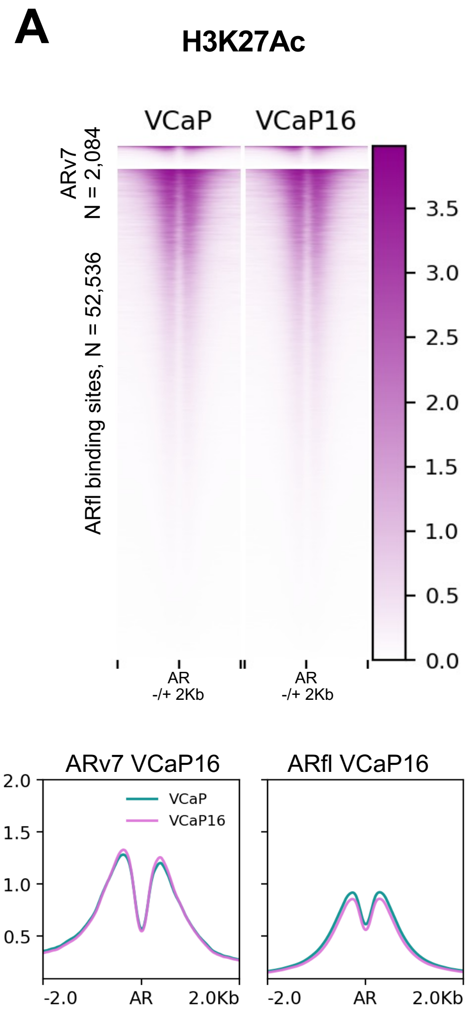
A
H3K27Ac
VCaP
VCaP16
ARv7
N = 2,084
ARfl binding sites, N = 52,536
3.5
3.0
2.5
2.0
1.5
1.0
0.5
0.0
AR
-/+ 2Kb
ARv7 VCaP16
ARfl VCaP16
VCaP
VCaP16
2.0
1.5
1.0
0.5
-2.0
AR
2.0Kb

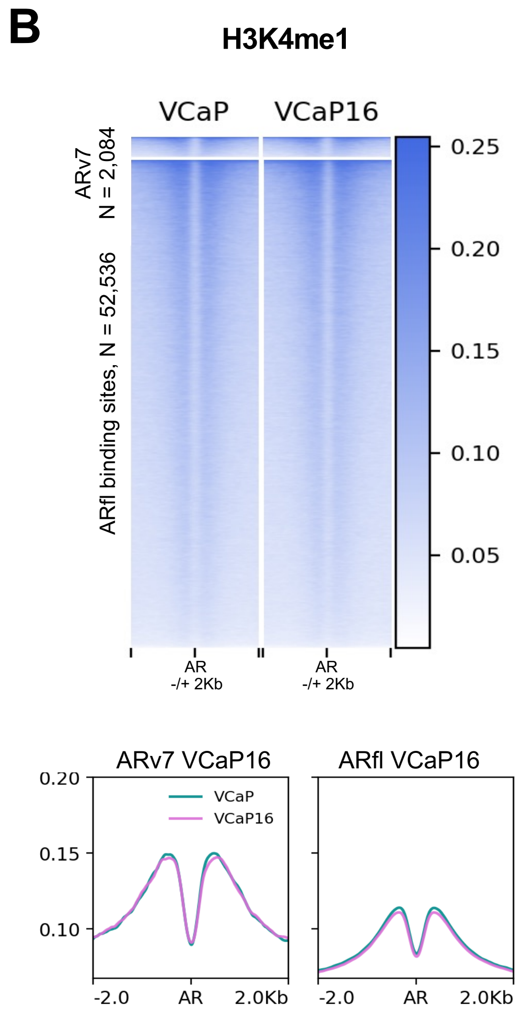
B
H3K4me1
VCaP
VCaP16
ARv7
N = 2,084
ARfl binding sites, N = 52,536
0.25
0.20
0.15
0.10
0.05
AR
-/+ 2Kb
ARv7 VCaP16
ARfl VCaP16
VCaP
VCaP16
0.20
0.15
0.10
-2.0
AR
2.0Kb

**Supplementary Figure S8. Overlap of active enhancer marks and ARv7 in VCaP16 cells. A)** Heatmaps for H3K27Ac and **B)** H3K4me1 at ARv7 and ARfl unique sites in VCaP and VCaP16.

Supp Fig S9

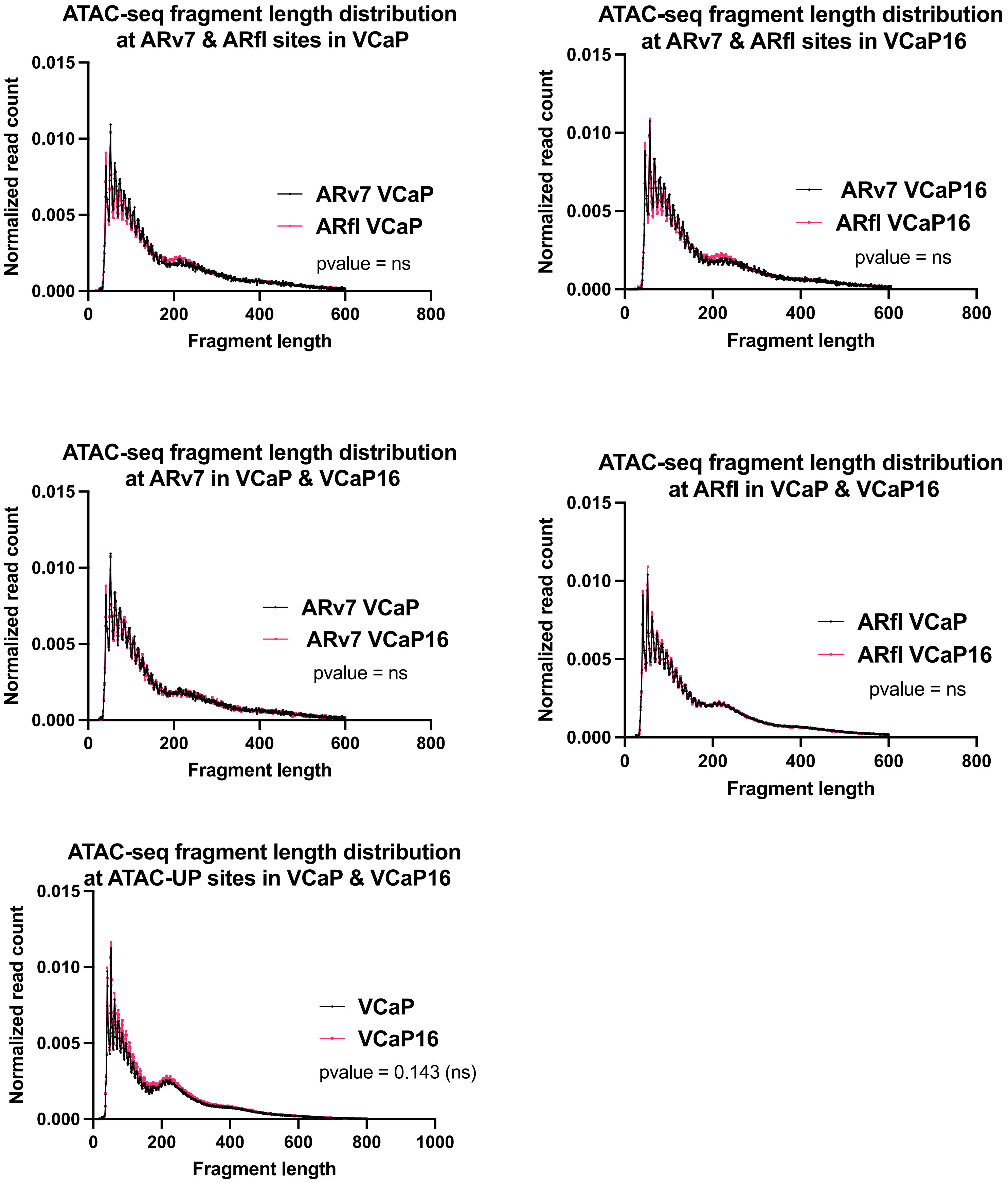

ATAC-seq fragment length distribution at ARv7 & ARfl sites in VCaP
Normalized read count
0.015
0.010
0.005
0.000
0
200
400
600
800
Fragment length
ARv7 VCaP
ARfl VCaP
pvalue = ns
ATAC-seq fragment length distribution at ARv7 & ARfl sites in VCaP16
Normalized read count
0.015
0.010
0.005
0.000
0
200
400
600
800
Fragment length
ARv7 VCaP16
ARfl VCaP16
pvalue = ns
ATAC-seq fragment length distribution at ARv7 in VCaP & VCaP16
Normalized read count
0.015
0.010
0.005
0.000
0
200
400
600
800
Fragment length
ARv7 VCaP
ARv7 VCaP16
pvalue = ns
ATAC-seq fragment length distribution at ARfl in VCaP & VCaP16
Normalized read count
0.015
0.010
0.005
0.000
0
200
400
600
800
Fragment length
ARfl VCaP
ARfl VCaP16
pvalue = ns
ATAC-seq fragment length distribution at ATAC-UP sites in VCaP & VCaP16
Normalized read count
0.015
0.010
0.005
0.000
0
200
400
600
800
1000
Fragment length
VCaP
VCaP16
pvalue = 0.143 (ns)

**Supplementary Figure S9. ATAC-seq fragment length distribution.** Distribution of ATAC-seq fragment lengths containing ARv7 or ARfl binding sites in VCaP and VCaP16 cells.

**Supp Fig S10**

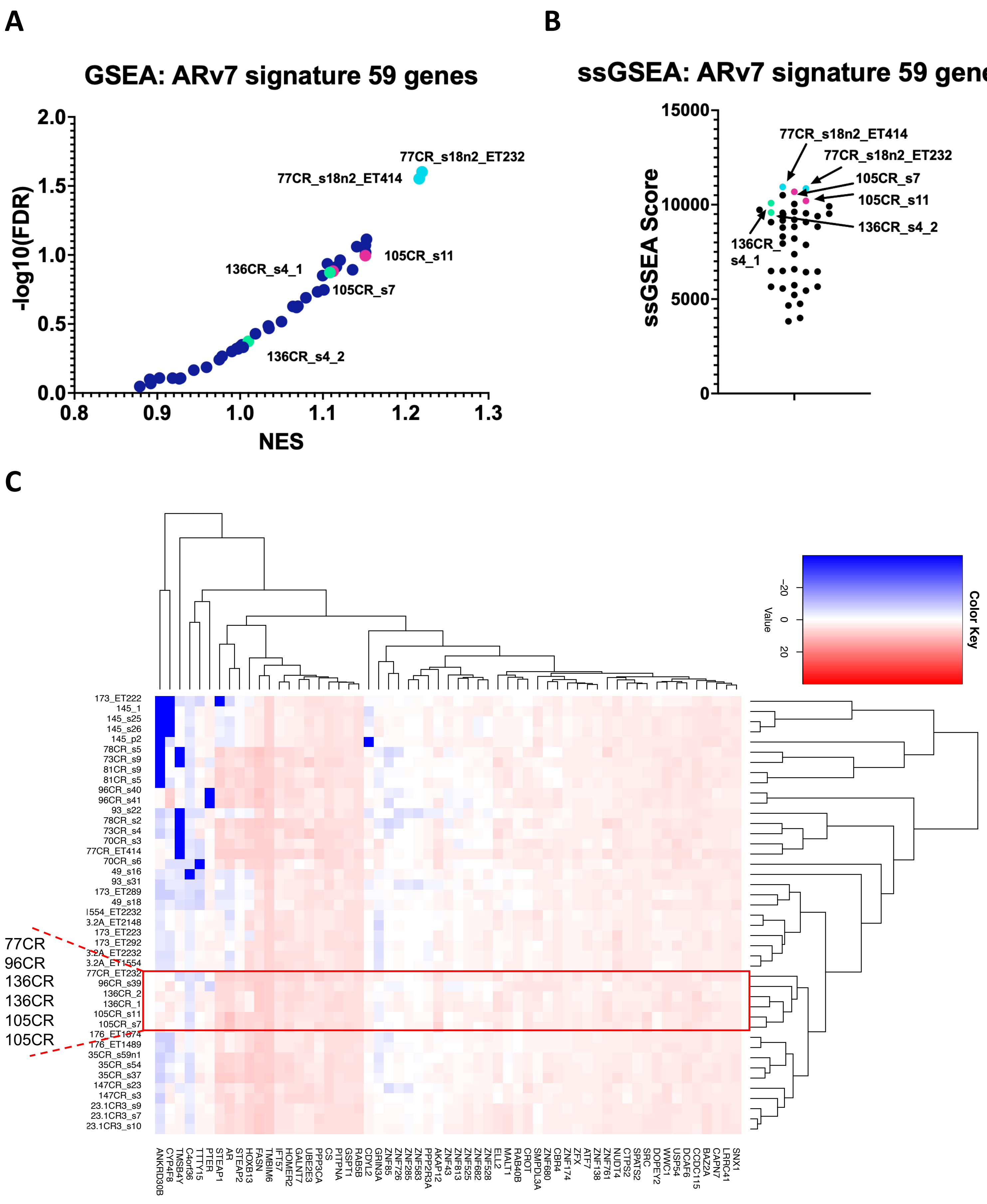

A
GSEA: ARv7 signature 59 genes
-log10(FDR)
NES
0.0
0.5
1.0
1.5
2.0
0.8
0.9
1.0
1.1
1.2
1.3
77CR_s18n2_ET232
77CR_s18n2_ET414
105CR_s11
136CR_s4_1
105CR_s7
136CR_s4_2
B
ssGSEA: ARv7 signature 59 genes
ssGSEA Score
0
5000
10000
15000
77CR_s18n2_ET414
77CR_s18n2_ET232
105CR_s7
105CR_s11
136CR_s4_2
136CR_
s4_1
C
Color Key
Value
−20
0
20
173_ET222
145_1
145_s25
145_s26
145_p2
78CR_s5
73CR_s9
81CR_s9
81CR_s5
96CR_s40
96CR_s41
93_s22
78CR_s2
73CR_s4
70CR_s3
77CR_ET414
70CR_s6
49_s16
93_s31
173_ET289
49_s18
1554_ET2232
3.2A_ET2148
173_ET223
173_ET292
3.2A_ET2232
3.2A_ET1554
77CR_ET232
96CR_s39
136CR_2
136CR_1
105CR_s11
105CR_s7
176_ET1074
176_ET1489
35CR_s59n1
35CR_s54
35CR_s37
147CR_s23
147CR_s3
23.1CR3_s9
23.1CR3_s7
23.1CR3_s10
77CR
96CR
136CR
136CR
105CR
105CR
ANKRD30B
CYP4F8
TMSB4Y
C4orf36
TTTY15
PTER
STEAP1
AR
STEAP2
HOXB13
FASN
TMBIM6
IFT57
HOMER2
GALNT7
UBE2E3
PPP3CA
CS
PITPNA
GSPT1
RAB5B
CDYL2
GRIN3A
ZNF85
ZNF726
ZNF285
ZNF583
PPP2R3A
AKAP12
ZNF43
ZNF813
ZNF525
ZNF682
ZNF528
ELL2
MALT1
RAB40B
CROT
SMPDL3A
ZNF680
CBR4
ZNF174
ZFX
ATF7
ZNF138
ZNF761
NUDT4
CTPS2
SPATS2
SRC
DOPEY2
WWC1
USP54
DCAF6
CCDC115
BAZ2A
CAPN7
LRRC41
SNX1

**Supplementary Figure S10. Enrichment of ARv7 signature genes in PDX models of PC. A)** Gene Set Enrichment Analysis (GSEA) and **B)** single sample GSEA (ssGSEA) of the 59 genes that make up the ARv7 gene set from Sharp et al. in PDX models of PC. **C)** Unsupervised clustering of RNA-seq data (FPKM values) based on the 59-gene set associated with ARv7 from Sharp et al. in PDX models.

**Supp Fig S11**

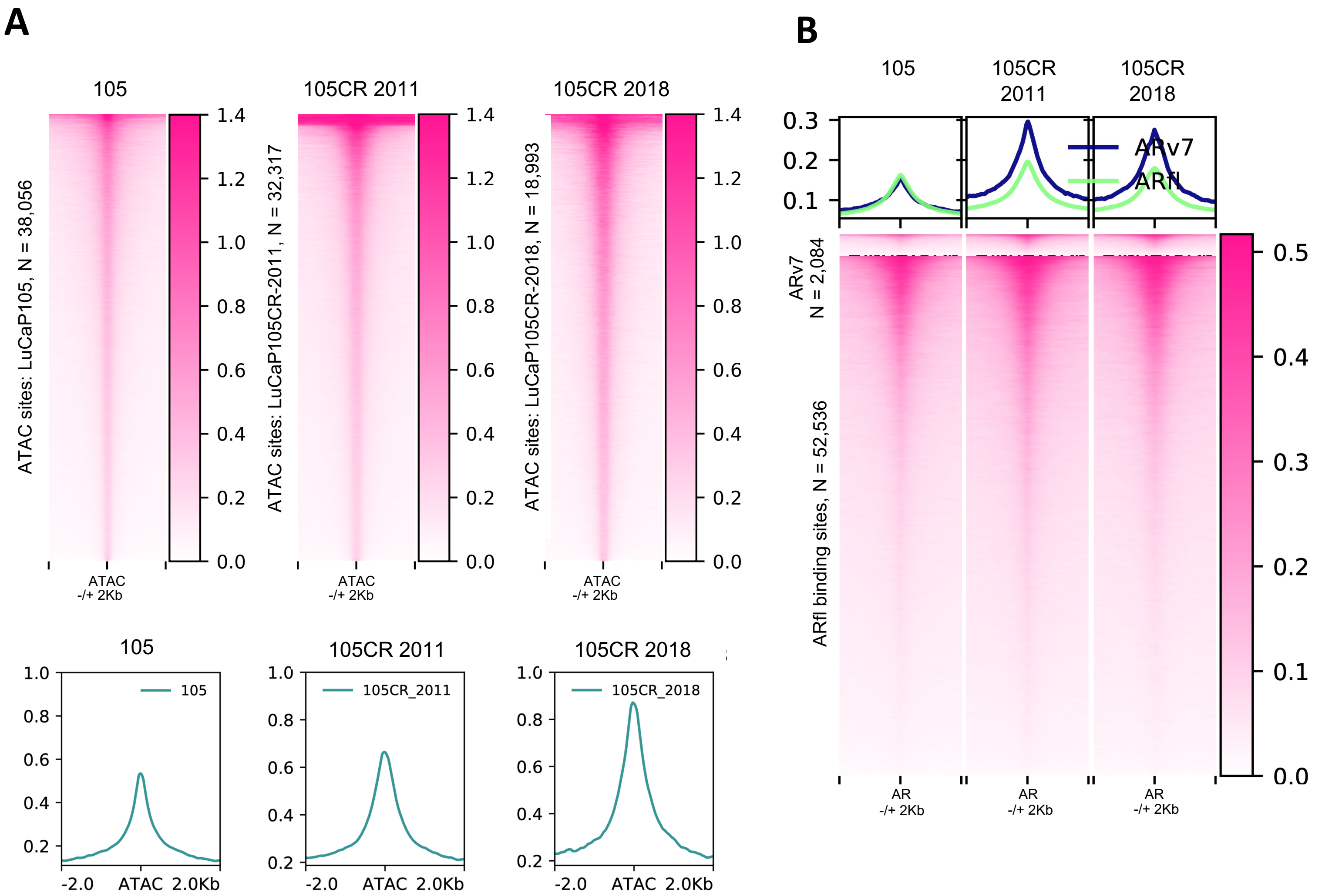

A
105
105CR 2011
105CR 2018
ATAC sites: LuCaP105, N = 38,056
ATAC sites: LuCaP105CR-2011, N = 32,317
ATAC sites: LuCaP105CR-2018, N = 18,993
ATAC
-/+ 2Kb
105
105CR_2011
105CR_2018
-2.0
ATAC
2.0Kb
B
105
105CR 2011
105CR 2018
ARv7
ARfl
ARv7 N = 2,084
ARfl binding sites, N = 52,536
AR
-/+ 2Kb

**Supplementary Figure S11. ATAC-seq in a matched pair of PDX models, LuCaP105 and LuCaP105CR. A)** Enrichment of ATAC-seq signal in PDX models LuCaP105 and LuCaP105CR (LuCaP105CR-2011 and LuCaP105CR-2018). ATAC-seq signal is centered at chromatin accessible sites identified for each sample by ATAC-seq. **B)** Heatmap showing different chromatin accessibility in a pair of PDX models LuCaP105 and LuCaP105CR at ARv7 vs ARfl binding sites. ATAC-seq signal is centered at shared ARfl/ARv7 (ARv7) sites and ARfl unique sites identified in VCaP16 cells**.**

**Supp Fig S12**

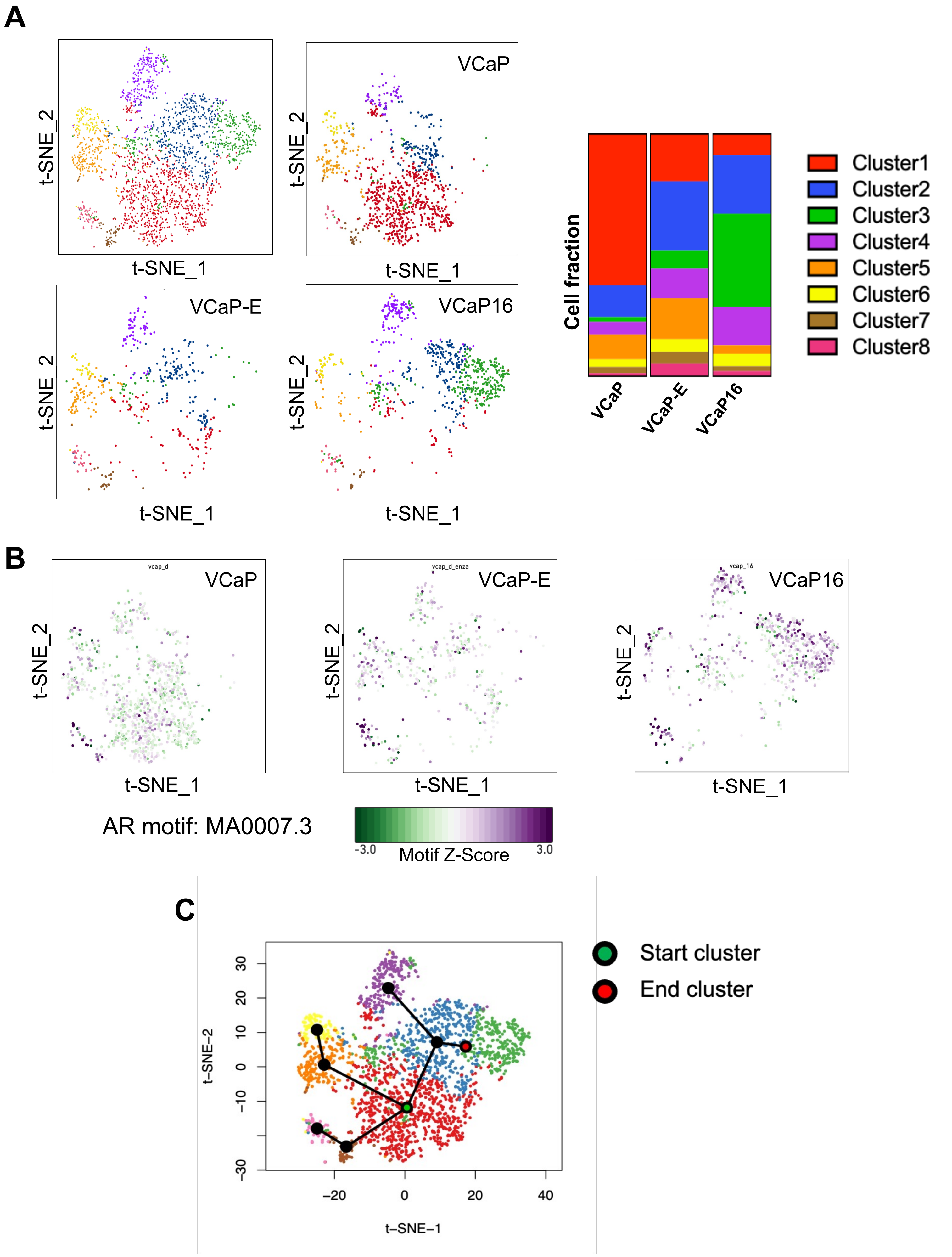

A
t-SNE_2
t-SNE_1
VCaP
VCaP-E
VCaP16
Cell fraction
VCaP
VCaP-E
VCaP16
Cluster1
Cluster2
Cluster3
Cluster4
Cluster5
Cluster6
Cluster7
Cluster8
B
vcap_d
vcap_d_enza
vcap_16
VCaP
VCaP-E
VCaP16
t-SNE_2
t-SNE_1
AR motif: MA0007.3
-3.0
Motif Z-Score
3.0
C
Start cluster
End cluster
t-SNE-2
t-SNE-1
30
20
10
0
−10
−30
−20
0
20
40

**Supplementary Figure S12. Subpopulations identified by scATAC-seq. A)** K-means clustering on the combined scATAC-seq data from VCaP, VCaP treated with ENZ for 96 hrs (VCaP-E), and VCaP16 (maintained in basal medium with 16 μM ENZ). Projection on t-SNE plots. Bar plot showing the distribution of cell fractions across identified clusters in VCaP, VCaP-E, and VCaP16. **B)** AR-motif (MA0007.3) enrichment. **C)** Pseudotime progression by trajectory inference analysis applied to the identified clusters in *A)*.

### Supp Fig S13

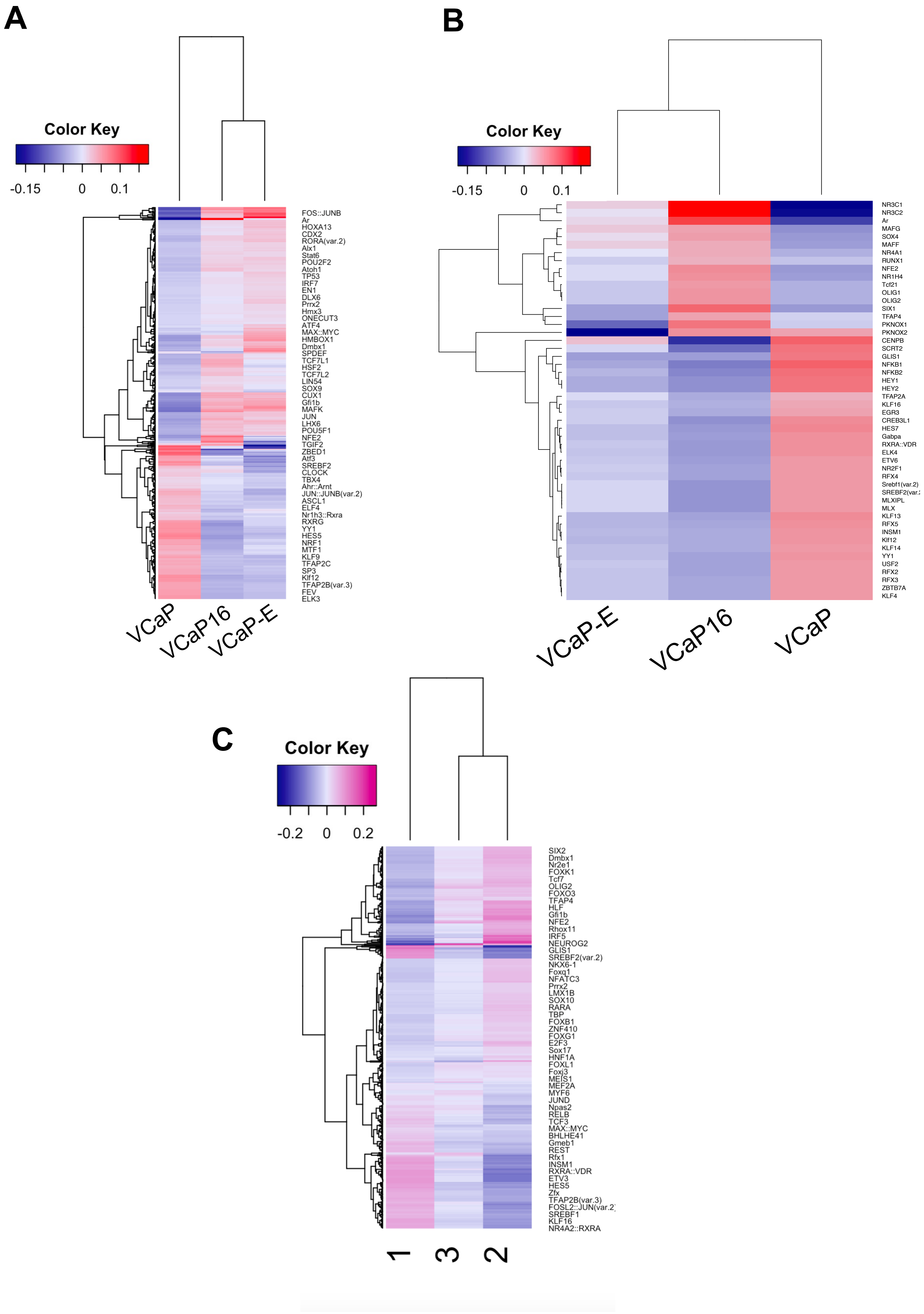

A
Color Key
-0.15
0
0.1
FOS::JUNB
Ar
HOXA13
CDX2
RORA(var.2)
Alx1
Stat6
POU2F2
Atoh1
TP53
IRF7
EN1
DLX6
Prrx2
Hmx3
ONECUT3
ATF4
MAX::MYC
HMBOX1
Dmbx1
SPDEF
TCF7L1
HSF2
TCF7L2
LIN54
SOX9
CUX1
Gfi1b
MAFK
JUN
LHX6
POU5F1
NFE2
TGIF2
ZBED1
Atf3
SREBF2
CLOCK
TBX4
Ahr::Arnt
JUN::JUNB(var.2)
ASCL1
ELF4
Nr1h3::Rxra
RXRG
YY1
HES5
NRF1
MTF1
KLF9
TFAP2C
SP3
Klf12
TFAP2B(var.3)
FEV
ELK3
VCaP
VCaP16
VCaP-E
B
Color Key
-0.15
0
0.1
NR3C1
NR3C2
Ar
MAFG
SOX4
MAFF
NR4A1
RUNX1
NFE2
NR1H4
Tcf21
OLIG1
OLIG2
SIX1
TFAP4
PKNOX1
PKNOX2
CENPB
SCRT2
GLIS1
NFKB1
NFKB2
HEY1
HEY2
TFAP2A
KLF16
EGR3
CREB3L1
HES7
Gabpa
RXRA::VDR
ELK4
ETV6
NR2F1
RFX4
Srebf1(var.2)
SREBF2(var.2)
MLXIPL
MLX
KLF13
RFX5
INSM1
Klf12
KLF14
YY1
USF2
RFX2
RFX3
ZBTB7A
KLF4
VCaP-E
VCaP16
VCaP
C
Color Key
-0.2
0
0.2
SIX2
Dmbx1
Nr2e1
FOXK1
Tcf7
OLIG2
FOXO3
TFAP4
HLF
Gfi1b
NFE2
Rhox11
IRF5
NEUROG2
GLIS1
SREBF2(var.2)
NKX6-1
Foxq1
NFATC3
Prrx2
LMX1B
SOX10
RARA
TBP
FOXB1
ZNF410
FOXG1
E2F3
Sox17
HNF1A
FOXL1
Foxj3
MEIS1
MEF2A
MYF6
JUND
Npas2
RELB
TCF3
MAX::MYC
BHLHE41
Gmeb1
REST
Rfx1
INSM1
RXRA::VDR
ETV3
HES5
Zfx
TFAP2B(var.3)
FOSL2::JUN(var.2)
SREBF1
KLF16
NR4A2::RXRA
1
3
2

**Supplementary Figure S13. Motifs enriched in accessible chromatin. A)** Heatmap showing differential motif activities between VCaP, VCaP treated with ENZ for 96 hrs (VCaP-E) and VCaP16 cells (maintained in basal medium with 16 μM ENZ). Top 50 features are shown in **B)**. **C)** Heatmap showing differential motif activities between cluster 1 (relatively specific to parental VCaP cells), cluster 2 (intermediate cluster observed across all cell lines), and cluster 3 (relatively specific to VCaP16).

**Supp Fig S14**

**A**

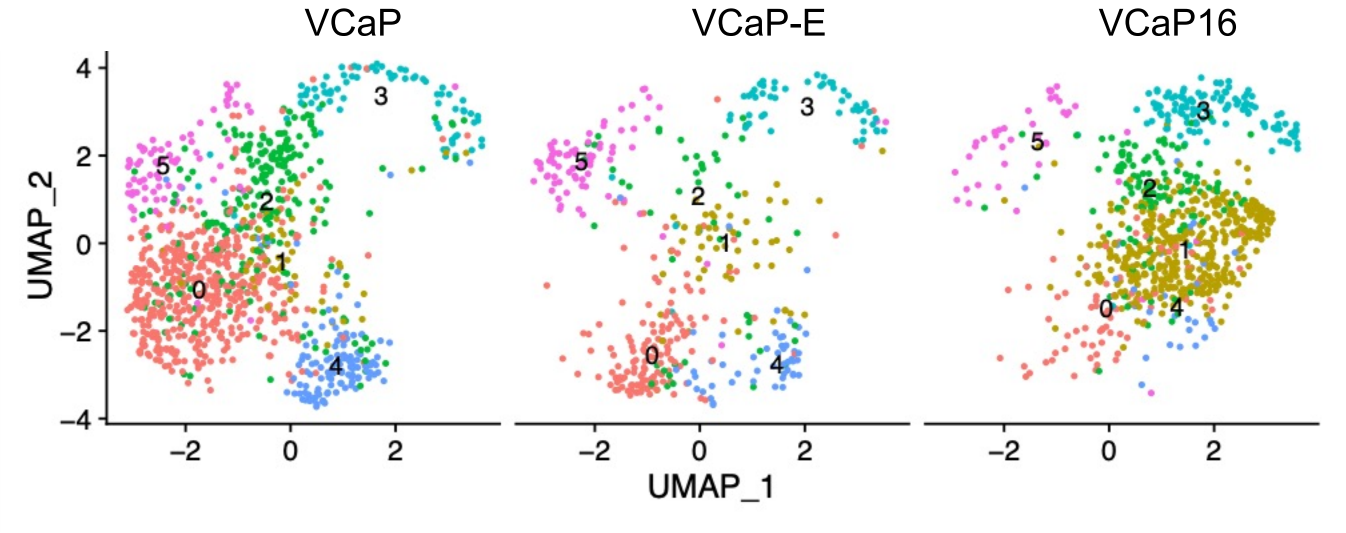


**B**

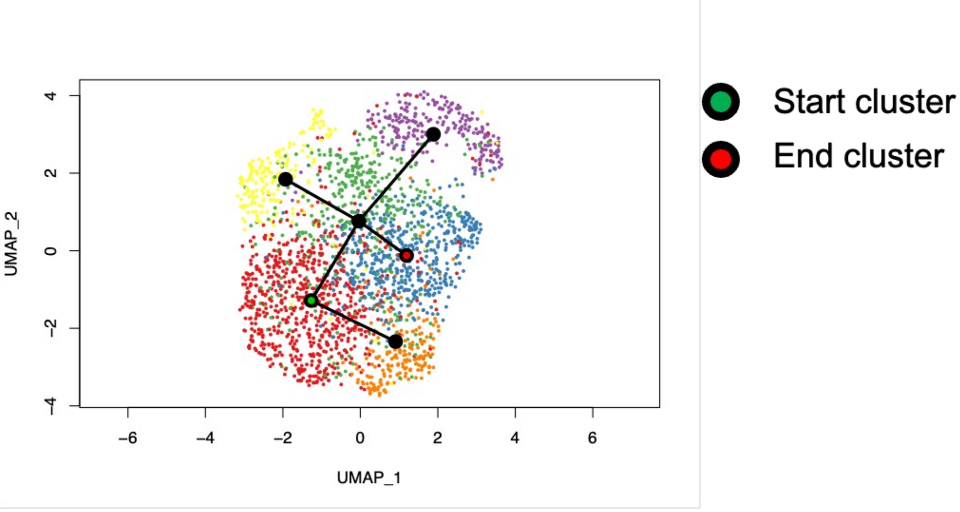

**Supplementary Figure S14. Subpopulations of cells that drive and maintain ENZ resistance. A)** Clustering on the combined single-cell ATAC-seq data (VCaP, ENZ-treated for 96 hrs VCaP (VCaP-E), and VCaP16) using Spectral Latent Manifold algorithm. Projection on UMAP plots. **B)** Pseudotime progression by trajectory inference analysis applied to the identified clusters in *A)*.

**Supp Fig S15**

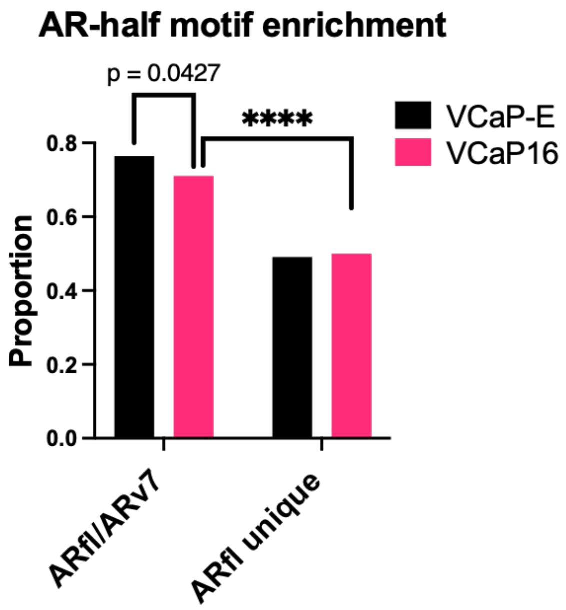
AR-half motif enrichment
p = 0.0427
****
VCaP-E
VCaP16
Proportion
0.8
0.6
0.4
0.2
0.0
ARfl/ARv7
ARfl unique

**Supplementary Figure S15. AR-half motif enrichment.** AR-half motif enrichment by HOMER at AR binding sites with canonical AREs versus those with non-consensus AREs (others) in VCaP cells after short term ENZ treatment (VCaP-E) versus in VCaP16 cells.

### Supp Fig S16

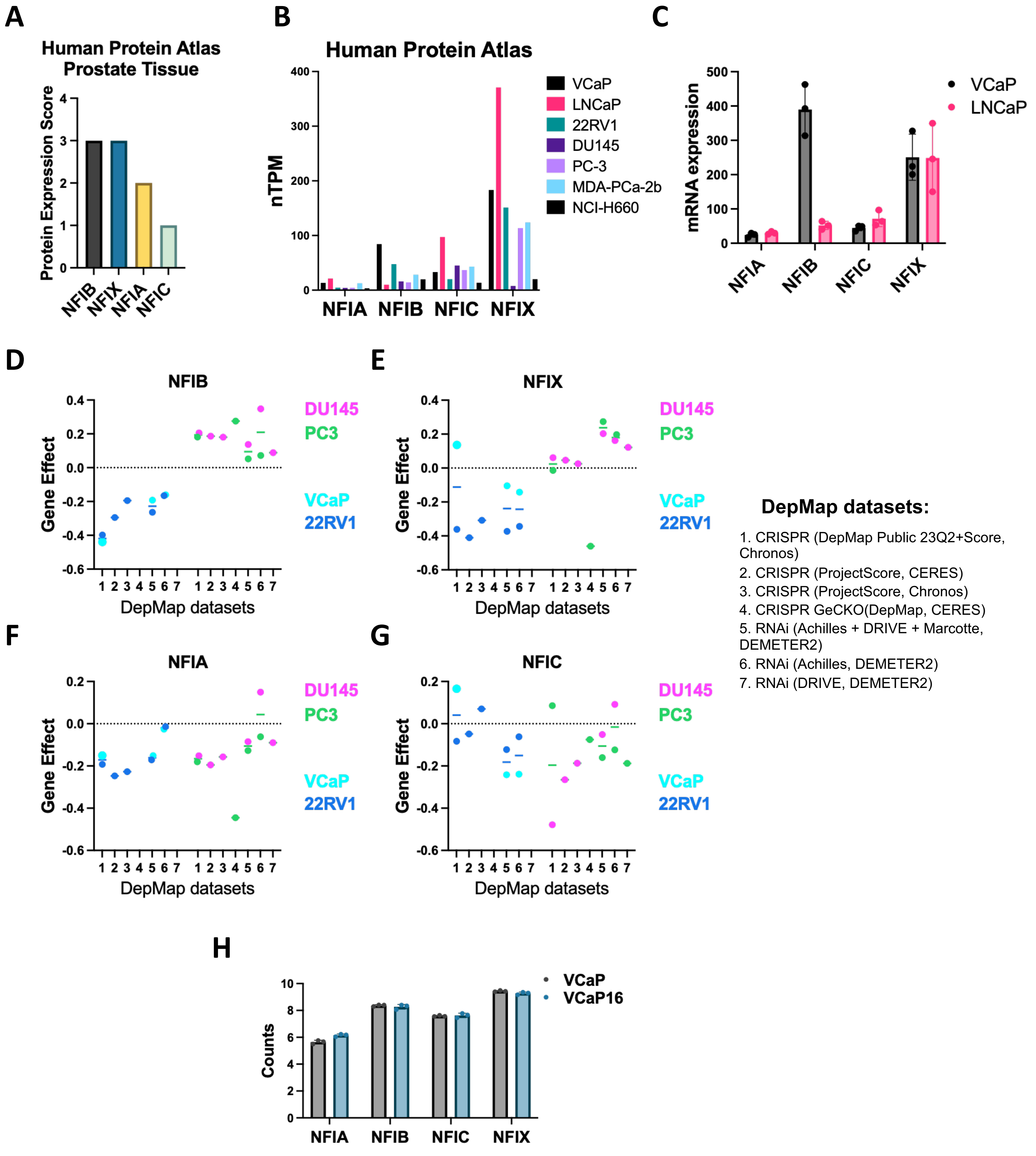
A
Human Protein Atlas
Prostate Tissue
Protein Expression Score
NFIB
NFIX
NFIA
NFIC
B
Human Protein Atlas
nTPM
VCaP
LNCaP
22RV1
DU145
PC-3
MDA-PCa-2b
NCI-H660
NFIA NFIB NFIC NFIX
C
mRNA expression
VCaP
LNCaP
NFIA
NFIB
NFIC
NFIX
D
NFIB
E
NFIX
F
NFIA
G
NFIC
Gene Effect
DepMap datasets
DU145
PC3
VCaP
22RV1
DepMap datasets:
1. CRISPR (DepMap Public 23Q2+Score, Chronos)
2. CRISPR (ProjectScore, CERES)
3. CRISPR (ProjectScore, Chronos)
4. CRISPR GeCKO(DepMap, CERES)
5. RNAi (Achilles + DRIVE + Marcotte, DEMETER2)
6. RNAi (Achilles, DEMETER2)
7. RNAi (DRIVE, DEMETER2)
H
Counts
VCaP
VCaP16
NFIA
NFIB
NFIC
NFIX

**Supplementary Figure S16. NFI expression levels in prostate cancer. A)** Expression of NFIA, NFIB, NFIC and NFIX proteins in prostate tissue (Human Protein Atlas). Protein expression scores: 0 – not detected; 1 – low; 2 - medium; 3 – high. **B)** Expression of NFIA, NFIB, NFIC and NFIX (nTPM) in prostate cancer cell lines (Human Protein Atlas). **C)** qRT-PCR determined expression of NFIA, NFIB, NFIC and NFIX in LNCaP and VCaP cell lines. **D-G)** NFIB, NFIX, NFIA and NFIC loss effects in CRISPR and RNAi screens in AR positive prostate cancer cells (VCaP and 22RV1) and AR negative prostate cancer cells (DU145 and PC3). Data was acquired from the DepMap portal. **H)** RNA-seq of RNA isolated from VCaP and VCaP16 cells showing expression of NFIA, NFIB, NFIC and NFIX (normalized counts) in VCaP and VCaP16 cells.

### Supp Fig S17

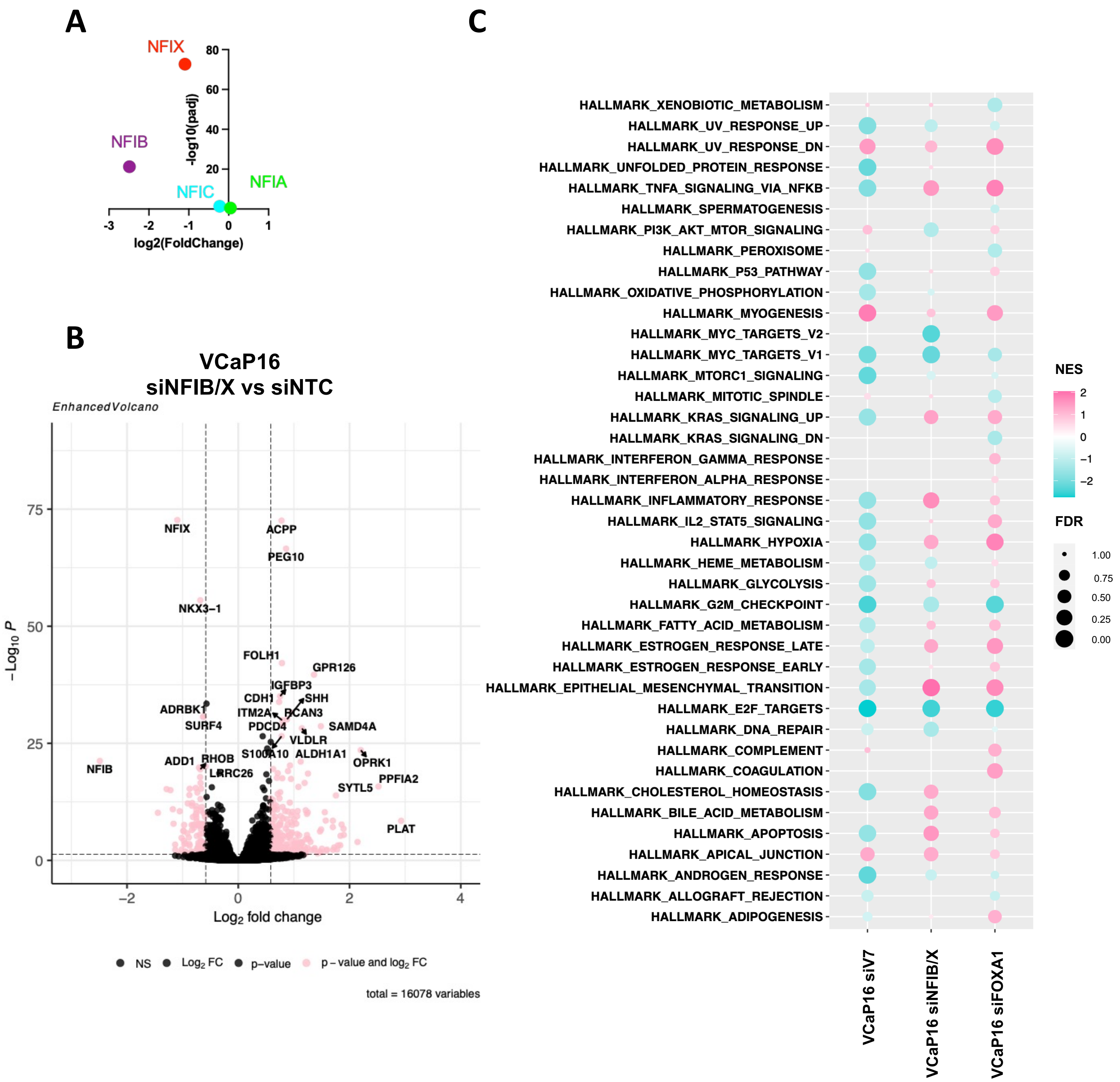
A
NFIX
NFIB
NFIC
NFIA
-log10(padj)
log2(FoldChange)
B
VCaP16
siNFIB/X vs siNTC
EnhancedVolcano
NFIX
ACPP
PEG10
NKX3−1
FOLH1
GPR126
IGFBP3
CDH1
SHH
ADRBK1
ITM2A
PCAN3
SURF4
PDCD4
SAMD4A
VLDLR
S100A10
ALDH1A1
ADD1
RHOB
OPRK1
NFIB
LRRC26
PPFIA2
SYTL5
PLAT
−Log10 P
Log2 fold change
NS
Log2 FC
p−value
p − value and log2 FC
total = 16078 variables
C
HALLMARK_XENOBIOTIC_METABOLISM
HALLMARK_UV_RESPONSE_UP
HALLMARK_UV_RESPONSE_DN
HALLMARK_UNFOLDED_PROTEIN_RESPONSE
HALLMARK_TNFA_SIGNALING_VIA_NFKB
HALLMARK_SPERMATOGENESIS
HALLMARK_PI3K_AKT_MTOR_SIGNALING
HALLMARK_PEROXISOME
HALLMARK_P53_PATHWAY
HALLMARK_OXIDATIVE_PHOSPHORYLATION
HALLMARK_MYOGENESIS
HALLMARK_MYC_TARGETS_V2
HALLMARK_MYC_TARGETS_V1
HALLMARK_MTORC1_SIGNALING
HALLMARK_MITOTIC_SPINDLE
HALLMARK_KRAS_SIGNALING_UP
HALLMARK_KRAS_SIGNALING_DN
HALLMARK_INTERFERON_GAMMA_RESPONSE
HALLMARK_INTERFERON_ALPHA_RESPONSE
HALLMARK_INFLAMMATORY_RESPONSE
HALLMARK_IL2_STAT5_SIGNALING
HALLMARK_HYPOXIA
HALLMARK_HEME_METABOLISM
HALLMARK_GLYCOLYSIS
HALLMARK_G2M_CHECKPOINT
HALLMARK_FATTY_ACID_METABOLISM
HALLMARK_ESTROGEN_RESPONSE_LATE
HALLMARK_ESTROGEN_RESPONSE_EARLY
HALLMARK_EPITHELIAL_MESENCHYMAL_TRANSITION
HALLMARK_E2F_TARGETS
HALLMARK_DNA_REPAIR
HALLMARK_COMPLEMENT
HALLMARK_COAGULATION
HALLMARK_CHOLESTEROL_HOMEOSTASIS
HALLMARK_BILE_ACID_METABOLISM
HALLMARK_APOPTOSIS
HALLMARK_APICAL_JUNCTION
HALLMARK_ANDROGEN_RESPONSE
HALLMARK_ALLOGRAFT_REJECTION
HALLMARK_ADIPOGENESIS
VCaP16 siV7
VCaP16 siNFIB/X
VCaP16 siFOXA1
NES
FDR
1.00
0.75
0.50
0.25
0.00

**Supplementary Figure S17. Transcriptomic effects following knockdown of ARv7, NFIB/X and FOXA1 in VCaP16. A)** Effects of combined siNFIB and NFIX on NFI transcripts confirming selective depletion of NFIB and NFIX. **B)** Volcano plot of differentially expressed genes in RNA-seq data from VCaP16 with siRNA knockdown of NFIB and NFIX (siNFIB/X) versus non-targeting control (siNTC). **C)** Gene Set Enrichment Analysis (GSEA) of RNA-seq data from VCaP16 cells with siRNA knockdown of ARv7 (left), with siRNA knockdown of NFIB and NFIX (middle), and with siRNA knockdown of FOXA1 (right).

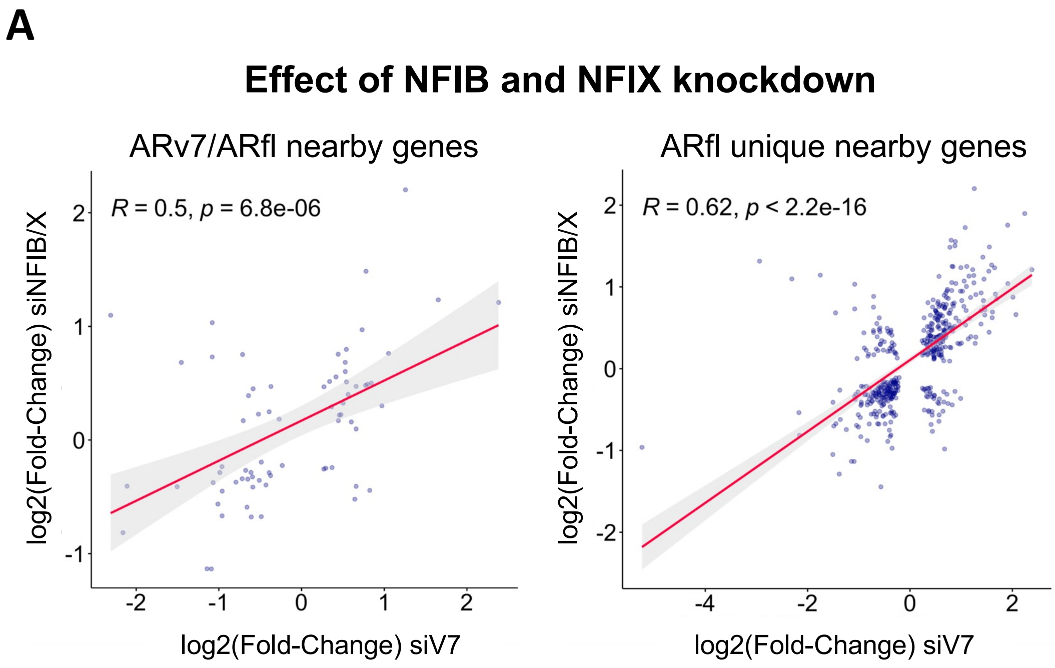
A
Effect of NFIB and NFIX knockdown
ARv7/ARfl nearby genes
R = 0.5, p = 6.8e-06
log2(Fold-Change) siNFIB/X
log2(Fold-Change) siV7
ARfl unique nearby genes
R = 0.62, p < 2.2e-16
log2(Fold-Change) siNFIB/X
log2(Fold-Change) siV7

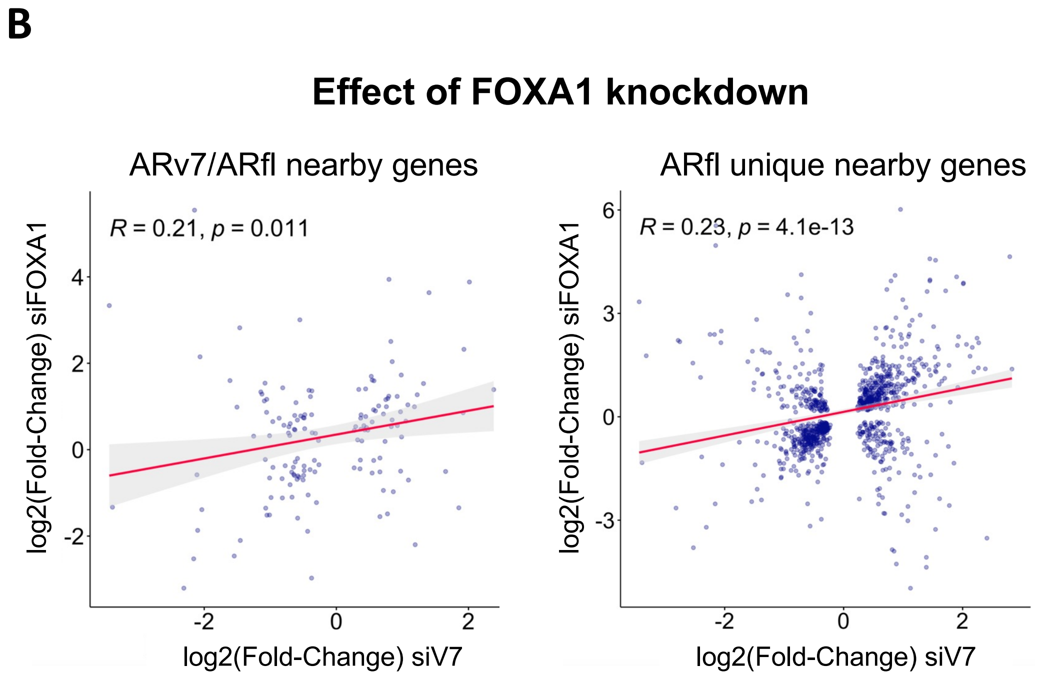
B
Effect of FOXA1 knockdown
ARv7/ARfl nearby genes
R = 0.21, p = 0.011
log2(Fold-Change) siFOXA1
log2(Fold-Change) siV7
ARfl unique nearby genes
R = 0.23, p = 4.1e-13
log2(Fold-Change) siFOXA1
log2(Fold-Change) siV7

**Supplementary Figure S18. Correlation between effects of ARv7, NFIB/X and FOXA1 knockdown at the ARv7 and ARfl unique binding site linked genes. A)** Pearson's correlation of log2(Fold-Change) values of significantly differentially expressed genes (padj < 0.1) altered with siRNA knockdown of ARv7 (siV7) versus siRNA knockdown of NFIB and NFIX (siNFIB/X) and **B)** siV7 versus siRNA knockdown of FOXA1 (siFOXA1) at the ARv7 and ARfl unique binding site linked genes in VCaP16 cells.

Figure S19

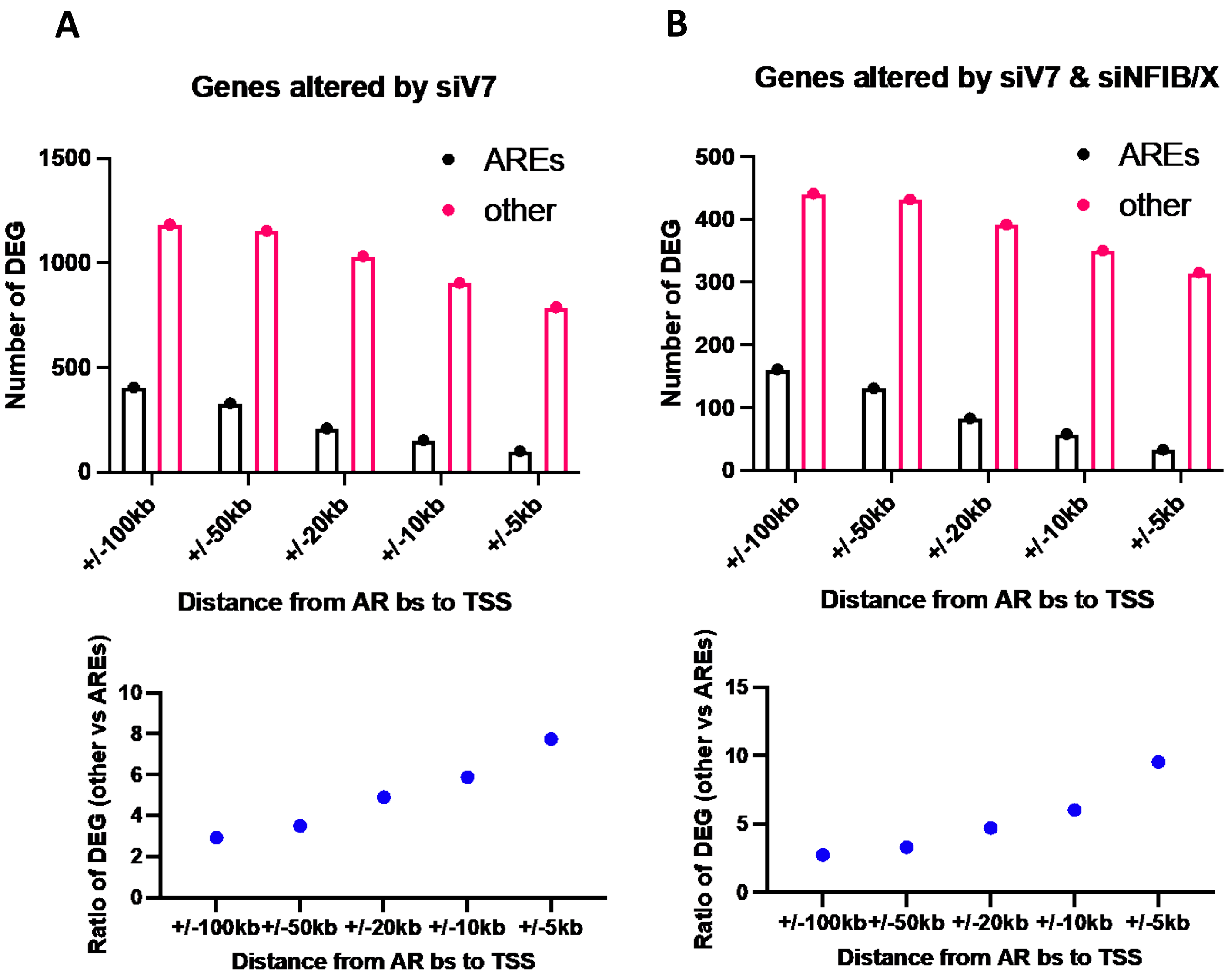
A
Genes altered by siV7
AREs
other
Number of DEG
1500
1000
500
0
+/-100kb
+/-50kb
+/-20kb
+/-10kb
+/-5kb
Distance from AR bs to TSS
Ratio of DEG (other vs AREs)
10
8
6
4
2
0
Distance from AR bs to TSS
B
Genes altered by siV7 & siNFIB/X
AREs
other
Number of DEG
500
400
300
200
100
0
+/-100kb
+/-50kb
+/-20kb
+/-10kb
+/-5kb
Distance from AR bs to TSS
Ratio of DEG (other vs AREs)
15
10
5
0
Distance from AR bs to TSS

**Supplementary Figure S19. Differentially expressed genes altered by depletion of ARv7 and NFIB/X linked to AR binding sites with canonical or non-canonical AREs. A)** Differentially expressed genes (DEG) altered with siRNA knockdown of ARv7 (siV7) (padj < 0.05) linked to ARfl binding sites identified in VCaP16 cells within +/-100kb, +/-50kb, +/-20kb, +/-10kb, +/-5kb with canonical or non-canonical AREs (above); and the ratios of DEG linked to ARfl finding sites with non-canonical AREs (others) versus canonical AREs (AREs) (below). **B)** DEG altered with both siRNA knockdown of ARv7 (siV7) and siRNA knockdown of NFIB and NFIX (siNFIB/X) (padj < 0.1) linked to ARfl binding sites identified in VCaP16 cells within +/-100kb, +/-50kb, +/-20kb, +/-10kb, +/-5kb with canonical or non-canonical AREs (above); and the ratios of DEG linked to ARfl finding sites with non-canonical AREs (others) versus canonical AREs (AREs) (below).

Figure S20

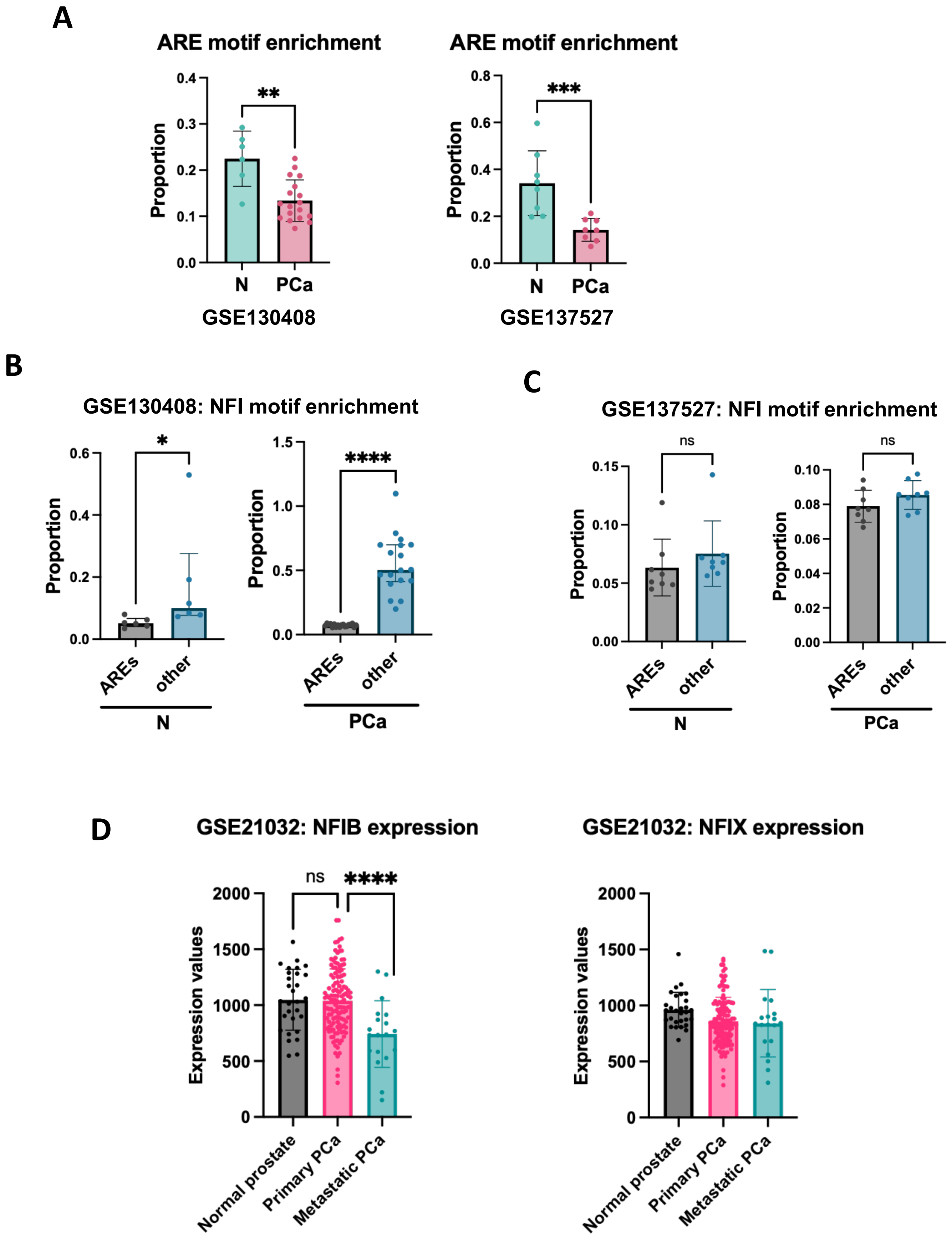
A
ARE motif enrichment
Proportion
**
N
PCa
GSE130408
ARE motif enrichment
Proportion
***
N
PCa
GSE137527
B
GSE130408: NFI motif enrichment
Proportion
*
****
AREs
other
N
PCa
C
GSE137527: NFI motif enrichment
Proportion
ns
ns
AREs
other
N
PCa
D
GSE21032: NFIB expression
Expression values
ns
****
Normal prostate
Primary PCa
Metastatic PCa
GSE21032: NFIX expression
Expression values
Normal prostate
Primary PCa
Metastatic PCa

**Supplementary Figure S20. NFI motif enrichment at non-canonical AREs in prostate cancer versus normal prostate. A)** ARE motif enrichment by HOMER at AR binding sites in normal prostate tissue (N) versus primary prostate cancer (PC) in two clinical datasets GSE139498 and GSE137527 (**$p<0.01$, ***$p<0.001$, Mann-Whitney test). **B - C)** NFI motif enrichment at canonical or non-canonical AREs at AR binding sites in normal prostate tissue versus prostate cancer. Data from two studies. **D)** NFIB and NFIX mRNA expression in normal prostate, primary, and metastatic prostate cancer.
